# Prediction and design of transcriptional repressor domains with large-scale mutational scans and deep learning

**DOI:** 10.1101/2024.09.21.614253

**Authors:** Raeline Valbuena, AkshatKumar Nigam, Josh Tycko, Peter Suzuki, Kaitlyn Spees, Aradhana, Sophia Arana, Peter Du, Roshni A. Patel, Lacramiora Bintu, Anshul Kundaje, Michael C. Bassik

**Author notes:** These authors contributed equally to this work (Raeline Valbuena, AkshatKumar Nigam).

## Abstract

Regulatory proteins have evolved diverse repressor domains (RDs) to enable precise context-specific repression of transcription. However, our understanding of how sequence variation impacts the functional activity of RDs is limited. To address this gap, we generated a high-throughput mutational scanning dataset measuring the repressor activity of 115,000 variant sequences spanning more than 50 RDs in human cells. We identified thousands of clinical variants with loss or gain of repressor function, including TWIST1 HLH variants associated with Saethre-Chotzen syndrome and MECP2 domain variants associated with Rett syndrome. We also leveraged these data to annotate short linear interacting motifs (SLiMs) that are critical for repression in disordered RDs. Then, we designed a deep learning model called TENet (<u>T</u>ranscriptional <u>E</u>ffector <u>Net</u>work) that integrates sequence, structure and biochemical representations of sequence variants to accurately predict repressor activity. We systematically tested generalization within and across domains with varying homology using the mutational scanning dataset. Finally, we employed TENet within a directed evolution sequence editing framework to tune the activity of both structured and disordered RDs and experimentally test thousands of designs. Our work highlights critical considerations for future dataset design and model training strategies to improve functional variant prioritization and precision design of synthetic regulatory proteins.

## Introduction

Transcriptional effector domains (EDs) are functional protein modules that confer gene activating or silencing potential. Typically found in transcription factors (TFs) and chromatin remodelers (CRs), they enable dynamic control of the transcriptome by recruiting co-factors, interacting with the basal transcription machinery, and modifying chromatin state ^1–3^. The structural and functional diversity of EDs has facilitated the evolution of complex gene regulatory programs defining numerous cellular functions, states, and fates (**Fig.1a**) ^4–6^. Especially in eukaryotes, the diversification of effectors with repressor activity has greatly expanded the number of mechanisms and potencies of silencing ^7,8^. Mutations that cause even subtle misregulation of repression can drive pathogenic outcomes ^7,9,10^. Therefore, deciphering the functional features and consequences of sequence variation in repressor domains is crucial to improving our understanding of the genetic basis of disease and enabling the design of more precise genetic tools for therapeutic intervention.

Experiments characterizing effector strength typically involve synthetic fusion of an ED to a DNA-binding domain (DBD), recruitment to a reporter gene, and measurement of subsequent changes to gene expression. Though human EDs are often regulated via post-translational modifications ^11^ – which may be highly species specific – many of the seminal experiments quantifying effector function were conducted in non-mammalian systems ^12–18^. Recently, pooled screening approaches were developed to simultaneously measure the effector activity of thousands of EDs in human cells (**Fig.1b**) ^19,20^. These assays have since been leveraged to identify hundreds of previously unannotated human domains with effector activity ^21–23^, which had been otherwise difficult to detect without experimental validation in a biologically relevant context.

However, scaling experiments to exhaustively probe the space of all possible EDs and variants is unfeasible. A promising solution is to develop predictive computational models that can accurately map sequence to function using sufficiently large datasets and leverage them to compute predictions for unmeasured sequences ^24^. These models could also guide the directed engineering of existing EDs or the *de novo* design of synthetic EDs with desired activities, as has been previously attempted for other protein classes ^25–30^.

Neural networks have emerged as a powerful class of computational models to learn informative representations of protein sequences, infer protein structure and function, and prioritize potentially pathogenic variants ^31–44^. Datasets generated from deep mutational scanning (DMS) experiments have proven valuable for training and evaluating sequence to function models ^43,45^. However, predictive sequence models of ED activity have been limited, and have primarily focused on activation domains (ADs) ^6,17,46^. ADs tend to be “acid blobs and negative noodles” with compositional preferences for charged and hydrophobic residues ^7,8,21,22,47,48^. In contrast, repressor domains (RDs) are generally less conserved than ADs, both in sequence and structure, and are hence more challenging to model ^7,8,21,22,47,48^. The unifying sequence and structural preferences of RDs remain poorly characterized, presenting significant challenges for estimating variant effects on RD activity and custom design of synthetic RDs. Optimal strategies for designing training datasets and models that would promote robust generalization across RDs with diverse sequences and structures are underexplored.

To address these challenges, we generated one of the largest DMS datasets reporting functional activity measurements of transcriptional repressor domains and variants in human cells, characterizing over 115,000 missense and multi-substituted coding variants across more than 50 RDs. We analyzed how variant-induced changes in repression correlated with pathogenicity and designed a second screen to characterize <u>v</u>ariants of <u>u</u>nknown <u>s</u>ignificance (VUS) and pathogenic variants in 189 additional RDs. These data collectively identified 1264 clinically observed variants with significantly altered repressor function, including TWIST1 HLH variants associated with Saethre-Chotzen syndrome and MECP2 domain variants associated with Rett syndrome. Using these DMS data, we also identified functional short linear interacting motifs (SLiMs) critical for repression in disordered domains. We developed one of the first deep learning models to accurately predict repressor activity for both structured and unstructured domains by integrating sequence, structure and biochemical representations of sequence variants. The model not only accurately predicts the activity of unseen sequence variants of RDs in the training set but also generalizes to new domains with performance decaying as a function of their homology to RDs in the training set. Remarkably, scaffolding a new domain with even a single functional measurement (one-shot context) allowed the model to accurately generalize to diverse variants of that domain. We used the model within a directed evolution sequence editing framework to optimize the activity of both structured and disordered RDs and experimentally tested thousands of designs with diverse activities. Altogether, this work demonstrates a strategy to design large-scale functional measurements of protein variants to power deep learning models capable of predicting and designing repressor activity of diverse RDs.

## Results

### Deep mutational scan of diverse repressor domains (Fig. 1)

To power a deep learning model to quantitatively predict repressor activity from AA sequence, we generated AA-resolution data measuring the repressive strength of over 115,000 sequence variants of 54 repressor domains using a high-throughput recruitmentment assay (HT-recruit) ^19,21–23,49,50^ (**Fig. 1a-b**). When selecting domains for the <u>R</u>epressor <u>D</u>omain <u>D</u>eep <u>M</u>utational <u>S</u>canning (RD-DMS) library, we sampled RDs representing various 1) protein families, 2) known repressive strengths and/or mechanisms, and 3) degrees of structure and disorder (**Fig. 1c, Supplementary Fig. 1a, Supplementary Table 1**). First, we selected RDs that have been functionally annotated in Pfam and InterPro ^51,52^; ^53,54,52^. These include domains from large and pervasive classes, such as the Zinc Finger (ZF) and Helix-loop-helix (HLH) families, and from core machinery involved in mediating heterochromatin, like the CBX and Polycomb proteins ^55,56^. Additionally, we selected 15 previously unannotated domains that had exhibited strong repressive strengths in HT-recruit screens ^19,21^. We classified each RD as structured, hybrid, or disordered based on computational prediction (**Methods: Domain classification and disorder prediction**) ^39^. Importantly, although all previously unannotated domains were selected on the basis of repressive strength – without considering structure – the majority (12/15) were classified as disordered. 19 RDs were disordered in total, with the highest representation coming from the unannotated class **(Supplementary Fig. 1a**). This is consistent with the observation that it has been difficult to functionally annotate disordered domains, especially those lacking sequence homology, without experimental discovery.

**Figure 1:**
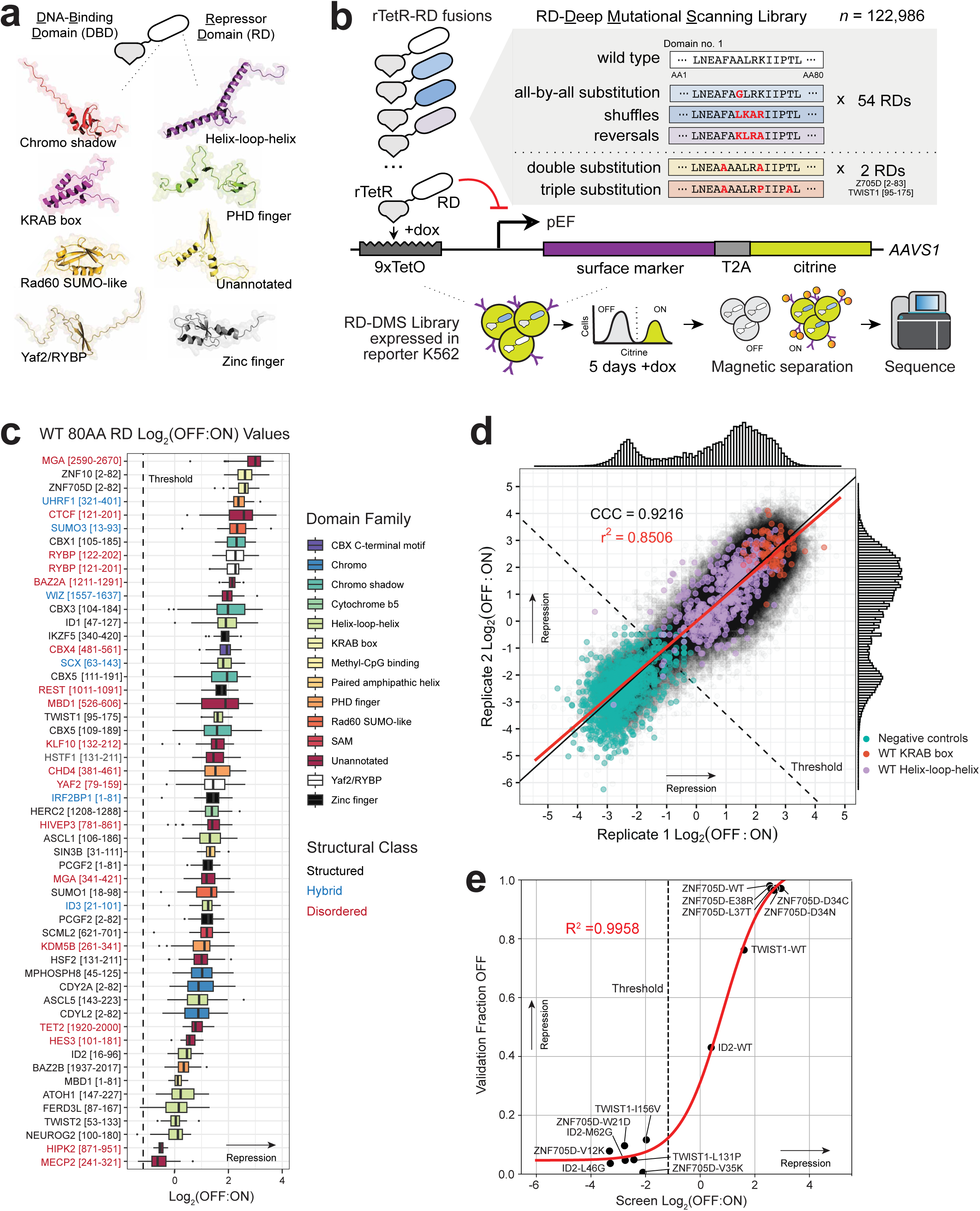
Generating a large-scale dataset of AA-resolution measurements of repressor strength by high-throughput mutational scans of 54 diverse repressor domains. a. <u>R</u>epressor <u>d</u>omains (RD) are structurally diverse and are identified by their ability to drive transcriptional silencing when recruited to genetic loci by <u>D</u>NA <u>b</u>inding <u>d</u>omains (DBD). Schematic showing structural predictions for a representative 80AA domain for each indicated repressor family, as generated by AlphaFold 2. b. Schematic illustrating the <u>r</u>epressor <u>d</u>omain <u>d</u>eep <u>m</u>utational <u>s</u>canning (RD-DMS) library composition and HT-recruit to measure repressor activity. An oligo library is synthesized encoding various RDs and variants. The domains are cloned as fusions to rTetR and are subsequently transduced into K562-reporter cells in a pooled fashion, such that the vast majority of cells receive one library element. RD-mediated silencing of the reporter post doxycycline (dox) enabled recruitment is monitored by citrine fluorescence, and a synthetic surface marker enables magnetic bead separation of cells with “OFF” or “ON” reporters. These populations are sequenced to determine element enrichment. c. Log_2_(OFF:ON) ratios for all WT codon variants (50 per RD), ranked from highest (top) to lowest (bottom) mean repressor activity. The data are colored by domain family and the y-axis data labels are colored to indicate structural class (**Methods: Domain classification and disorder prediction**). Each box extends from the 25th to 75th quartiles; within each box, vertical lines denote median values; lines outside the box extend to minimum and maximum values except in cases where these values are denoted by dots (potential outliers). d. Reproducibility from two biological replicates of the RD-DMS screen. The concordance correlation coefficient (CCC = 0.9216) is relative to the line with slope = 1 and y-intercept = 0 as shown in black. The coefficient of determination (r^2^ = 0.8506) is derived from the best fit linear regression as shown in red. The repressor threshold (dashed line) is set two standard deviations above the mean (**x̄ + 2σ**) of the negative controls, indicated in turquoise. All WT measurements for RDs belonging to the KRAB box (Pfam ID: PF01352) and (b)HLH (Pfam ID: PF00010) repressor families are indicated in red and purple, respectively. e. Comparison between screen Log_2_(OFF:ON) values and the proportion of OFF cells measured in individual screen validations. A logistic model fit is drawn as a solid red line (R^2^ = 0.9958, n = 14).

We systematically designed DMS variants for each selected RD to assess features known and hypothesized to be important for repressor function. First, we generated exhaustive single substitutions where each wild-type AA was swapped for every other AA (**Fig. 1b**). This mutational schema assesses the essentiality of each residue and can reveal key biochemical properties, such as charge and hydrophobicity, necessary for function ^57^. We then generated mutants to assess features known to be functionally important in disordered domains; here, residues within windows of 4 or 8 AAs were either randomly shuffled or reversed (**Fig. 1b, Methods: RD-DMS library design**). Primarily, these mutants should help identify <u>S</u>hort <u>Li</u>near <u>M</u>otifs (SLiMs) in disordered domains, where the relative positions of invariant residues within a stretch of 3-10AAs are important for maintaining function ^58,59^. These windowed-shuffle mutants were additionally motivated by observations that AA composition permits the formation of functional microdomains or creates hydrophobic patterns that sometimes enable activator activity, but have yet to be clearly articulated in RDs ^12,15,16,18^. Altogether, these mutants probe functionally relevant properties in both structured and disordered RDs.

To capture potential epistatic, additive, or multiplicative effects between intra-domain AA interactions, we also generated all possible alanine double mutants and 16,000 triple mutants within the 80AA tiles of the ZNF705D Krüppel-associated box (KRAB, Pfam ID: PF01352) and the TWIST1 basic helix-loop-helix (bHLH, Pfam ID: PF00010) domains ^60,61^, **Methods: RD-DMS library design**). KRAB domains have been widely adopted for synthetic biology and HLH domains are a large and highly modular family ^29,30,62, 63–65,29,30,62^. Finally, we included 3,546 scramble/negative controls (**Methods: RD-DMS library design**). In all, the total size of the RD-DMS library was 122,986 elements (**Fig. 1b, Supplementary Table 2**).

This library was cloned into a lentiviral vector where each element was expressed as a fusion to rTetR and subsequently transduced into K562 reporter cells (**Supplementary Table 3**); the reporter locus contains a 9xTetO array upstream a constitutively active pEF promoter driving expression of citrine and a surface marker (**Fig. 1b**) ^19^. Doxycycline (Dox) was then added to induce recruitment (**Supplementary Fig. 1b, Methods: Pooled delivery of libraries to K562 reporter cells via lentiviral transduction**). After 5 days of recruitment, the resulting ON and OFF populations were sorted by magnetic separation (**Supplementary Fig. 1c, Methods: Magnetic separation**). The RDs in each population were identified by NGS and Log_2_(OFF:ON) enrichments were calculated for each library element as a measure of repressor strength (**Supplementary Fig. 1d**). The screen was repeated in biological duplicate and the correlation between replicates was strong (*r* = 0.9223, Concordance correlation coefficient (CCC) = 0.9216) (**Fig. 1d**). Less than 5% dropout of library elements was observed with 95.06% of the total library retained across both replicates (94.37% in BioRep1, 94.06% in BioRep2) (**Supplementary Table 4**). In all, our screen retained high diversity and representation of all library elements (**Supplementary Fig. 1e**).

We calculated a threshold (Log_2_(OFF:ON) = −1.173) for calling mutants with repressor activity on the basis of the distribution of activity of the negative controls (**Methods: Computing enrichments and hit thresholds**). Activity of 94.08% of all negative controls fell under the repressor threshold and 98.60% of all positive controls exceeded the threshold (**Supplementary Fig. 1f**). The WT activity for the ATF7IP [962-1042] domain did not exceed the repressor threshold in our screen, so this domain and its associated data were not considered in subsequent discussion. To test the reproducibility of mutations in variable sequence contexts, we compared the effects of homologous mutants in 3 cases where the selected 80AA tiles contained overlap (RDs for CBX5, PCGF2, and RYBP). Although the overall sequence context of these mutants varied slightly, the effects of the same point mutation show high reproducibility in all cases (**Supplementary Fig. 1g**).

We performed individual validations for a subset of domains and mutants by recloning them, stably integrating them in the K562-reporter line, and measuring their repression of citrine after dox-induced recruitment by flow cytometry (**Methods**). For the ZNF705D KRAB domain, we tested a series of mutants that showed clear phenotypes in the screen and were expected to either break or retain function based on previous studies of KRAB domains. V12K should disrupt required hydrophobic interactions with the co-repressor KAP1, W21D likely disrupts silencing independent of KAP1 binding, and V35K destroys the functionally required VMLEN motif ^19,66,67^. D34C, D34N, L37T and E38R were previously characterized as non-perturbative at homologous sites in the ZNF10 KRAB domain ^19^. Indeed, our validations agree with the measured and hypothesized activities of all these mutants (**Supplementary Fig. 1h**, i). Additional individual validations performed with selected mutants from HLH domains also recapitulated screen results quantitatively in this low-throughput assay (**Supplementary Fig. 1j, 1k**). Western blots for FLAG confirm that loss of function is likely not due to loss of protein stability in these cases (**Supplementary Fig. 1l**). Overall, the fraction of cells where citrine is repressed at day 5 of dox-treatment in these validations was highly concordant with screen Log_2_(OFF:ON) values (R^2^ = 0.9958, n = 14) (**Fig. 1e**). These results demonstrate that the RD-DMS HT-recruit screen delivered reproducible and quantitative data for a diverse set of repressor domains and robustly identifies mutants that suppress or augment repressor activity.

### Characterization of clinically-relevant missense variants (Fig. 2)

Because loss of function (LoF) or gain of function (GoF) by a repressor could result in dysregulated gene expression and pathogenic outcomes, identification of these variants is of considerable clinical interest ^68,69^.

**Figure 2:**
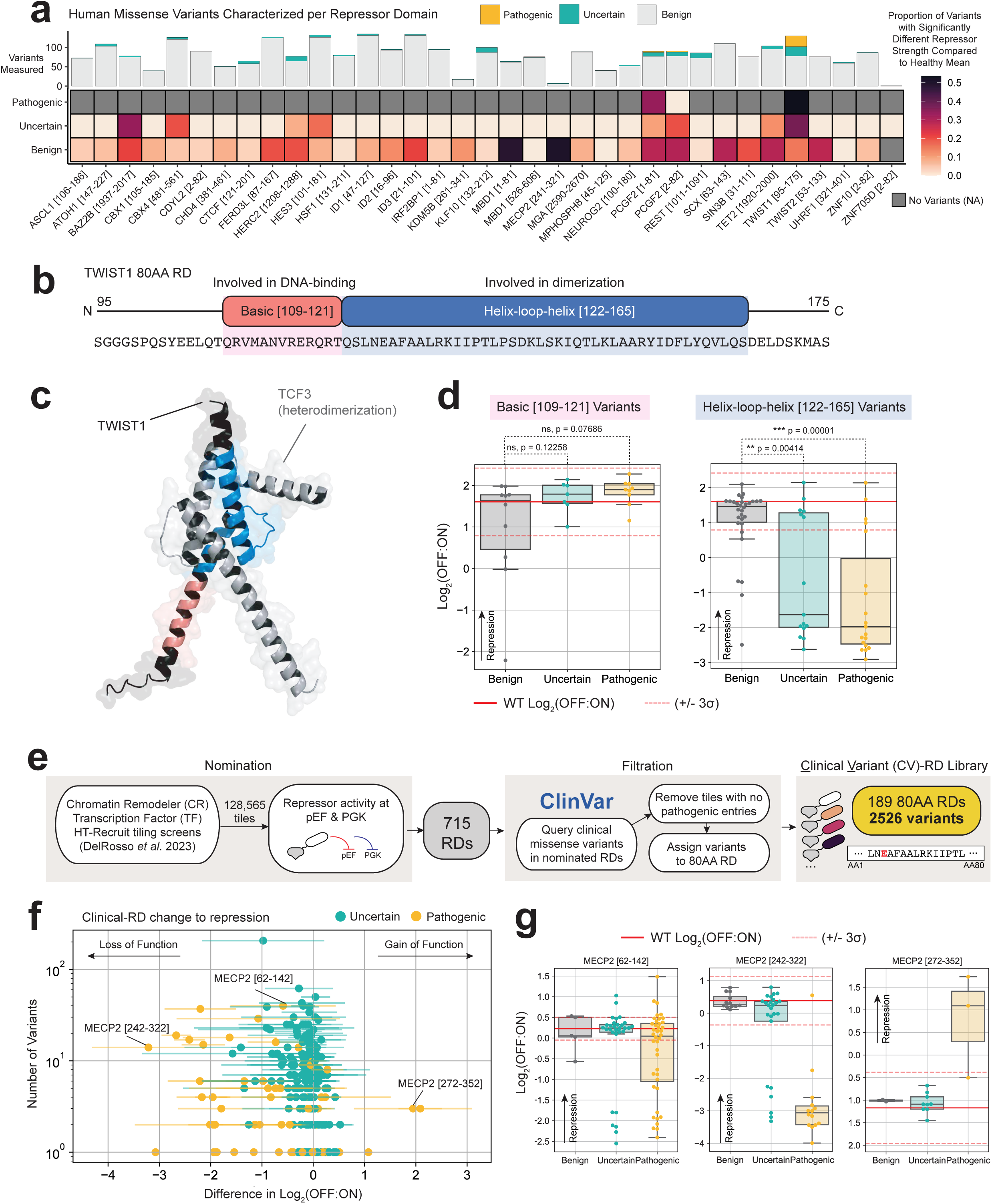
Functional characterization of clinically-relevant missense variants. a. Heatmap summarizing the proportion of pathogenic variants and variants of uncertain significance where the measured Log_2_(OFF:ON) value is greater than three standard deviations from the benign mean (**x̄ + 3σ**) per repressor domain. b. Annotated subdomains along the (b)HLH domain (Pfam ID: PF00010) of TWIST1 [95-175]. The N-terminal end of the RD [109-121] contains a basic domain (red) known to be involved in DNA-binding in an endogenous context. The helix-loop-helix (HLH) domain [122-165] (blue) is a known be involved in hetero- and homodimerization with other (b)HLH domains. c. The AlphaFold-Multimer predicted complex of the TWIST1 bHLH RD interacting with the bHLH domain of known cofactor TCF3, forming an HLH heterodimer. The basic (red) and HLH subdomains (blue) of TWIST1 are highlighted by their respective colors in the structure. d. Log_2_(OFF:ON) values for human variants of the basic (left) and HLH (right) subdomains of the TWIST1 bHLH RD. Variants are classified by reported clinical significance. Differences in repressive strength between all benign variants (gray) and either all uncertain (turquoise) or pathogenic (yellow) variants were tested by Welch’s t-test; p-value significances are reported (ns = not significant, * < 0.05, ** < 0.01, *** <0.001). Individual data points are plotted as dots; boxplots are drawn as described previously (Fig. 1c), though no outliers are indicated. Red horizontal lines show the average Log2(OFF:ON) values for all WT encodings (solid) and +/- three standard deviations (dashed) (**x̄ + 3σ**). e. Schematic illustrating the <u>c</u>linical <u>v</u>ariant <u>r</u>epressor <u>d</u>omain (CV-RD) library design. HT-recruit data from Del Rosso *et al*. 2023 was mined to identify domains with repressor activity in our system. We queried ClinVar to identify RDs that had known pathogenic variants, then encoded all variants from the filtered list of RDs into 80 AA sequences. The final library contained elements encoding 189 WT RDs and 2526 missense variants (**Methods: CV-RD library design**). f. Scatterplot summarizing the change to repressor activity observed in uncertain (turquoise) and pathogenic (yellow) variants tested in the CV-RD library. Each dot represents the average difference in Log_2_(OFF:ON) of the indicated variant type vs. the average WT value of the respective RD: along the x-axis, negative values indicate loss of function (LoF) while positive values suggest gain of function (GoF). Horizontal lines show +/- three standard deviations (**x̄ + 3σ**). The data for the pathogenic variants of three MECP2 RDs are labeled. g. Log_2_(OFF:ON) values for three 80AA MECP2 RDs and their respective variants. Variants are classified by reported clinical significance and WT values are plotted as previously described (**d**).

Unlike mutations resulting in an premature stop codon or a frameshift mutation, it is much harder to decipher missense variants with functional ramifications ^70^. Since we measured the repressor activities of nearly all possible missense variants for the selected RDs in our RD-DMS screen, we next investigated whether any variants with significantly altered function correlated with pathogenicity.

To filter our data for germline variants known to exist in the human population, we considered single mutants in our RD-DMS library that overlapped with missense variants in gnomAD 4.0 and/or ClinVar ^9,71^. All mutants were then classified according to clinical significance as “benign” variants (including potentially benign variants from healthy individuals in gnomAD), variants of “uncertain” significance (VUS), or “pathogenic” variants (**Methods: Clinical annotations**). We measured variants classified as benign in all 54 RDs, and 35 of 54 RDs overlapped with known human variants annotated as either pathogenic or uncertain (**Fig. 2a, Supplementary Fig. 2a**, b). Pathogenic variants were found in RDs from PCGF2 and TWIST1, though as expected, few of our designed mutants are currently categorized as pathogenic, likely because the majority of the RDs screened are from essential genes, where variants resulting in LoF or GoF typically undergo negative selection and are purged from the population ^71–75^. For tiles with uncertain and pathogenic variants, we then calculated the proportion of these variants that had significantly altered repressive strengths compared to their benign counterparts (**Methods: Functional annotation of clinical variants**). Clinically-relevant missense variants in BAZ2B, HEC2, HES3, PCGF2, TET2, and TWIST1 were detected with significant LoF (**Supplementary Table 5**). We report all measured repression strengths for these variants as a record of their functional annotations (**Supplementary Table 5**).

The TWIST1 bHLH domain had the highest number of pathogenic and uncertain variants (52) and was the domain where the highest proportion (0.46) of these clinically-relevant variants had significant LoF compared to variants annotated as benign (**Fig. 2a**). Belonging to the bHLH family of repressors, this domain contains both basic [109-121] and HLH [122-165] subdomains, which are involved in DNA-binding and dimerization, respectively ^63,76^ (**Fig. 2b, c**). Fewer human variants were observed in the basic domain (26) than the HLH domain (64), consistent with observations that DBDs are less amenable to mutation than effectors ^7^. Additionally, because we use rTetR to recruit RDs to our reporter, we synthetically override LoF mutations that may disrupt DNA binding, which could explain why the vast majority of TWIST1 missense mutations with detectable LoF in our screen fall within the HLH subdomain (87.12%) (**Extended Data**). Therefore, we subsetted our TWIST1 mutants into their respective subdomains for subsequent analysis and performed statistical tests on all basic and HLH variants grouped separately (**Methods: Functional annotation of clinical variants**). We found that VUS and pathogenic variants in the basic region of the TWIST1 domain do not have statistically significant LoF compared to benign variants in our data (*p* = 0.12258, *p* = 0.07686, respectively; Welch’s *t*-test). In contrast, both VUS and pathogenic variants in the helix-loop-helix domain of TWIST1 were significantly different from benign variants and correlated with LoF (*p* = 0.00414, *p* = 0.00001, respectively; Welch’s *t*-test) (**Fig. 2d**).

TWIST1 plays essential regulatory roles in development, and its aberrant activity is associated with various pathogenicities ^63,77,78^. For example, germline heterozygous LoF missense mutants of TWIST1 are known to cause diseases leading to craniosynostosis such as Saethre-Chotzen syndrome (SCS) and Sweeny-Cox syndrome ^64,65^. Our data identified known pathogenic variants associated with SCS that occur in highly conserved residues of the HLH subdomain, such as I156V and L131P, as LoF variants (**Extended Data**). Single validations of these two variants confirmed LoF in our reporter assay, and Western blotting did not show decreased expression relative to WT (**Supplementary Fig. 3a**, b). Importantly, we functionally classify 9 VUS with similar degrees of LoF as known pathogenic variants, and 15 with similar repressive strengths as benign variants (**Supplementary Table 5**).

Like other HLH family domains, TWIST1 is known to form both homodimers and heterodimers with other ubiquitously expressed bHLH proteins, most notably E12/E47 (both coded by *TCF3*) ^63^. The residues found to be particularly sensitive to perturbation in our screen lie along the helical planes of the known dimerization interface (**Extended Data**). Because endogenous TWIST1 is not expressed in K562, we modeled TWIST1 dimers with TCF3, the highest expressed partner and therefore a likely corepressor in our system (**Fig. 2c, Supplementary Fig. 3c**). Previous molecular dynamics (MD) studies performed using TWIST1 dimers found that mutations in the HLH region of TWIST1 led to wider solvent-accessible cleft between monomers and destabilized dimers ^76^. For pathogenic variants like I156V and L131P, we also see an increase in atomic space at the dimerization interface through crude measurements of atomic distances between predicted AF-Multimer complexes (**Supplementary Fig. 3d**). This suggests that LoF in TWIST1 HLH variants could be due to decreased stability of dimer complexes. Altogether, our data highlight the TWIST1 bHLH as an example where functional measurements identify LoF and flag potential pathogenicity, and can provide rationale for future molecular-level inquiries.

Inspired by our observations that VUS and pathogenic variants could correlate with measured LoF, we designed a second screen to comprehensively generate functional annotations for additional RDs with clinically relevant variants. In short, we used ref. ^21^ to obtain a list of all WT RDs previously identified with HT-recruit, then cross-referenced ClinVar for variants that occurred within these RDs to create the <u>C</u>linical <u>V</u>ariant <u>R</u>epressor <u>D</u>omain (CV-RD) library (**Methods: CV-RD library design**). We then cloned and measured the repressor activities of all benign, uncertain, and pathogenic missense variants for 189 additional RDs, as identified based on these criteria (**Fig. 2e**). Importantly, this screen identifies 1264 additional human variants with significantly altered repressor activity, and 28 RDs where VUS and/or pathogenic variants categorically differ from WT values (**Supplementary Fig. 4**, **Methods: Functional annotation of clinical variants**). For example, this work highlights MECP2 as another repressor of interest: multiple 80AA RDs from this protein were identified where VUS and/or pathogenic variants had significantly different repressor activity. MECP2-mediated misregulation caused by changes in DNA-binding affinity are known to contribute to diseases such as Rett syndrome and neonatal-onset encephalopathy ^79,80^. These results provide more functional evidence implying that MECP2-mediated misregulation resulting from changes to its RD function could also potentially drive disease-associated phenotypes. Of particular interest, we find that pathogenic variants of MECP2 [242-322] tend to exhibit LoF, whereas in MECP2 [272-352] they tend to have gain of function (GoF) (**Fig. 2f, g**). The possibility that different variants of this gene could cause opposing transcriptional phenotypes could be studied in future work to better understand disease phenotypes at the cellular level. We also report the measured strength and all functional annotations for variants measured in the CV-RD library (**Supplementary Table 5**).

### Key functional patterns in structured and disordered domains (Fig. 3)

For structured domains, single substitutions tend to decrease repressor activity when introduced in functionally important domain architectures (**Extended Data**). For example, in SIN3B [31-111], a paired amphipathic helix (PAH) repeat (Pfam ID: PF02671), our data show intolerance to mutation in residues occurring at a periodicity of about 3 AAs, consistent with patterning expected for α-helices ^81,82^ (**Fig. 3a, b**). These residues tend to occur at the intra-molecular interfaces between paired helices, in line with previous observations that they are critical for maintaining PAH structure by segregation of the hydrophobic and polar residues between helical interfaces ^83^. We also observe this periodic patterning in the helical subdomains of all HLH family members – these occur along the known dimerization interface when mapped to their respective predicted structures (**Extended Data**). The AAs at the dimerization interfaces of these domains are known to mediate binding affinities between monomers and are therefore important for determining the role of each HLH and the strength of their repressor activity ^84,85^. For structured domains, we also generally observe high intolerance to windowed-shuffle or reversal mutations (**Fig. 3c, Extended Data**). Additionally, the same residues that are sensitive to substitution tend to be highly conserved, another indicator that specific residue compositions maintain functionally important secondary structures (**Fig. 3d, Extended Data**).

**Figure 3:**
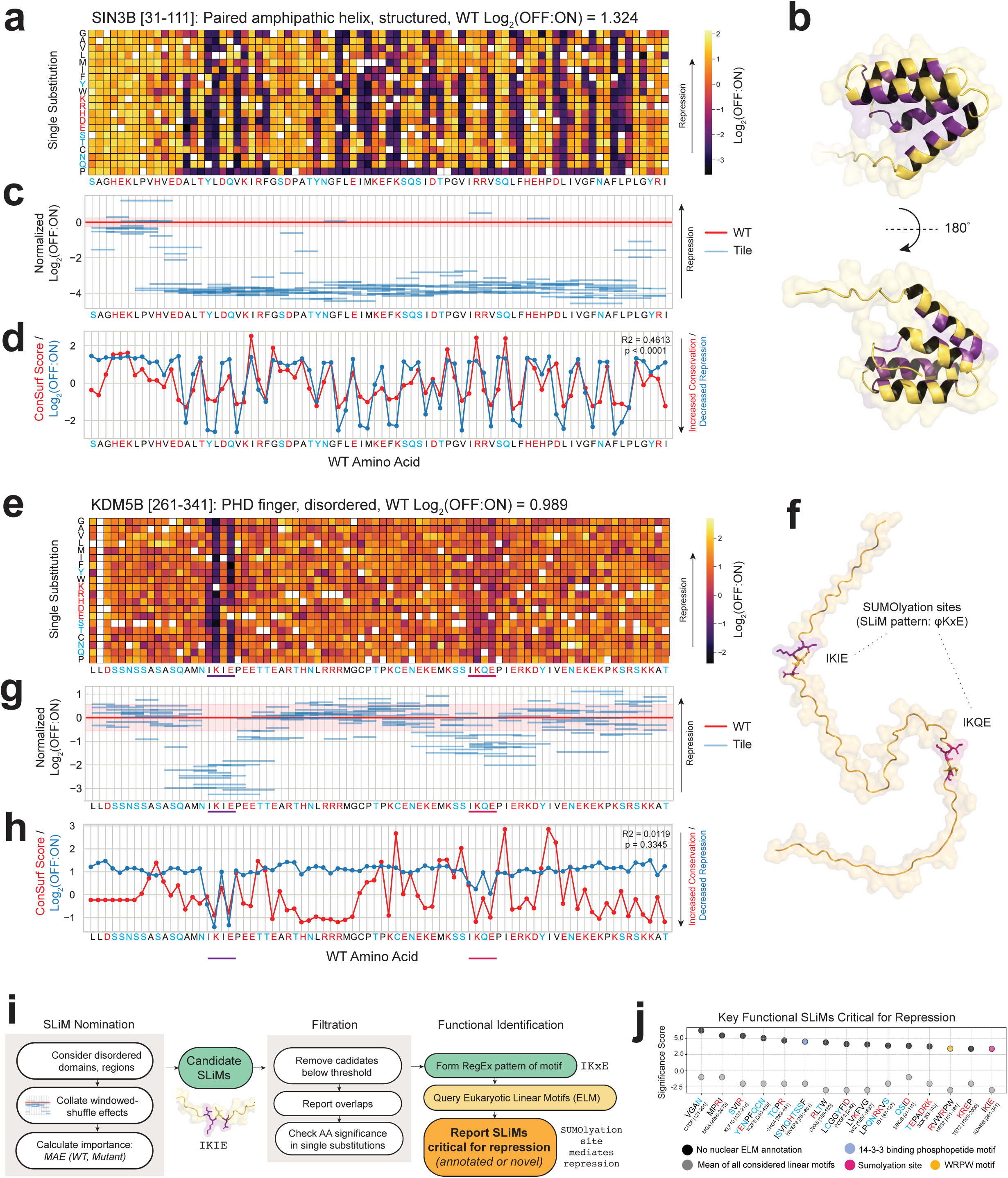
Structures and motifs important for repressor activity are identified with the DMS data. a. Heatmaps filled by the Log_2_(OFF:ON) values for each missense mutation along the 80 AA SIN3B RD. The wildtype sequence is shown along the x-axis while the corresponding substitutions are shown on the y-axis. Polar residues (S, Y, Q, T, N, C) are colored blue and charged (R, H, K, D, E) in red. Fill values correspond to the relative strengths where decreased strength (purple) are negative values and increased strength (yellow) are positive values. White boxes represent missing or wildtype data, while gray boxes represent values where data was missing for one BioRep. b. Predicted AlphaFold2 secondary structure for SIN3B [31-111]. Residues where the average of all single mutant substitutions is greater than two standard deviations away from the mean of the WT values (**x̄ ± 2σ**) are colored purple, while those that are functionally-resilient to substitution and do not pass this threshold are colored yellow. c. Normalized Log_2_(OFF:ON) values for windowed-shuffle mutants of SIN3B [31-111]. For normalization, the average Log_2_(OFF:ON) is normalized to the mean of the WT values for each domain. Tiles span the residues covered by the indicated mutant. The normalized WT value is indicated by y = 0 (red) and one standard deviation above and below the WT values (**x̄ ± σ**) is indicated by the shaded region. d. Relationship between observed conservation and the effect of missense mutation on function. ConSurf Conservation Scores report on the conservation of residues (red), where lower scores are given to more highly conserved AAs. Average Log_2_(OFF:ON) values for a given residue are superimposed (blue), where lower values indicate lower repressor activity. Pearson’s R-squared and p-value (Wald Test) calculated from a linear least-squares regression between ConSurf Scores and Log_2_(OFF:ON) values are indicated in the top-right corner. e. (**a**) for KDM5B [261-341]. Known SUMOylation sites are underlined (IKIE, purple; IKQE, hot pink). f. (**b**) for KDM5B [261-341]. Residues where the average of all single mutant substitutions is greater than two standard deviations away from the mean of the WT values (**x̄ ± 2σ**) are colored purple, while those that are functionally-resilient to substitution are colored orange-yellow. Additionally, critical residues of the IKQE SUMOylation site are colored purple and hot pink. g. (**c**) for KDM5B [261-341]. h. (**d**) for KDM5B [261-341]. i. Schematic illustrating the workflow of the Functional <u>s</u>hort <u>li</u>near <u>m</u>otif (SLiM) Finder. Log_2_(OFF:ON) scores from the windowed-shuffle mutants are used to nominate motifs for consideration. These motifs are then further filtered and cross-referenced in the ELM database before being identified as a SLiM critical for the repressive function of the RD, within the context of the screen (**Methods: Functional SLiM Finder**). j. Selected SLiMs with the highest significance scores within disorder regions of their respective RDs, with colors representing their corresponding ELM annotation in the nucleus. Background scores for each domain, or the mean of all considered linear motifs, are shown in gray. Within SLiM labels, polar residues are colored blue, and charged residues are indicated in red, as in panel (**a**).

For the higher-order TWIST1 bHLH and ZNF705D KRAB mutants, we observed that different combinations of AA substitutions at the same positions can lead to varied outcomes in repression strength (**Supplementary Fig. 5**-7). For example, for triple substitutions in TWIST1 [95-185] at 102, 135, and 142, some combinations of substitutions result in LoF while other combinations enhance repressor activity, highlighting the substitution-dependent effects of these mutations (**Supplementary Fig. 6**). Similarly, in ZNF705D [2-82], triple substitutions at positions 20, 25, and 80 demonstrate a range of effects on repression strength (**Supplementary Fig. 7**). This variability underscores the potential complexity of predicting the effects of accumulating more than one mutation and highlights the necessity of more complex mathematical models, such as neural networks, for accurately predicting the outcomes of such mutations.

In disordered domains, which are not as well annotated with repressive function and sequence conservation, our data were able to identify functionally important motifs and SLiMs. For example, in the completely disordered KDM5B [261-341] RD, the most detrimental substitutions are within known SUMOylation sites (φKxE, <u>E</u>ukaryotic <u>L</u>inear <u>M</u>otif (ELM) accession ELME000251) (**Fig. 3e, f**, ^86,87^). AAs within and immediately adjacent to these sites are intolerant to shuffling, while the rest of the domain is relatively agnostic to a particular linear order (**Fig. 3g**). We identify the known TLE-binding SLiM (WRPW, ELM accession ELME000104) in the disordered HES3 [100-180] RD in the same manner (**Extended Data, Fig. 5d**). We found that substitutions in SUMO-interacting motifs in the disordered C-termini of SUMO1 [18-98] and SUMO3 [13-93] are especially perturbative in both family members (**Extended Data**).

Interestingly, the trends between conservation and functional sensitivity are less apparent in disordered domains: for instance, the residues in the SUMOylation sites of KDM5B [261-341] are not the only residues that are highly conserved, yet they are the only AAs found to be functionally important for repression (**Fig. 3h**). Some of these conserved residues contain annotated phosphosite motifs, which may be important for other modes of regulation. Overall, we reasoned that our data were therefore detecting contextually relevant patterns and would be suited for guiding a model designed to capture sequence level features that confer repressor activity, potentially with higher success than models trained on structure or homology alone.

Inspired by the trends we observed in disordered domains and regions, we developed an algorithm to identify all motifs that are essential for repression in our results, termed Functional SLiM Finder (**Methods: Functional SLiM finder**). We scanned all disordered domains and regions in our library and found significant functional motifs in 29 RDs (**Fig. 3i-j, Supplementary Fig. 8**-9). Our approach finds known ELM motifs, such as those previously mentioned: in some of these cases, we find that additional AAs immediately adjacent to the motifs are also critical for repression, suggesting that these residues may provide additional degrees of modularity to known SLiMs within the context of a specific RD (**Fig. j, Supplementary Fig. 8**). We also identify 19 previously uncharacterized functional motifs within disordered regions that are crucial for repression within the context of their 80AA RDs (**Supplementary Fig. 9, Supplementary Table 6**).

### A neural network predicting repressor function from AA sequence (Fig. 4)

Next, we developed neural network models encoding sequence, structural and biochemical features to map AA sequence to repressor activity and evaluated their ability to predict effects of sequence variation across diverse RDs in our RD-DMS dataset ^88^. Models were fit directly to the measured sequence read counts in OFF and ON populations using a binomial loss function, accounting for higher variance at lower read counts ^89^ (**Methods: Modeling repression strength**). Models predict the probability of each sequence in the OFF (*P*_*off*_) and ON populations (*P*_*on*_ = 1 − *P*_*off*_). We assessed performance based on the concordance of measured and predicted *P*_*off*_ for held-out sequences not used in training (**Supplementary Table 7**). We split RDs and their sequence variants into different training and test set configurations to systematically evaluate generalization of the models within and across RDs with varying levels of homology. We first evaluated model generalization across the highly homologous Chromo domains (Pfam ID: PF00385), CBX1, CBX3 and CBX5, which show high structural homology and similar functional patterns despite some sequence divergence (**Supplementary Fig. 10a**-c**, Extended Data**) ^90,91^. We used sequence variants of CBX5 [109-189], CBX3 [104-184], and CBX4 [481-561] (an unrelated Cytochrome b5-like Heme/Steroid binding domain from a Polycomb family member) for training, while reserving those of CBX1 [105-185] as a test set to assess prediction accuracy on a held-out, homologous domain.

**Figure 4:**
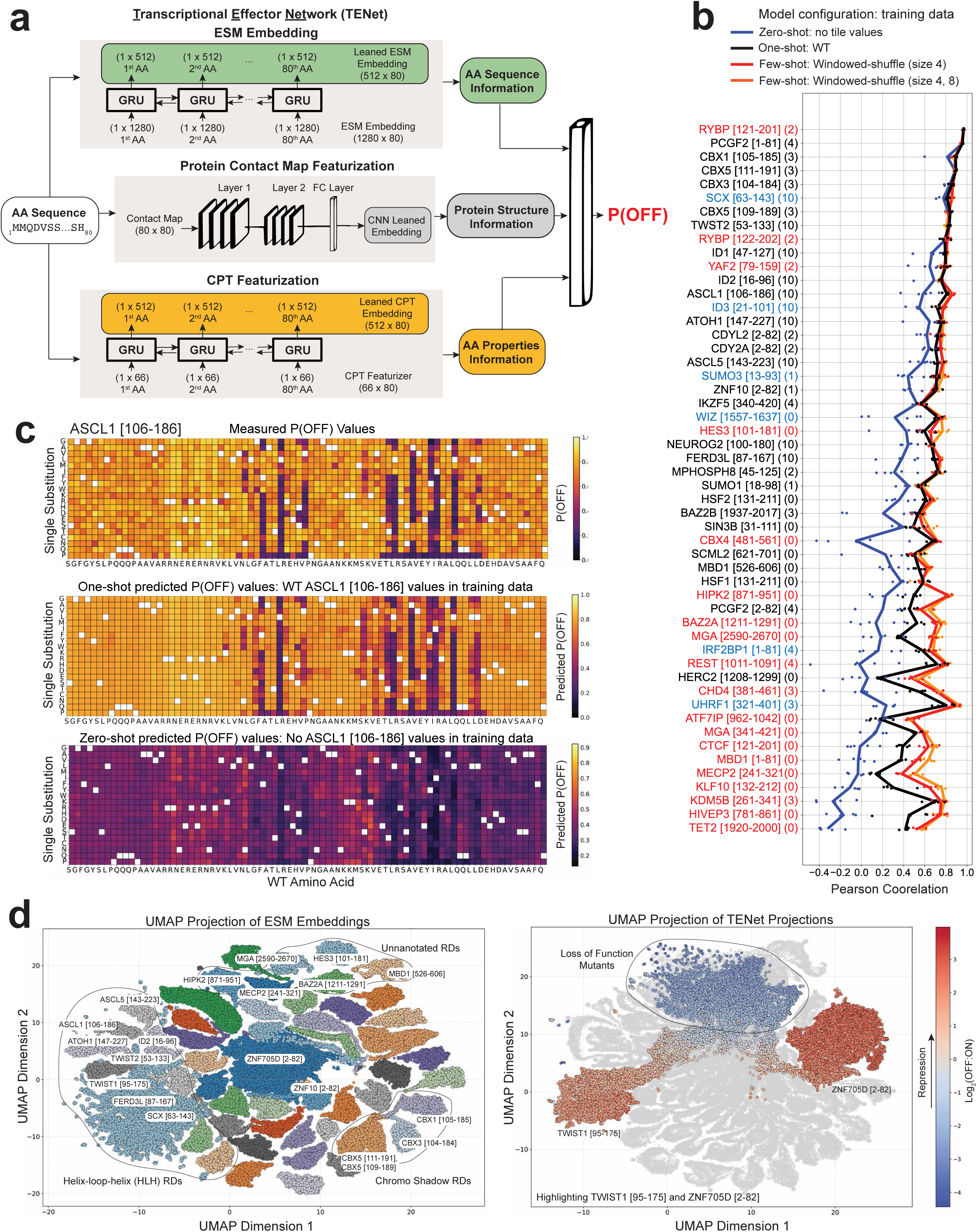
TENet: a neural network predicting repressor strength from AA sequence. a. Schematic representation of <u>T</u>ranscriptional <u>E</u>ffector <u>Net</u>work (TENet), which is designed to model repressor strength. The model incorporates three primary featurizers: ESM embeddings, protein contact maps derived from ESMFold, and CPT AA descriptors. The sequential characteristics of ESM embeddings and CPT descriptors are captured using a bi-directional GRU. The protein contact maps are processed through two convolutional layers. The integrated output from these featurizing layers is subsequently channeled into a fully connected neural network, culminating in the prediction of P(OFF) values (**Methods: Neural Network**). b. Pearson correlations between measured vs. predicted P(OFF) values for single mutants of respective domains (y-axis) as test sets. Four configurations were investigated by utilizing training data subsets that either excluded domain reads (Zero Shot), included wild-type reads (One Shot), or included windowed-shuffle reads of sizes 4 and/or 8 (Few-Shot). Data points are results of six independent runs with the lines passing through the average values, per configuration. Y-axis domain labels are colored to indicate structured (black), hybrid (blue), and disordered (red) domains. c. Heatmaps of the measured (top) and predicted (middle, bottom) P(OFF) values for the single substitutions of the ASCL1 [106-186] domain. Predicted P(OFF) values are shown for one-shot (middle) and zero-shot (bottom) training configurations. d. UMAP visualization of all RDs and variants using ESM embeddings (left). Projections are color-coded by gene. Three major domain families are highlighted: Helix-Loop-Helix (HLH) domains, Chromo shadow domains, and unannotated domains. UMAP visualization of all EDs and variants using embeddings from our trained neural network, TENet (right). Data for ZNF705D [2-82] and TWIST1 [95-175] are colored by Log_2_(OFF:ON) repression strength.

**Figure 5:**
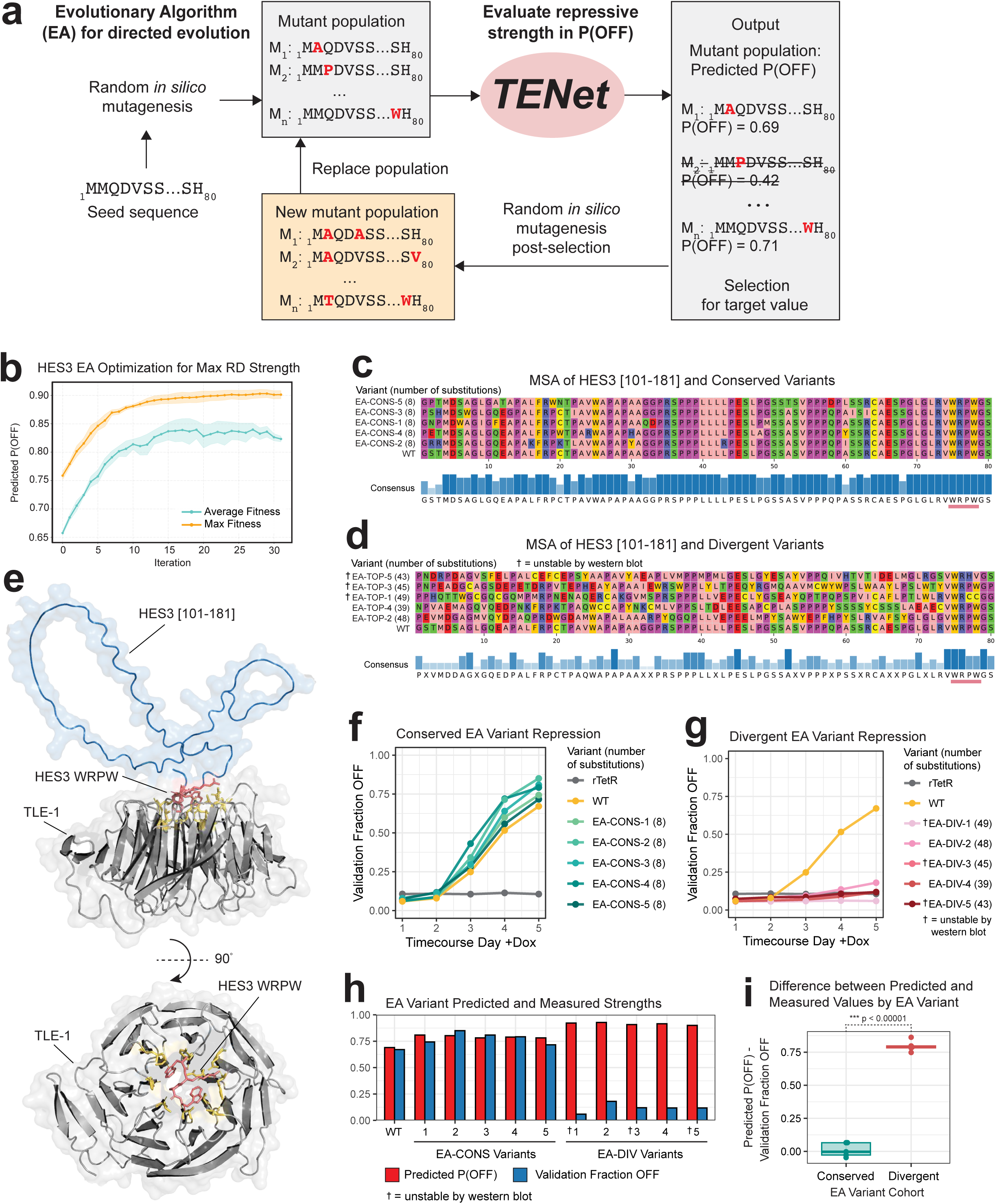
Using an evolutionary algorithm in conjunction with TENet to increase the repressive strength of a disordered domain. a. Overview of the evolutionary algorithm (EA) combined with TENet to perform *in silico* mutagenesis and directed evolution of RDs. From a selected seed sequence, a population of mutants are generated. Then TENet predicts the repressive function of each of the mutants, and mutants with values closest to the target strength are selected for while the remaining are withdrawn from the population. The selected mutants are used as an input for the next generation. The cycle iterates until sequences close to the desired values are obtained (**Methods: Evolutionary algorithm**). b. The trajectory of predicted P(OFF) values for evolved mutants of WT HES3 [101-181] where the target value is the maximum tenable P(OFF). The average (turquoise) and max (yellow) P(OFF) of the mutant populations generated in all 5 trials are shown vs. EA iteration. The shaded regions represent +/- one standard deviation (**x̄ + 1σ**). c. AlphaFold Multimer predicted interaction between the WRPW SLiM of HES3 [101-181] (red) in complex with the WD40 β propeller domain of TLE1 (gray). The rest of the HES3 RD is colored in blue (top). WRPW is predicted to facilitate recruitment of TLE1 through direct binding interactions with a combination of hydrophobic and polar residues (yellow) in all seven blades of the propeller (bottom). d. MSA of the WT HES3 [101-181] domain and all evolved, relatively conserved variants. The WRPW motif is underscored (coral bar). e. (**i**) for all divergent evolved variants. Variants that were unstable by western blot (**Supplementary Fig. 13d**) are indicated by an asterisk. f. Timecourse for individual “conserved” evolved variants showing the fraction of cells with a silenced citrine reporter measured 1-5 days after the addition of doxycycline. g. Timecourse as in (**f**) for individual “divergent” evolved variants. Variants that were unstable by western blot (**Supplementary Fig. 13d**) are indicated by an asterisk. h. Paired-bar plots of the predicted P(OFF) values (red) and fraction of cells with a silenced citrine reporter in single validation experiments (blue) for all evolved mutants. Variants that were unstable by western blot (**Supplementary Fig. 13d**) are indicated by an asterisk. i. Log_2_(OFF:ON) values for human variants of the basic (left) and helix-loop-helix subdomains of TWIST1 [95-175]. Variants are classified by reported clinical significance. j. The difference between the predicted P(OFF) and measured validation fraction OFF for evolved variants by cohort. We report the significance of the difference between the cohorts by Welch’s t-test.

We trained a fully-connected neural network anchored on one-hot encoded AA sequences on this data ^92^. However, we observed very poor performance on the single mutant variants of the CBX1 domain in the test set (*r* = 0.13), suggesting that even for highly homologous RDs, this model is likely underpowered to learn generalizable features ab-initio from AA sequence.

Hence, we turned to potentially more generalizable encodings and representations derived from the AA sequences. First, we used the protein language model ESM-2 to extract amino-acid resolution embeddings for all sequences in our dataset ^37^. ESM-2 embeddings encode homology without the need for multiple sequence alignments. ESM2 embeddings also enable accurate protein structure prediction from AA sequence via ESMFold ^37^. Hence, we also extracted 2D contact maps from the predicted ESMFold structures for each of our sequences. Third, we encoded biochemical and biophysical properties of individual AAs in our sequences using the Cross-Protein Transfer (CPT) AA descriptors ^42^. We suspected that these CPT encodings may improve predictions for disordered domains, which often have minimal sequence homology with other domains. We used separate neural networks composed of bi-directional Gated Recurrent Units (GRU) ^93^ anchored on the AA resolution sequential encodings from ESM-2 embeddings and CPT descriptors respectively (**Methods: Neural Network**). We used a convolutional neural network (CNN) architecture ^94^ anchored on the contact maps represented as 2D images, since CNNs are ideally suited to learn multi-scale features from imaging data. We trained separate models on each of the three representations on the Chromo domain data split and evaluated their performance on the different types of CBX1 sequence variants (fully scrambled, single mutants, window shuffled and all mutants) in the test set (**Supplementary Table 8**). The ESM2 model performed the best (*r* = 0.82, all mutants), followed by the CPT model (*r* = 0.8, all mutants). The ESMFold contact map model performed relatively poorly (*r* = 0.34, all mutants). However, each of these models outperformed the one-hot encoded fully connected model, showcasing the importance of pre-trained embeddings for generalizability.

We then trained and evaluated models that integrated multiple combinations of embeddings via fully connected layers assimilating representations from each of the submodels anchored on each embedding (**Methods: Neural Network**). Integrating ESM2 embeddings with ESMFold contact maps enhanced performance (*r* = 0.86, all mutants), and integrating all three types of embeddings yielded our most effective model (*r* = 0.91, all mutants) (**Supplementary Fig. 10f**) (**Supplementary Table 8**), which we named <u>T</u>ranscriptional <u>E</u>ffector <u>Net</u>work (TENet) (**Fig. 4a**). Notably, a significant portion of this performance gain was observed in our subset of domain-specific negative controls ("Fully scrambled Mutants"). These mutants, characterized by AA sequences that retain the same AA composition as the WT CBX1 RD but with a randomized order, highlight capacity to recognize the functional significance of specific arrangements of AA, a critical facet of understanding repressor domains.

We further benchmarked TENet against PADDLE ^6^, a 6-layer CNN designed to predict ADs from one-hot encoded AA sequences. Similar to our approach, PADDLE leverages DMS data from high-throughput activation assays conducted in yeast. For direct comparison, we re-trained PADDLE specifically on our CBX subset task. Similar to our fully connected one-hot model, PADDLE demonstrated poor generalizability on this task (*r* = 0.06) (**Supplementary Table 7**). Additionally, we experimented with graph convolutional and attention architectures ^95,96^ (**Supplementary Table 8**). Here, structure predictions from ESMFold were used to construct a graph scaffold with AAs as nodes and spatially adjacent AAs/nodes connected by edges. We investigated two approaches for constructing the graphs: 1) A complete graph (GCN - Complete Graph) where all nodes were interconnected and edges were weighted by the alpha-carbon distance between AA pairs, and 2) a sparse graph (GCN - Sparser) where edges were established between two AAs only if their alpha-carbon distance was under 5 Angstroms (Å). Each AA node in the graph was encoded with its ESM-2 embedding and CPT AA embedding. Interestingly, these graph models performed similarly to our GRU models anchored on ESM-2 embeddings alone (**Supplementary Table 8**), but fell short of TENet, despite the graphs being provided with equivalent information. Lastly, we evaluated the significance of using a sequential model to learn how to aggregate AA-resolution ESM2 embeddings: when we simply used ESM-2 embeddings averaged over all AA positions for each mutant sequence, we observed a very large decline in performance compared to TENet (**Supplementary Table 7**).

Next, we used hold-one-out data split configurations to systematically evaluate the performance of TENET in predicting the activity of each held-out RD while using all other RDs for training (**Fig. 4b, Methods: Neural Network, Model Training**). Four distinct hold-out scenarios were examined: 1) zero-shot prediction, where the model is provided with no sequences from the test-set RD; 2) one-shot prediction, where only the reads from the wild-type (WT) sequence of the test-set RD are included in training; and 3, 4) few-shot scenarios, where windowed-shuffle variants of length 8AA or both sizes 4AA & 8AA, respectively, from the test-set RD were provided as training data. These configurations were chosen to assess model generalization to held-out RDs as a function of their homology relative to RDs in the training set and the amount of exposure to the sequences from the held-out RDs in training ^97,98^. We observed a systematic decay of zero-shot prediction performance with decreasing homology of held-out RDs with respect to RDs in the training set. For instance, held-out Chromo domains exhibited high accuracy in zero-shot predictions due to strong homology with other Chromo domains in the training set. In contrast, zero-shot performance on a held-out TET2 domain was the worst (*r* = −0.3) due to very low homology with other RDs in the dataset. However, in several cases such as the ASCL1 HLH domain with moderate (*r* = 0.75) zero-shot performance, we observed visible contrasts between predicted activity of strong repressive mutants and those with no effects, suggesting that the models may have the capacity to generalize better with in a one-shot and few-shot setting (**Fig. 4c, bottom panel**). Indeed, one-shot and few-shot configurations improved performance (**Fig. 4b**). For the ASCL1 HLH domain, exposure to only the WT sequence in the one-shot setting improved performance (*r* = 0.82) and calibration (**Fig. 4c, middle panel**). Even for the challenging RDs in the zero-shot settings, such as the TET2 domain, one-shot and few-shot configurations resulted in dramatic improvements (*r* = −0.3, zero-shot to *r* = 0.45-0.6, one and few-shot) (**Fig. 4b**).

We also measured the effect of including higher-order mutants in training on predictive performance. Specifically, we trained additional TENet models excluding all triple substitution variants and either with or without the TWIST1 [95-175] and ZNF705D [2-82] double substitution variants, and then evaluated their performance on the held-out triple mutants (**Methods: Neural Network**). Including double mutant variants in training improved performance on held-out triple mutant variants for each RD. This effect was more pronounced for ZNF705D [2-82] (**Supplementary Table 9**).

Finally, we compared the information encoded in TENet embeddings relative to their source ESM2 embeddings by estimating distances for all pairs of sequences using these embeddings and visualizing them in 2D with the Uniform Manifold Approximation and Projection (UMAP) dimensionality reduction algorithm (**Fig. 4d**). ESM2 embeddings clustered sequence variants more strongly based on structural homology of their source RDs (Silhouette score = 0.369) regardless of their repression activity (Silhouette score = −0.034) (**Fig. 4d, Supplementary Fig. 11, a-f, Methods: 2D Projection**). In contrast, TENet embeddings clustered sequences more strongly based on their repression activity (Silhouette score = 0.237), than structural homology (Silhouette score = 0.080) (**Fig. 4d**, **Supplementary Fig. 12**). This suggests that while ESM2 embeddings are biased towards structural features, they likely encode repression-relevant features as well. TENet leverages and transforms these features when trained on repressor activity, thereby delivering powerful generalizable predictions of sequence variation across diverse RDs even in few-shot settings.

### Guiding *in silico* directed evolution of repressor domains (Fig. 5)

Because TENet identified sequences important for repression, we next tested whether it could be used to guide the design of repressor domains to tune their repression desired activity. We integrated TENet into an evolutionary algorithm (EA) to drive *in silico* directed evolution ^99^ (**Methods: Evolutionary algorithm**). In this framework, TENet evaluates the repressive strengths of mutant populations as a measure of fitness, then the EA applies evolutionary pressure towards the desired target activity through simulated selection (**Fig. 5a**). The EA first performs random mutagenesis of a single seed sequence to generate a mutated population. TENet then predicts *P*_*off*_ values for each mutant, and a subpopulation with predicted activity closest to the desired target *P*_*off*_ are selected as seed sequences for the next generation. Through multiple rounds of mutagenesis and selection, the mutant RDs trend toward the desired repression strength (**Fig. 5a, b**).

We hypothesized that the *P*_*off*_ values of our evolved designs would be more accurate if the seed sequence were selected from our RD-DMS data, since TENet’s predictions were much better in the one-shot and few-shot settings, where it was exposed to related sequences in training. We also aimed to tune the activity of a disordered domain, a novel challenge in the field ^100^. We tasked the EA to evolve the WT HES3 RD (*P*_*off*_ = 0.66), a disordered domain without a known family annotation (**Fig. 1f**), into a stronger repressor (maximization task). We believed this task was feasible given the numerous examples of stronger repressors in our RD-DMS data. To evaluate reproducibility and evolutionary trends, we conducted five separate trials. In each trial, the maximum predicted *P*_*off*_ achieved by mutant populations increased with each iteration of the EA, plateauing around 0.9 (**Fig. 5b**). The evolved populations tended to accumulate more mutations and adopt secondary structures over time (**Supplementary Fig. 13a**, b). Given TENet’s higher predictive accuracy for structured domains (**Fig. 4b**), the predicted *P*_*off*_ values for mutants with recognizable structural patterns were less variable, suggesting a positive feedback loop favoring increased structural features in mutants moving towards the desired *P*_*off*_ values. This hypothesis is supported by the trajectory of evolved mutants within the latent space of ESM2 embeddings, which captured structural similarity trends toward structured KRAB domains (**Supplementary Fig. 13c**).

Previous protein engineering attempts have found that designs with fewer mutations tend to have higher success rates ^26,101–103^. We selected two cohorts of evolved HES3 mutants for validation to assess whether the EA powered by TENet would yield better designs with increased homology to the training data. The first set of “conserved” variants had predicted *P*_*off*_ values increased through the acquisition of 8 AA substitutions (10% divergence from the 80AA seed sequence), and the second set of “divergent” variants were predicted to have the highest *P*_*off*_ values, regardless of the number of accumulated substitutions. The conserved variants retained high homology to both the seed sequence and other HES3 mutants in the training data, but had not been tested in the RD-DMS screen (**Fig. 5c**). In contrast, the divergent variants acquired more mutations and were far less homologous to the seed sequence (48-62% divergence with 39-49 substitutions) (**Fig. 5d**).

Consistent with other disordered domains, the HES3 RD was mostly unaffected by single substitutions, though mutations in the WPRW SLiM near the C-terminal end significantly reduced repression (**Extended Data**). This aligns with observations that the WRPW SLiM is essential for binding the TLE family of co-repressors via the WD40 β propeller domain ^104–107^. In the predicted HES3 RD and human TLE1/4 complex, every residue of the WRPW SLiM contributes to interactions with the cofactor, while the rest of the domain does not (**Fig. 5e**). Notably, the complete WRPW domain was retained in all conserved variants and most divergent variants, with the N-terminal tryptophan and arginine retained in every variant (**Fig. 5c, d**). All variants stable as measured by Western blot retained the entire WRPW SLiM (**Supplementary Fig. 13d**), suggesting that our pipeline recognized the WRPW SLiM’s importance for repression and applied selective pressure to maintain it.

We measured the repression strengths of the evolved cohorts in our reporter cell line. For the conserved cohort, the fraction of cells in the OFF state increased for all variants compared to WT (**Fig. 5f**). In contrast, all variants in the divergent cohort lost their ability to mediate repression, which could partly be attributed to protein instability (**Fig. 5g, Supplementary Fig. 13d**). The *P*_*off*_ values predicted by TENet matched the measured fraction OFF for the conserved cohort (**Fig. 5h**), though the prediction error was significantly higher (*p* < 0.00001; Welch’s *t*-test) for the divergent cohort (**Fig. 5i**). These results indicate that TENet is less effective at guiding directed evolution for sequences that significantly diverge from the training data. Nonetheless, the evolution of stronger HES3 variants in the conserved cohort demonstrates that the EA powered by TENet can effectively optimize proteins, even disordered domains, when constrained to a degree of homology with the seed sequence.

### Engineering synthetic repressors with specified strengths (Fig. 6)

To systematically characterize when the evolutionary algorithm (EA) powered by TENet yields the most accurate designs, we constructed a library of evolved RDs by sampling mutants with incremental degrees of sequence divergence along multiple evolutionary trajectories and measured their activity using HT-recruit (**Fig. 6a, Methods: EA-RD library design**). To determine if the target repression strength affects the relationship between predictive accuracy and homology, we tasked the algorithm with evolving sequences toward varying *P*_*off*_ values indicative of weak (0.4), moderate (0.6 or 0.7), and strong repression (maximization task), starting from the same seed sequence (**Fig. 6a**). Additionally, to assess if the choice of seed sequence impacts model performance, we repeated each task using five different seed sequences (WT sequences for CBX5, HES3, ID2, MECP2, and NEUROG2 RDs). These sequences represented varying degrees of predicted structure, measured strength, and homology in the RD-DMS training data (**Fig. 6a**, **Fig. 1c, Extended Data**).

**Figure 6:**
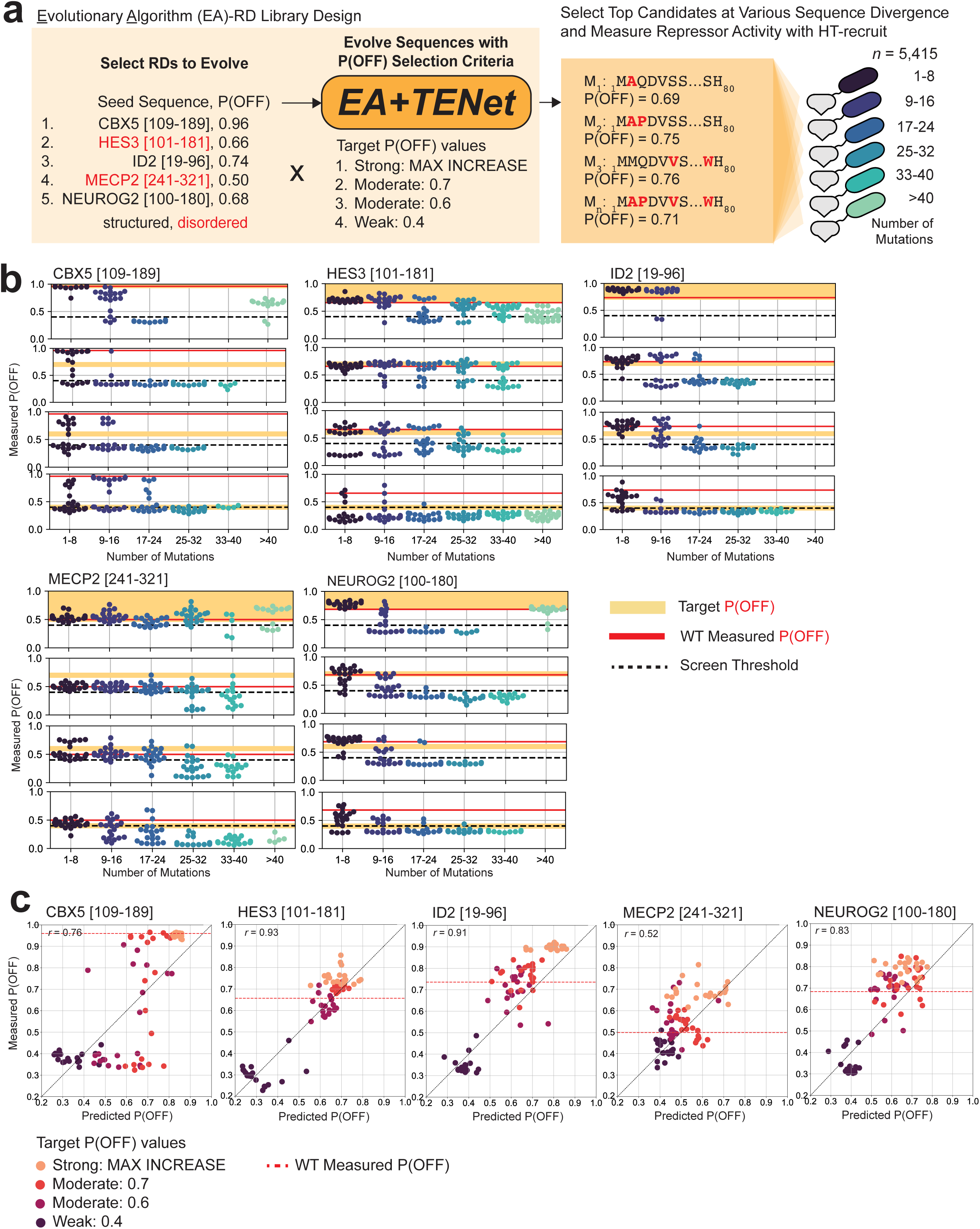
*In silico* directed evolution of repressor domains with targeted strengths a. Schematic illustrating the design and compositions of the EA-RD library. Seed sequences (5 WT RDs) are independently evolved toward specified target P(OFF) values (4 tasks). The predicted P(OFF) values are evaluated as selection criteria. The candidate sequences predicted to function closest to the target strengths were ordered as oligos and cloned as fusions to rTetR (in a pooled format). This selection is repeated separately for various categories of sequence divergence (number of mutations). b. Distributions of the Measured P(OFF) values for all evolved mutants, stratified by seed sequence. Results are subgrouped by target P(OFF) values, from top to bottom, in the order MAX INCREASE, 0.7, 0.6, and 0.4. The dotted black line delineates the screen threshold as a P(OFF) value and the solid red line indicates the measured P(OFF) value of the seed sequence. Target P(OFF) values are shaded in yellow, for each task. Evolved mutants are grouped by number of mutations or substitutions (x-axis). c. Correlations between predicted P(OFF) and actual P(OFF) values measured by the EA-RD screen for 15 mutants that survived through the last iteration of evolution for each indicated target value. The Pearson’s r is indicated in the top left corner of the plot of each respective dataset.

Based on these results, we designed a new experimental library—the <u>E</u>volutionary <u>A</u>lgorithm <u>R</u>epressor <u>D</u>omain (EA-RD) library—containing 1,805 mutants spanning a broad range of predicted P(OFF) values (**Supplementary Fig. 14a**). These sequences were encoded by three oligos with varied usage of degenerate codons to improve measurement reliability. An HT-recruit screen with the EA-RD library produced high-quality data, showing strong correlation between biological replicates (*r* = 0.9861, CCC = 0.9855) and well-distributed sequencing counts across all library samples (**Supplementary Fig. 14b**, c). This screen identified 99.58% of negative and 96.49% of positive controls correctly (**Supplementary Fig. 14d**), with low variability in measurements for sequences encoded by more than one codon (average σ = 0.1424). Additionally, the standard deviation of Log_2_(OFF:ON) ratios for all library elements encoded by multiple codons was not dependent on the average Log_2_(OFF:ON) ratio (**Supplementary Fig. 14e**).

Next, we assessed the EA’s accuracy in fine-tuning repressor activity across a range of strengths and aimed to pinpoint its limitations. Design accuracy, defined as the ability to evolve variants closest to target values, was highly dependent on the number of accumulated mutations or percent identity: design accuracy decreased when evolved sequences accumulated more than 16 mutations (<80% percent identity), while the designs closest to the target values had only 1-8 mutations (>90% percent identity) (**Fig. 6b, Supplementary Fig. 15**). Additionally, the *P*_*off*_ value of the seed WT sequence was crucial; larger differences between starting *P*_*off*_ and target *P*_*off*_ correlated with lower design accuracy (**Fig. 6b, Supplementary Fig. 15**). This finding underscores a critical tradeoff between achieving maximal design improvements and maintaining sequence similarity, a balance previously noted in other studies ^26–28^. However, we also observed that design accuracy for variants with more than 16 mutations was higher for disordered domains (HES3 and MECP2 RDs) than for structured domains, suggesting potential differences in the risk-reward balance based on domain rigidity.

We also evaluated the accuracy of TENet’s predictions for the evolved variants. We found strong correlations between predicted and measured values among the top 15 mutants that persisted to the end of the EA across all domains (average *r* = 0.79) (**Fig. 6c**). The most accurately predicted mutants generally accumulated fewer than 16 mutations (>80% percent identity), indicating a threshold beyond which TENet’s prediction accuracy declines, likely due to the model making out-of-distribution predictions. This trend was consistent for both structured and disordered domains, demonstrating the versatility of our modeling approach. Overall, the results from the EA-RD screen highlight TENet’s capability to guide the editing of RDs to tune their activity.

## Discussion

In this study, we present an integrative approach using machine learning models trained on large-scale DMS assays to characterize and design transcriptional repressor domains (RDs). Our dataset, one of the largest DMS screens of repressor activity in diverse human protein domains, powers our deep learning model, TENet, which accurately predicts repressor activity from amino acid sequences with minimal training data. TENet, incorporated into an *in silico* evolution framework, allows us to tune the repressor strengths of both structured and disordered domains, achieving high design success rates confirmed experimentally. This work marks a significant advancement in sequence-to-function modeling and design, enabling the directed evolution of new effector domains by fine-tuning their functional properties.

Clinically, our work offers a functional interpretation of 1,264 disease-associated variants, previously classified as pathogenic or VUSs, highlighting variants that may disrupt regulatory activity of RDs. Our assays provide a complementary readout to previous DMS screens, which focused on protein stability, enzymatic function and cell fitness ^108^. These insights could inform future studies exploring somatic mutations in cancer or disease-associated coding variants in GWAS loci. While our work used the HT-recruit system in K562-derived lines, future validation in disease-relevant tissues and native genetic loci will be critical.

Our study also illustrates the utility of rationally designed mutants to map functional features within complex domains, such as identifying short linear motifs (SLiMs) in disordered RDs. Window-shuffle mutants proved essential in identifying SLiMs, while single mutants helped pinpoint key residues involved in repression, offering insights into regulatory mechanisms like post-translational modifications ^109,110^.

Our study demonstrates the potential of predictive models to extend the utility of DMS screens to proteins and variants that are not directly assayed. Traditionally, these models have relied heavily on structural and evolutionary constraints as proxies for function ^111^. However, the poor correlation between conservation and functional effects in our repression screens suggests that these constraints are not particularly useful for deciphering functional features of unstructured, evolutionarily divergent disordered domains ^7,112,113^. Our TENet model shows that repression activity can be accurately predicted from sequence alone, but emphasizes the importance of training directly on experimental repression data.

Through systematic exploration of model design choices, we uncovered several key insights. First, despite the size and diversity of our DMS dataset, ab-initio models trained on raw sequence tend to overfit and struggle to generalize, even to highly homologous protein domains. Sequence embeddings and structural contact maps from the ESM2 pre-trained protein language model significantly enhanced prediction performance and generalization. While protein language models primarily encode evolutionary and structural constraints, our results suggest that ESM2’s latent representations capture features relevant for functional prediction, such as repression activity. The best-performing models incorporated additional biochemical features, which proved especially beneficial for disordered domains. Thus, TENet highlights the power of combining sequence and structure representations from protein language models with biochemical properties for accurate protein function prediction.

Second, homology between training and test sequences significantly influenced model performance, with accuracy declining as evolutionary distance increased, particularly for divergent disordered domains. Remarkably, the inclusion of even a single functional measurement (one-shot context) of the wild-type sequence led to dramatic improvements in generalization to diverse variants of the same domain. Including a few windowed mutants (few-shot context) further improved performance. Higher-order mutants, such as double and triple substitutions, also boosted model accuracy, which is particularly important for refining models to address non-additive and epistatic fitness landscapes—an ongoing challenge in protein modeling. These findings suggest cost-effective strategies for future library designs, maximizing prediction accuracy and generalization while minimizing library size by carefully selecting exemplar domains and balancing the depth and breadth of mutants. Active learning strategies, involving iterative data generation, model training, and model-assisted library design, could further enhance performance and generalization while reducing experimental costs.

Our study also makes important contributions to protein engineering through the *in silico* design of proteins with desired functional properties ^25^. Previous machine learning approaches for protein design have primarily focused on optimizing structural properties ^26–28^. However, this approach is not well-suited for repression domains, which are predominantly disordered ^28,118–120^. To our knowledge, no computational methods have specifically addressed the challenge of designing or tuning disordered effector domains, representing an unexplored frontier in synthetic design. We introduce a simple yet effective strategy using TENet within an *in silico* directed evolution framework to optimize the activity of both structured and disordered RDs, without the need for detailed structural information. Our designs demonstrated high success rates across a broad range of activities in experimental evaluations. However, the framework was more effective when sequences were not highly divergent from the seed sequences, as TENet tends to make less accurate predictions for out-of-distribution variants. Future improvements could include incorporating approaches that explicitly flag out-of-distribution predictions to enhance performance in such cases. Despite this limitation, our models and design framework already offer substantial utility for designing compact, precise, and potent domains to modulate gene expression at desired strengths, particularly when structural information is lacking ^29,30,121,100^. More broadly, our study exemplifies how deep learning models, combined with high-throughput experimental techniques, can advance the understanding of gene regulation and protein design.

Together, this study lays the groundwork for future modeling efforts that will couple DMS assays with deep learning models and novel model-driven, cost-effective library design strategies to decipher the functional effects of coding variants in disease and design transcriptional effectors for precise modulation of gene expression.

## Acknowledgements

We thank Abby Thurm, Nicole DelRosso, David Yao, Tanner Jensen, and members of our laboratories for helpful conversations and assistance. This work was supported by NIH/NHGRI R01HG011866 (M.C.B. and L.B.), Stanford Bio-X Bowes Fellowship (R.V.), Stanford Bio-X Interdisciplinary Graduate Fellowship (A.N.), and NIH-4K00DK126120-03 (J.T.).

## Contributions

A.N., R.V., A.K., L.B., J.T. and M.C.B. designed the study. A.N. designed the RD-DMS library with significant contributions from J.T. and R.V. R.V. designed the CV-RD library, and A.N. and R.V. designed the EA-RD library. K.S. cloned all libraries. K.S. and R.V. performed quality control on all libraries with help from J.T. R.V. performed all high-throughput screens and next-generation sequencing. R.V. processed raw screen data and R.V and A.N. analyzed results. A.N. generated all predicted 3D models. R.V. cloned all individual and validation constructs with help from S.A. and A. R.V. performed individual recruitment assays, Western blot experiments, and RNA-seq analyses. R.V. performed analysis of clinical and pathogenic variants with help from R.P. and P.D. A.N. designed all neural networks and evaluated performance with A.K. and help from P.S. A.N. designed and employed the evolutionary algorithm. R.V., A.N., L.B., J.T., A.K. and M.C.B. wrote the manuscript with significant contributions from P.S., along with contributions from all authors. M.C.B., A.K., and L.B. supervised the project.

## Methods

### Domain classification and disorder prediction

Each 80AA wild-type (WT) repressor domain (RD) was queried in the InterPro database ^53^ and classifications were derived either from Pfam ^51,52^ or PROSITE annotations ^122^. For domains which fully contained or overlapped with greater than 80% of an annotated Pfam, the Pfam annotation was used to classify the domain family. If the domain did not contain a Pfam annotation but did contain a PROSITE profile, the PROSITE annotation was used for family classification. If a domain did not contain either a Pfam or PROSITE annotation, it was considered “Unannotated” in our study.

Computational predictions of percent disorder and calculations of longest disordered subdomains were performed with metapredict V2-FF (V2.6) ^39^ in Python 3.8.5. From these quantifications, each WT RD was categorized as “disordered”, “hybrid”, or “structured”. Domains were classified as disordered if they contained a contiguous disordered region equal to or longer than 30AA (>37.5% of the domain) or were predicted to be greater than 50% disordered. Else, if >30% of the total domain was predicted to be disordered, it was considered hybrid. Domains that did not meet either of these criteria were classified as structured.

### 3D structure prediction

To generate 3D structural renderings of our RDs, we utilized the AlphaFold2 algorithm ^34^ via the ColabFold (V1.3.0) implementation (<u>GitHub</u>). Specifically, we made use of the "AlphaFold2_mmseqs2" notebook. In this computational workflow, a single query was submitted for each RD, and templates were sourced from the "pdb70" database. For the multiple sequence alignment (MSA), the MMseqs2 algorithm was set to search against the UniRef+Environmental database. We employed the AlphaFold2-ptm model type and generated a total of five models for each sequence with three recycles performed during folding simulation. Ensemble averaging considered only a single model, and the generated models were ranked based on their pLDDT scores. Both unpaired and paired sequences were considered in the computational process.

### RD-DMS library design

54 human RDs were selected to be representative of multiple and variable domain families, and to sample a broad range of repressor strengths as quantified in previous HT-recruit screens ^19,21^. We first selected the 1-2 strongest RDs from Pfams with significant hits from previous datasets. Then 11 domains were chosen from the large HLH Pfam (PF00010) with varied repressor strengths and bifunctional activities. To sample previously unannotated RDs, we used data from the unbiased tiling screen and considered all tiles that had measured repressor activity above the screen thresholds, but which did not overlap an known Pfam annotation. We selected the strongest 80AA tile within each repressor peak, which could comprise several overlapping tiles. Then, we then designed 1,520 all-by-all single mutants for each RD, encompassing every possible single amino acid substitution. Windowed-shuffle mutants were created using sliding windows of 4 or 8 AA, with a step size of 2 AA from the N- to the C-terminus of each RD. These mutants were either a 1) reversal or 2) complete randomization of the AA sequence within each window. In total, 80 reversal and 80 randomized windowed-shuffle mutants were generated for each RD. For ZNF705D [2-82] and TWIST1 [95-175], we further designed 6,320 double mutants and 6,000 triple mutants per RD. The double mutants were characterized by double alanine substitutions at all possible locations and in all possible combinations across the WT RD. The triple mutants were generated through random combinations of bulky and/or charged residues (tyrosine, proline, tryptophan, histidine, valine, glutamic acid) and alanine across 20 randomly-selected positions over the 80AA RD. Additionally, we incorporated 3,546 negative controls into our study. These controls comprised two categories: 1) 1,419 randomizations of our RDs, and 2) 2,127 sequences previously identified as having no repressor activity. 50 degenerate sequences each with a unique DNA encoding were generated for each WT RD to serve as positive controls within our library. To facilitate distinct identification of each library component during short-read sequencing, we required uniqueness within the first 120 bp of the DNA sequence of each library element. Additionally, sequences were codon matched for human codon use, 7xC homopolymers were removed, BsmbI restriction sites were removed, rare codons (less than 10% frequency) were avoided, and the GC content was constrained to be between 20-75% in every 50 bp window. All sequence requirements were resolved using DNAchisel ^123^ with a process of codon optimization to re-encode each element until the criterion for uniqueness and other features was met.

### CV-RD library design

To create a list of RDs to consider for the CV-RD library, all measurements from previous Chromatin Remodeler (CR) and Transcription Factor (TF) tiling HT-recruit screens measuring repression at either pEF or PGK promoter were queried ^21^. All tiles that passed the repressor threshold for their respective screens were considered RDs. For the CV-RD Library, we only considered tiles that were considered RDs at both the pEF and PGK promoter, as these were most likely to have less context-specific activity. All missense variants that fell within these defined RD windows were queried from the ClinVar database ^9^. In cases where RDs were defined by multiple tiles in the original screen and thus had coordinates that exceeded 80AAs in length, we truncated and defined 80AA tiles as follows: if variants were within the first or last 80AAs of the RD, the first or last 80AAs of the RD were selected, respectively. Otherwise, 30AAs of the RD were cropped from the N-terminus of the RD until the variant fell within the first 80AA of the cropped domain. Oligos were designed to encode all variants within their respective 80AA RDs, and 3 degenerate WT tiles were designed per RD to act as positive controls within matching 80AA contexts. Altogether, this resulted in 3,093 library elements.

Controls for the CV-RD and EA-RD (below) libraries were shared in a pooled screen: for positive controls, three WT sequences for each of the 54 RDs used for the RD-DMS library were included, for a total of 162 positive control elements. For negative controls, 250 negative controls that scored as bonafide negative controls, or fell below the repressor threshold, of the RD-DMS screen were randomly selected and included in the screen.

### EA-RD library design

To evolve RD mutants for our EA-RD library, we employed the evolutionary algorithm (EA) to evolve five distinct RDs (CBX5 [109-189], HES3 [101-181], ID2 [19-96], MECP2 [241-321], NEUROG2 [100-180]) toward four tasks or predicted P(OFF) values (0.4, 0.6, 0.7, MAX). For each RD, we conducted five independent runs for each task. Each EA run consisted of 20 iterations, utilizing a population size of 500 members. All evolved mutants were concatenated across all five runs for each given RD and task. From each of these populations, we selected the top five mutants based on their fitness scores, or those with predicted P(OFF) values closest to the task target, at various mutational burdens: these ranged from 2-80 substitutions, or total number of substitutions divergent from the respective WT or seed sequence. Five mutants were selected at each even value of substitutions. Then, all selected mutants for all RDs and tasks were encoded using three degenerate DNA encodings, for a total of 5,415 sequences. Finally, all DNA sequences used to clone the individual mutants in the pilot HES3 study were added with two additional degenerate DNA sequences, for a total of 33 additional sequences in the EA-RD library. The positive and negative controls for this library are detailed in the description of the CV-RD library design (above).

### Library cloning

300bp oligonucleotides were synthesized as pooled libraries (Twist Biosciences) and resuspended to 10 ng/µL in Elution Buffer (EB). The RD-DMS Library was ordered as 3 sublibraries (of 39812, 43362, and 39812 elements), and the RD-Predict Library was broken into 2 sublibraries (of 7264 and 11320 elements). For each sublibrary, 6 x 50uL PCR amplification reactions were set up in a clean PCR hood using 5 ng template, 1uL of each 1mM primer (**Supplementary Table 3**), 1 µL of DMSO, 1 µL of 10mM dNTPs, 10 µL of 5X Herculase Buffer (Agilent), and 1 µL of Herculase II Fusion DNA Polymerase (Agilent, catalog no. 600679). The thermocycling protocol was 3 mins at 98°C, then cycles of 98°C for 20s, 61°C for 20s, 72°C for 30s, and then a final extension at 72 C for 3 min. The cycle number was optimized for each sublibrary to find the lowest cycle that resulted in a clean, visible product for gel extraction. 21-30 cycles were used for the RD-DMS sublibraries and 19 cycles were used for both RD-Predict sublibraries. After PCR amplification, the double-stranded DNA libraries were run on a 2% TBE gel with SYBR Safe Gel Stain (ThermoFisher, catalog no. 17-0329-42) and the products around the expected size (300 bps) were excised and purified following the QIAquick Gel Extraction Quick-Start Protocol using MinElute columns (Qiagen, catalog no. 28006). The target lentiviral recruitment vector pJT126 (Addgene no. 161926) was predigested with Esp3I (New England Biolabs (NEB), catalog no. R0734L) for 15 mins at 37°C and then purified by gel extraction. The sublibraries were then cloned into pJT126 as 16 x 10µL Golden Gate reactions consisting of 75 ng predigested backbone, 5 ng library (2:1 molar ratio of insert:backbone), 2 µL of 10X T4 Ligase Buffer and 1 µL of NEB Golden Gate Assembly Kit (NEB, catalog no. E1602L) with 65 cycles of digestion at 42°C and ligation at 16°C for 5 min each, followed by a final 5 min digestion at 42°C and then 20 min of heat inactivation at 70°C. The reactions were pooled and purified in MinElute columns (Qiagen), with elution in 6-12 µL of Nuclease Free dH_2_O. 4 uL of the elutant per tube were transformed into 50 mL Endura electrocompetent cells (Lucigen, catalog no. 60242-2) following the manufacturer’s instructions. Three tubes were transformed for all RD-DMS sublibraries and 1 tube was transformed for each of the RD-Predict sublibraries. After recovery, the cells were plated on either 30 (RD-DMS) or 10 (RD-Predict) 10 x 10 inch Luria-Bertani plates with carbenicillin such that about individual 100,000 colonies were seeded per plate, maintaining a minimum of 50X library coverage. The plates were incubated at 37°C to allow overnight growth, and the bacterial colonies were scraped into 20mL of LB per plate and collected in 500mL falcon tubes. Then the total volume was spun down and plasmid pools were extracted from the bacterial pellets using 2-3 columns of the Hi-Speed Plasmid Maxiprep kit (Qiagen, catalog no. 12663) per sublibrary. To determine the quality of the libraries, 1 µL of the purified plasmid pools was transformed and plated on a small agar petri dish with carbenicillin. After overnight incubation at 37°C, 20-30 colonies were picked for Sanger sequencing (Quintara) to estimate cloning efficiency and diversity of plasmid pools.

### Cell culture

All screens and *in vitro* measurements were performed in K562 (ATCC, CCL-243, female) derived lines. Cells were maintained at 37°C in 5% CO_2_ in RPMI1640 (Thermo Scientific, catalog no. 11875135) supplemented with 10% heat-inactivated fetal bovine serum (FBS) (Sigma-Aldrich, catalog no. F6765-500ML), 1% Penicillin-Streptomycin (P/S) (Gibco, catalog no. 15-140-163), and 1% L-Glutamine (Gibco, catalog no. 25030081). Heat-inactivation of the FBS was performed by holding the FBS at 56°C for 30 minutes in a sterile water bath before addition to media. The K562-JT039 pEF reporter lines were generated as described in ref. ^19^. Briefly, the pEF reporter was inserted at the *AAVS1* safe harbor locus by TALEN-mediated homology-directed repair to integrate the pEF reporter donor plasmid (pJT039, Addgene no. 161927). 1000 ng of pJT039 and 500 ng of each TALEN-L (Addgene no. 35431) and TALEN-R (Addgene no. 35432) plasmids were introduced to K562 cells via electroporation, then 1000 ng/mL puromycin antibiotic (Cayman Chemical no. 28240) was added seven days later for five days to select for stable integrations at the donor locus. Reporter expression (Citrine) was verified by fluorescence microscopy and flow cytometry. HEK293T-LentiX (Takara Bio, 632180, female) cells were used to produce lentivirus for all transductions. Cultures were maintained at 37°C in 5% CO_2_ in DMEM (catalog no.) supplemented with 10% FBS and 1% P/S.

### Pooled delivery of libraries to K562 reporter cells via lentiviral transduction

Large scale lentivirus production and spinfection of K562-JT039 cells were performed in biological duplicate as follows for each library. To generate sufficient lentivirus such that the infection coverage was at least 300X for each library, we plated HEK293T cells in 15 cm tissue culture plates, using 12 plates for the RD-DMS library and 8 plates for the CV-RD and EA-RD libraries (jointly screened). On each plate, approximately 1.5 x 10^7^ HEK293T cells were plated in 30mL of DMEM such that the cultures were about 85-90% confluent 20-24 hours later. Each plate was transfected with 11μg of an equimolar mix of the third-generation packaging plasmids (pMD2.G Addgene plasmid no. 11259, psPAX2 Addgene plasmid no. 12260, and pMDLg/pRRE Addgene plasmid no. 12251), 11μg of purified library plasmid, and 150μL polyethylenimine (PEI) (Fisher Scientific, catalog no. NC1014320) mixed in 2mL Opti-MEM (Thermo Scientific, catalog no. 31985070). All components were mixed and incubated for 20 mins at room temperature before dropwise addition to the HEK293T. Lentivirus was harvested after 48 hrs, 30mL new complete DMEM was added to each plate, and a second harvest was performed 24 hrs after the first harvest. The lentivirus was then pooled and filtered through 500mL 0.45μm PES filtration units (ThermoFisher, catalog no. 166-0045) to remove any cellular debris. Polybrene (PBR) (Fisher Scientific catalog no. TR1003G) was added to the filtered viral supernatant to a final concentration of 8 µg/mL to improve transduction efficiency.

K562-JT039 reporter cells were transduced with the lentiviral libraries by spinfection for 2 hours at 33°C at 1000 x g. 3.9 x 10^6^ K562-JT039 were plated in 2.5mL of lentivirus per well of a 6-well plate. A total of 3.73 x 10^8^ cells were spinfected for each biological replicate of the RD-DMS library, and a total of 1.03 x 10^8^ were used for each replicate of the RD-Predict library. Infected cells were then recovered and grown in complete RPMI for two days and subsequently selected with 10μg/mL blasticidin S HCl (Thermo Scientific, A113903) for 9-11 days. Infection and selection efficiency were monitored daily using flow cytometry to measure the mCherry in the library backbones (Thermo Fisher Attune NxT). From these measurements, we estimate that we achieved greater than 700X coverage per biological replicate for the RD-DMS library, and greater than 3500X for the CV-RD and EA-RD libraries, with MOIs between 0.2-0.4. After selection, antibiotic was spun out of the cultures, and recruitment was induced by treating the cells with 1000 ng/mL doxycycline for five days. Cultures were maintained in 1L or 3L spinner flasks in log growth conditions by diluting the cell concentrations back to 3-6 x 10^6^ live cells per mL every 1-2 days, for the duration of the screens. We aimed to have 3000-20,000X maintenance coverage per biological replicate.

### Magnetic separation

Doxycycline (Fisher Scientific, catalog no. 409050) was added at a concentration of 1000 ng/μL to induce recruitment. Population-wide silencing was assessed daily by flow cytometry measurements of Citrine median fluorescence intensity (MFI). After five days of recruitment by doxycycline, cells were spun down in 500mL falcon tubes at 1000xg for 10 min and the remaining media was aspirated. Cells were washed and resuspended in the same volume of Dulbecco’s Phosphate Buffered Saline (DPBS) (VWR, 21-030-CM) twice to remove any remaining IgG from the serum. 60uL of Dynabeads M-280 Protein G (ThermoFisher, catalog no. 10003D) were prepared per 10 million cells. Beads were resuspended by vortexing for 30 sec, then washed with 1mL Blocking Buffer (2% BSA, 2mM EDTA, pH 8.0 in DPBS) per 200uL bead suspension. The bead mixtures were vortexed and allowed to equilibrate in the Blocking Buffer for 1 min before separation on a magnetic column and removal of the supernatant. Beads were then resuspended in Blocking Buffer and added to the cell pellet at a concentration of 1mL beads per 100 million cells. Magnetic separation was then performed as previously described in ref. ^19^. Briefly, cells were incubated with the beads on a rocker for 60-75 mins at room temperature. Then the mixtures were placed on a magnetic stand for three minutes and the Unbound supernatant was saved as the “OFF” population. The beads were then resuspended in the same volume of Blocking Buffer, rocked for 10 mins, and then reseparated on the column. The wash off was removed and the remaining cell-bound beads were saved as the Bound or “ON” population. Cell populations were stored as pellets at −20°C until extraction of genomic DNA.

### Library preparation and sequencing

Genomic DNA was extracted using the Qiagen Blood Maxi Kits following the manufacturer’s instructions with up to 1×10^8^ cells per column. Elution of DNA was performed with Elution Buffer (EB) and not AE buffer to avoid inhibition of subsequent PCR steps. The domain sequences were then amplified from gDNA using the NEBNext High-Fidelity 2X PCR Master Mix with primers containing Illumina adapters as extensions (**Supplementary Table 3**) in a clean hood. Single test PCRs were performed varying the reaction volume (25-100µL), stock primer concentration (10-100µM), and cycle number (23-32X cycles) until a condition was found that resulted in a clean, single amplicon at the lowest possible cycle number. The optimized conditions were 25 µL reactions with 0.125 µL of each 20µM primer, 12.5 µL of NEBNext High-Fidelity 2X Master Mix, and 2.5 µg gDNA with nuclease free dH_2_O up to the reaction volume. The thermocycling protocol was 3 mins at 98°C, then cycles of 98°C for 10s, 63°C for 30s, 72°C for 30s, and then a final extension at 72°C for 2 mins. For the RD-DMS libraries, 768 x 25µL reactions were performed with Unbound samples to maintain 1000X coverage, and 384 x 25µL reactions were performed with Bound samples to maintain 500X coverage, each with 25X cycles. For RD-Predict libraries, 80 x 25µL reactions were performed for all samples to maintain 1000X coverage with 23X cycles for Unbound libraries and 26X cycles for Bound libraries. For plasmid libraries, around 250 ng of plasmid pool was amplified in 2 x 25µL reactions with 8-9X cycles. All subsequent steps were performed outside the PCR hood. All PCR reactions per sample were pooled in 15mL falcon tubes and vortexed. Then 100 µL were run per sample on a 2µM TBE gel and the library bands at around 420 bp were extracted and purified following the QIAquick Gel Extraction Quick-Start Protocol using MinElute columns (Qiagen, catalog no. 28006). These libraries were then quantified using a Qubit dsDNA HS Assay Kit (Thermo Scientific, catalog no. Q32854) and sequenced on an Illumina NextSeq 550 following the NextSeq System Denature and Dilute Libraries Guide (Illumina Document #15048776) Protocol A. 20% PhiX Control v3 (Illumina, catalog no. FC-110-3001) was loaded with the denatured libraries and paired-end reads (1×200, 1×100) were collected.

### Computing enrichments and hit thresholds

Sequencing reads were demultiplexed using bcl2fastq (Illumina). All subsequent processing steps were performed using the HT-Recruit Analyze package (<u>Github</u>). A Bowtie reference (v.1.2.3) was generated using the library oligo sequences with the script ‘makeIndices.py’. The i5 read length used for alignment to the index and number of allowed mismatches were varied and tested with a subset of 50,000 reads to find a condition that would yield the highest percent of aligned reads and lowest percent of mismatched reads (**Supplementary Fig. 1d**). Conditions using 100bp of the i5 reads with 0 mismatch allowance were selected and used for ‘makeIndices.py’. The enrichments for each domain were computed as Log_2_(OFF:ON) values using the ‘makeRhos.py’ script for each paired Unbound and Bound sample. Domains with fewer than five reads in both Unbound and Bound samples for a given replicate were assigned 0 counts and filtered out from that replicate. Domains with fewer than five reads in one of either the Unbound or Bound sample had those reads adjusted to five to avoid the inflation of enrichment values from low sequenced depth. For all screens, the hit thresholds were set based on the distribution of score of negative controls, calculated as two standard deviations away from the mean of the negative controls (**x̄ + 2σ**). The thresholds varied between screens depending on coverage, sequencing depth, and reproducibility between replicates.

### Individual rTetR-RD cloning

RDs were ordered as 300bp gBlock gene fragments (Integrated DNA Technologies (IDT)) with Esp3I/BsmBI restriction sites and cloned as fusions with rTetR(SE-G72P) in pJT126 (Addgene #161926). gBlocks were resuspended to 10-20 ng/μL in EB. pJT126 (Addgene no. 161926) was predigested with Esp3I (NEB, catalog no. R0734L) for 15 mins at 37°C and then purified by gel extraction. 10μL Golden Gate reactions were performed using a 2:1 molar ratio of insert to digested backbone with 75 ng pJT126 and 5.3 ng of each 300bp insert, with 1 μL T4 DNA Ligase Reaction Buffer (NEB, catalog no. B0202S), 0.125 μL T4 DNA Ligase (NEB, catalog no. M0202T), 0.75 μL Esp3I (NEB, catalog no. R0734L), and nuclease free dH_2_O up to the final volume. Reactions underwent 30 cycles of digestion at 37°C at ligation at 16°C for 5 min each, followed by a final 5 min digestion at 37°C for and then 15 min heat inactivation at 70°C. 2 μL of each Golden Gate reaction were transformed into 15 μL Endura cells (Lucigen, catalog no. 60242-2) and single colonies were recovered from overnight incubation on Carbenicillin agar plates. Correct plasmid clones were confirmed by whole-plasmid sequencing (Primordium) and plasmids were purified following the QIAprep Miniprep Handbook using MinElute columns (Qiagen, catalog no. 28006).

Individual point mutants were cloned following the Improved Methods for Site-directed Mutagenesis using NEBuilder HIFi DNA Assembly Master Mix (NEB). Briefly, two PCR products created with complimentary flanking primers were amplified from the corresponding WT RD vector to introduce the desired mutation and facilitate insertion between the Psil-v2 and SbfI-HF digestion sites in pJT126 upon relegation. To amplify the fragments, PCR reactions were performed with 5μL each of 5X Q5 Reaction Buffer and 5X Q5 High GC Enhancer (NEB, catalog no. B9027S), 0.5μL 10 mM dNTPs, 0.125μL of each 100 μM primer (IDT), 5ng of template vector, 0.25μL Q5 High-Fidelity DNA Polymerase (NEB, catalog no. M0491L), and nuclease free dH_2_O up to 25μL. Amplification was performed with 98°C for 30 sec, followed by 35 cycles of 98°C for 10 sec, 64°C for 30 sec, and 72°C for 30 sec, then a final extension at 72°C for 2 min. pJT126 was digested with 1μL each of PsiI-v2 (NEB, catalog no. R0744S) and SbfI-HF (NEB, catalog no. R3642S), 5μL 10X rCutSmart Buffer (NEB, catalog no. B6004S), 2μg pJT126 template, and nuclease free dH_2_O up to 50μL, then purified by gel extraction. 10μL Gibson Assembly reactions were set up with 0.01pmol each of PCR fragment and digested backbone, 5μL of 2X NEBuilder HiFi DNA Assembly Mix (NEB, catalog no. E2621S), and nuclease free dH_2_O up to total volume. Products were transformed, confirmed, and purified as described above.

### Individual recruitment assays and flow cytometry measurements

Single rTetR-RD constructs were introduced into K562-pJT038 reporter cells via lentivirus transduction at an MOI of 0.1-0.4 (estimated from mCherry+). 2-3 days post-transduction, cells expressing the RDs were selected with 10μg/mL blasticidin S HCl (Thermo Scientific, A113903) for 8-10 days, and the active reporter was subjected to 1μg/mL puromycin (Cayman Chemical no. 28240) selection during expansion. The cells were then removed from all selection, washed with DPBS, and resuspended in complete RPMI with or without 1000 ng/μL doxycycline for up to 7 days for recruited and non-recruited conditions, respectively. Flow cytometry measurements were taken daily to monitor RD (mCherry) and reporter expression (Citrine) (Thermo Fisher Attune NxT). 10,000-20,000 live, single cell events were collected per sample, per timepoint.

### Flow cytometry analysis

Data were analyzed using Cytoflow (v.1.2, <u>Github</u>) and custom Python scripts. For all analyses, events were first gated for viability using FSC-A vs. SSC-A and only “live” events were considered. Citrine and mCherry gates and compensation matrixes were set using double-negative K562 or mCherry-negative K562-pJT039 cells. Events were considered ON or positive if they exceeded these thresholds.

### Western blots

One million live cells were pelleted and supernatant was aspirated. Cell pellets were resuspended in 50uL of pre-chilled lysis buffer (1X RIPA, 1X Halt Protease and Phosphatase Inhibitor Cocktail) and incubated on ice for 30 mins. Lysates were spun down at 10,000xg for 10 minutes at 4C and the lysed supernatant was transferred to a fresh pre-chilled 1.5uL Eppendorf. Protein amounts were quantified using the BCA assay against BSA standards. The samples were then denatured by the addition of 1X laemmli loading buffer (LDS, DTT) followed by a 5 min incubation at 90C. 20ug per sample were subsequently loaded onto a NuPAGE 4-12% Mini Protein Gel and run for 2 hours at a constant voltage of 100V. The gel was then transferred to a PVDF membrane for 35 mins at 120mA in a BioRad semi-dry rig. The membrane was first blocked with 7% nonfat dry milk (Bio-rad, catalog no. 1706404) in PBS for 1 hour at room temperature, then probed using anti-FLAG M2 monoclonal antibody (1:1000, mouse, Sigma-Aldrich, catalog no. F1804) and anti-Histone H3 polyclonal antibody (1:2000, rabbit, Abcam, catalog no. ab1791) with gentle rotation at 4C overnight. Next, the membrane was washed with TBST 3×5 min before being probed with goat anti-mouse IgG IRDye 680RD (1:20,000, LICOR Biosciences, catalog no. 926-68070) and goat anti-rabbit IgG IRDye 800CW (1:20,000, LICOR Biosciences, catalog no. 926-32211) secondary antibodies for 1 hour at room temperature.

### Clinical annotations

All missense variants were queried from both ClinVar and gnomAD 4.0 for each gene that contained a tile in the RD-DMS library. Note that no missense variants existed in the ClinVar database for ASCL5, CBX5, RYBP, or CDY2A (Accessed December 28th 2023). ClinVar entries labeled “Pathogenic”, "Pathogenic/Likely pathogenic", or “Likely pathogenic" were classified “Pathogenic”. Variants of unknown significance (VUS), or those labeled with “Conflicting interpretations of pathogenicity” or “Uncertain significance” were classified as “Uncertain”. Those labeled “Benign”, “Benign/Likely benign”, and “Likely Benign” were classified as “Benign”. All variants found in gnomAD 4.0 were classified as “Benign” unless otherwise annotated with a clinical significance in the aforementioned “Pathogenic” or “Uncertain” categories.

### Functional annotation of clinical variants

To determine if an “Uncertain” or “Pathogenic” variant has significantly altered repressor activity, we considered the measured values of each variant compared to WT values. The mean and standard deviation of all WT Log_2_(OFF:ON) values per domain were calculated and thresholds were set 3 standard deviations above and below the WT mean. If an individual variant had a Log_2_(OFF:ON) value exceeding either threshold, it was reported to have significantly altered repressor activity in our screen.

For the vignette on the TWIST1 HLH RD, missense mutants that were within residues [109-121] were considered part of the basic subdomain whereas those within [122-165] were considered part of the HLH subdomain. Welch’s t-tests were performed testing if the Log_2_(OFF:ON) values for “Benign” variants significantly differed from those of either “Uncertain” or “Pathogenic” variants, between variants within the same subdomain. P-values < 0.05 were considered to indicate significant population differences. P-values < 0.01 are represented by a double asterisk and < 0.001 by a triple asterisk.

### RNA-seq analysis

Raw fastq reads from K562 were retrieved from ENCODE (SRA:SRR3192408) using sratool. Adaptors were trimmed using Trim Galore then pseudoaligned via kallisto and identified on GRCh38. List of all TFs annotated as HLH family members was obtained from ^55^ and the Fragments Per Kilobase of transcript per Million mapped reads (FPKM) for each gene was calculated in R. Gene were classified as having “high”, “medium”, “low”, or “no” expression if their FPKM values were >1000, >10, >1, and <1, respectively.

### Functional SLiM finder

To systematically identify short linear motifs (SLiMs) that are potentially critical to biological function, our analysis was restricted to domains or domain regions exclusively categorized or predicted to be disordered. SLiMs within these disordered sequences were identified by contrasting wild-type (WT) sequences with their variant counterparts, focusing on regions with 4 or 8 contiguous mutated residues. From these regions, candidate SLiMs were proposed based on their potential to disrupt or create functional sites. The difference between the median Log_2_(OFF:ON) values of the WT sequences and mutants was quantified as a measure of the functional impact of each motif. Motifs exhibiting the highest MAE values above zero were considered to have the largest biological impact, reflecting significant functional alterations. Any SLiMs corresponding to mutants with a log repression value greater than −0.5 were excluded. Subsequently, we determined if any identified SLiMs overlapped with others within the same domain, and we reported both the individual and overlapping SLiMs as separate entries. To extract a regex pattern from a corresponding SLiM, we assessed the importance of each amino acid by comparing its corresponding importance to our single mutational data. The importance of single amino acids was determined by averaging the log-repression scores across all 19 single amino acid substitutions. Unless the averaged score for a given amino acid was at least 0.5 less than the WT log repression value, a wildcard character ‘*’ was assigned in the regex pattern. Additionally, we reported only those SLiMs that contained at least three characters besides the wildcard character and removed any leading and trailing wildcard characters. Lastly, to quantify any novel SLiMs, we compare identified patterns against the Eukaryotic Linear Motif (ELM) resource. A SLiM is reported as novel if no regex matches are found within the database. In **Fig. 4j**, we only provide annotations from ELM corresponding to the nucleus cell compartment. Additionally, in **Supplementary Table 6**, ELM annotations for both the nucleus cell compartment and across the full ELM database are provided. Background motifs in **Fig. 4j** are derived from regions of low functional importance within the corresponding domains. These motifs are calculated by averaging the importance scores of the 10 least significant regions.

### Multiple-Sequence alignments of evolved sequences

To perform multiple sequence alignments of our domains, the sequences were first parsed into a FASTA file, then input into the Multiple Sequence Alignment with High Accuracy and High Throughput (MUSCLE) algorithm via the command line program *muscle*. The resulting MSAs were visualized using the pyMSAviz package.

### Homology Searches

To study the conservation of our target protein sequence, we first retrieved homologous sequences using the Basic Local Alignment Search Tool (BLAST). The BLAST search was conducted using the Biopython library’s NCBIWWW.qblast function, which facilitates online querying of the NCBI databases. The query was performed using the BLASTP program against the non-redundant protein sequences (nr) database. Search results were filtered for txid9606 (“homo sapiens”). Following the BLAST search, the resulting sequences were parsed and written into a FASTA file using the NCBIXML.read and SeqIO.write functions, respectively, from the Biopython library.

### Sequence conservation calculation

To study the conservation of our target sequences, we first retrieved homologous sequences as described above, without a filter for “homo sapiens’’. This collection of sequences represented homologous proteins from a wide range of organisms. Next, we performed multiple sequence alignment (MSA) on the collection of homologous sequences using Clustal Omega. The Clustal Omega command line tool was accessed using Biopython’s ClustalOmegaCommandline function. The aligned sequences were written into a new FASTA file. To calculate the conservation score for each position, we extracted the column of amino acids from the alignment and counted the occurrences of each amino acid. The conservation score was calculated as the fraction of sequences with the most common amino acid at that position.

### Neural Network

The proposed neural network model aims to tackle the challenges associated with capturing sequence and structural homology of proteins. As proteins are complex macromolecules that serve a broad range of functions, capturing both their sequence and structural features is crucial for understanding their function, interactions, and evolutionary relationships.

#### Modeling repression strength

Traditional count-enrichment modeling typically relies on calculating the Log_2_ ratio of the OFF to ON states to estimate repression strength, and then employs mean squared error (MSE) to train standard regression models. In contrast, we fit models directly to the measured sequence read counts in OFF and ON populations using a binomial loss function, accounting for higher variance at lower read counts ^124^. Models predict the probability of each sequence in the OFF (*P*_*off*_) and ON populations (*P*_*on*_ = 1 − *P*_*off*_).

#### Custom Loss Function

Models were fit directly to the measured sequence read counts in OFF and ON populations using a binomial loss function, accounting for higher variance at lower read counts.It is defined by the following formula:

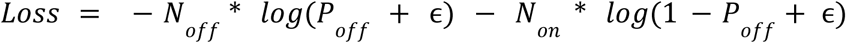

where epsilon is a small constant to avoid numerical instability. For calculating the loss function, the final *N*_*off*_ and *N*_*on*_ counts are derived using a weighted average of counts from both populations across the two biological replicates.

The model is trained using the Adam optimizer with a configurable learning rate. A learning rate scheduler adjusts the rate based on the plateau of the validation loss. Early stopping is implemented, based on a patience parameter, to prevent overfitting. The model is trained on a shuffled dataset and validated on a separate dataset. The maximum number of training epochs is set to 100, with early stopping if validation loss does not improve.

We assessed performance based on the concordance of measured and predicted *P*_*off*_ for held-out sequences not used in training where the measured *P*_*off*_ = *N*_*off*_/(*N*_*on*_ + *N*_*off*_)

#### Sequence Encoder (ESM Embedding)

The sequence encoder is specifically designed to capture sequence homology. It operates on ESM embeddings with an input size of 1280. A Gated Recurrent Unit (GRU) network is employed, with 256 hidden units across 2 layers. This GRU is bidirectional to account for both forward and backward sequence information. The last hidden states from both directions are concatenated and followed by batch normalization and dropout layers for regularization.

#### AA Descriptor Encoder

To supplement the sequence homology information, a separate GRU-based sequence encoder operates on AA descriptor features with an input size of 66. Similar to the ESM embedding encoder, this component employs a bidirectional GRU with 256 hidden units across 2 layers. The output hidden states from both directions are concatenated and passed through batch normalization and dropout layers for regularization.

#### Contact Map Encoder

The contact map encoder is aimed at capturing structural homology. It uses a Convolutional Neural Network (CNN) architecture that starts with a single channel contact map, which is passed through two convolutional layers. Each convolutional layer is followed by batch normalization and dropout layers. The first and second convolutional layers both have 32 output channels and utilize 3×3 kernels with stride and padding set to 1. Max-pooling operations are applied after each convolutional layer to reduce dimensionality.

#### Fully Connected Layer for Prediction

The outputs from the ESM sequence encoder, AA descriptor encoder, and the contact map encoder are concatenated to form a pooled feature vector. This concatenated vector is fed into a fully connected layer with 512 units, followed by batch normalization and dropout layers for regularization. Finally, a single output unit is applied with a sigmoid activation function to map the output to a range between 0 and 1.

#### Feature Generation

1. ESM Embedding: We used the Evolutionary Scale Modeling (ESM-2), specifically the esm2_t33_650M_UR50D pretrained model, to generate embeddings for amino acid sequences. This model has 650 million parameters and is designed to be especially effective at capturing contextual and evolutionary information in protein sequences. 2) Amino Acid Descriptors: Complementary to ESM embeddings, we also included 66 physicochemical descriptors for each amino acid in the sequence. These descriptors were obtained from a precompiled data frame and include properties such as hydrophobicity, charge, and size. 3) To encapsulate the structural homology of proteins, we leveraged contact maps generated from ESM-Fold structures of mutants. Specifically, we generate the C-alpha distances between amino acid residues to generate a distance matrix. This matrix has dimensions of 80×80 and serves as an additional feature that provides insights into the spatial relationships between amino acids in the folded protein structure. The C-alpha distance matrices were normalized to ensure scale invariance and better convergence during the training process. These normalized features were then fed into the corresponding encoders in our neural network architecture for further processing.

#### Model Training

In this study, we utilized PyTorch ^125^ for the implementation and training of our model on GPUs to accelerate the training process. The optimization of the model was carried out using the Adam optimizer with an initial learning rate of 0.001. To further enhance the model’s performance, a ReduceLROnPlateau learning rate scheduler was employed, dynamically adjusting the learning rate based on the validation loss with a patience of 5 epochs and a reduction factor of 0.5. The training regimen was designed to span 100 epochs, incorporating an early stopping mechanism to prevent overfitting. This mechanism halted the training if no improvement in validation loss was observed for 30 consecutive epochs, ensuring the model’s generalization to unseen data by saving the weights of the best-performing model during the training process. As depicted in **Fig. 4b**, we trained 6 replicates for each held-out (test set) domain (x-axis), presenting the average performance across these 6 runs. For all runs, the validation set comprised RYBP [121-201], CBX5 [111-191], and PCGF2 [1-81]. These domains were chosen because they are variants of the original set, with minor alterations involving slight displacements of 1-2 amino acids. Notably, these domains were excluded from the training and validation sets for models employed in predicting the analogous domains depicted in **Fig. 4b**.

### 2D projection

We generate 2D projections using two distinct sets of features: (1) with ESM-2 Embeddings, and (2) neural network embeddings derived from TENet. For our ESM Embeddings, we produce positionally averaged descriptors of shape (1280 dimensions), achieved by averaging the features across the 80 amino acids, akin to the strategy employed for deriving features for the ESM Metagenomic Atlas ^37^. In contrast, our neural network features are extracted from the last layer of our neural network, resulting in each amino acid being represented by a 512-dimensional vector. For both sets of features, we use a multi-step process to achieve our 2D projections. Initially, we implement Principal Component Analysis (PCA) ^126^ to reduce the dataset’s dimensionality to 50 principal components. Following PCA, we generate a K-Nearest Neighbors (KNN) Graph ^127^ based on these principal components, using 15 neighbors for each point to establish connectivity while excluding the point itself. Subsequently, we apply Uniform Manifold Approximation and Projection (UMAP) ^128^ with specific settings (15 neighbors, a minimum distance of 0.3, and Euclidean metric) to the PCA-reduced data to generate a 2D representation, effectively preserving the structural relationships among data points. Finally, we run the Leiden algorithm ^129^ on the KNN Graph to optimize clustering for data interpretation. We note that the UMAP algorithm used in our analysis is a stochastic method, meaning that it can produce slightly different results each time it is run on the same dataset. This randomness is inherent to the algorithm and may lead to minor variations in the visualizations and results obtained.

To evaluate the quality of clustering of sequence variants based on their ESM-2 and TENet embeddings, we calculated two Silhouette scores using two different types of labels for each sequence. The first score tested how well sequences separated based on structural homology, with each sequence variant labeled according to the structural class of the domain (collected via Uniprot annotations). The Silhouette score was computed using the scikit-learn function by comparing intra-cluster cohesion with inter-cluster separation. The second score was based on repression activity, where log-transformed repression values were discretized into 5 quantile-based bins to ensure even representation across the dynamic range of activity. Both Silhouette scores were calculated using the pairwise PCA distance matrix, gated by a K-Nearest Neighbors (KNN) graph constructed from the PCA-reduced data (50 principal components). The KNN graph, which connected each data point to its 15 nearest neighbors based on Euclidean distances, was used to assess clustering for both structural homology and repression activity.

### Evolutionary algorithm

The Evolutionary Algorithm (EA) was initialized using wild-type sequences from the specified domains: CBX4 [109-189], ID2 [19-96], NEUROG2 [100-180], HES3 [101-181], and MECP2 [241-321]. In each iteration, up to four random substitutions were permitted per AA sequence, with modification sites randomly selected. The algorithm maintained a consistent population size of 500 sequences per iteration. From one iteration to the next, only the top 2% of sequences, selected based on their fitness scores, were retained. The remaining 98% were discarded and replaced with new sequences generated through random mutations of the retained 2%. The fitness of an AA sequence was determined by its performance on a fitness function, which varied depending on the task. For maximization tasks, the fitness function was defined as the direct P(OFF) output from our neural network. For tasks aimed at designing domains with specific repression levels, the fitness function was the mean absolute error between the desired P(OFF) value and the value predicted by the neural network. For the different domains, the best corresponding models were used, where the windowed-shuffled reads were shown for training. For each domain, the most suitable models were employed to predict the P(OFF) values. These models were trained using windowed-shuffled reads from the corresponding domains, excluding any single mutational information during the training process.

---

**Supplementary Figure 1:**
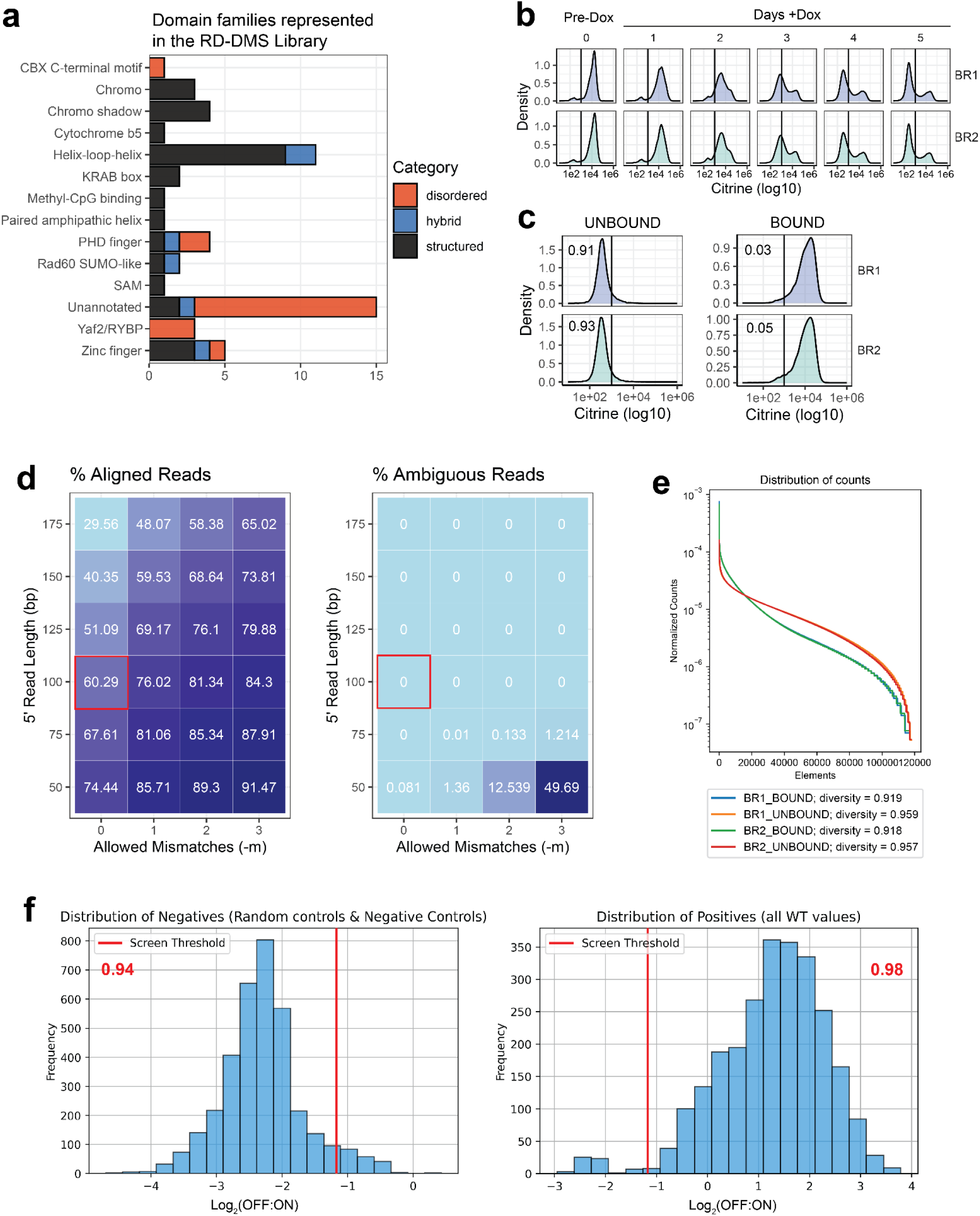
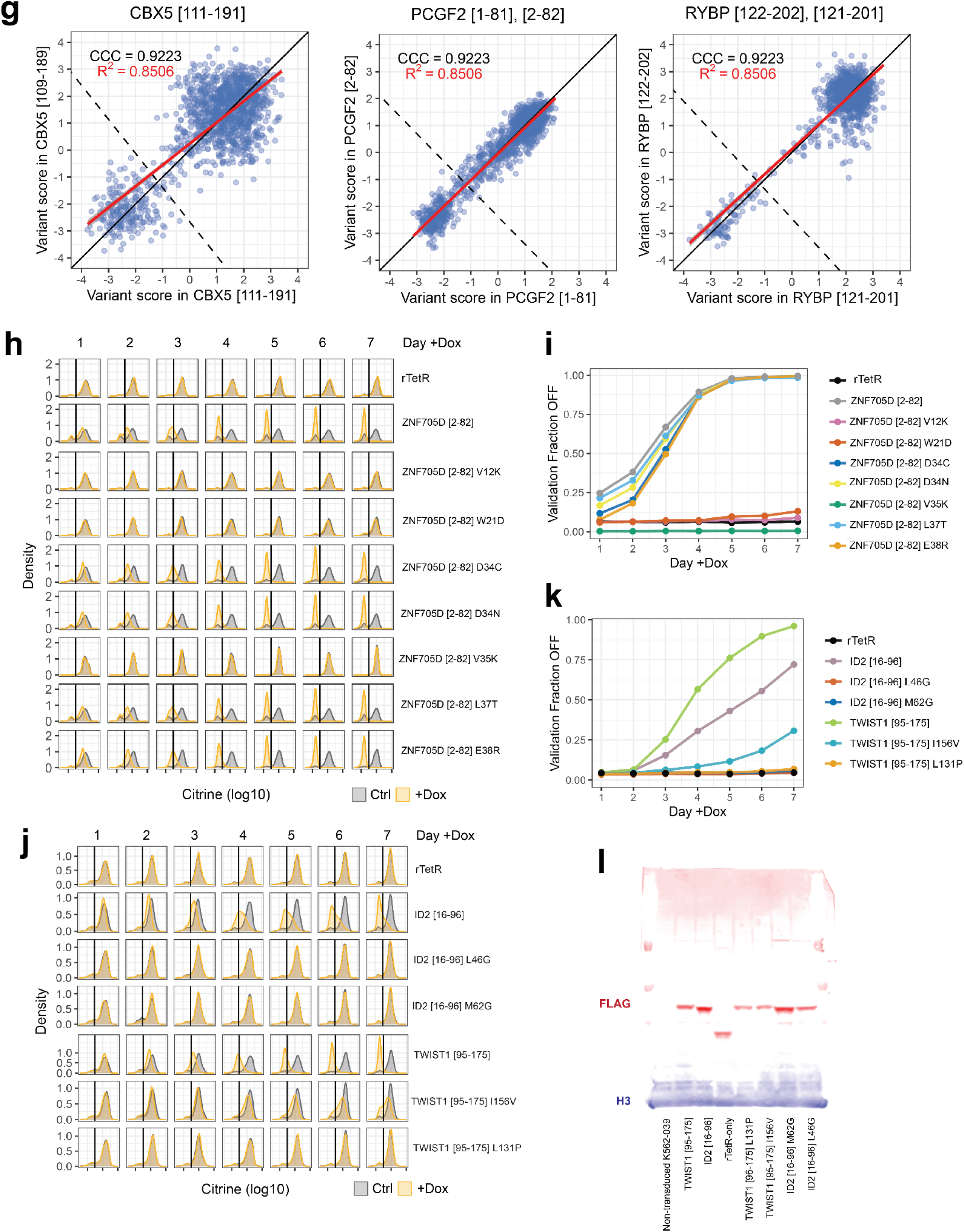
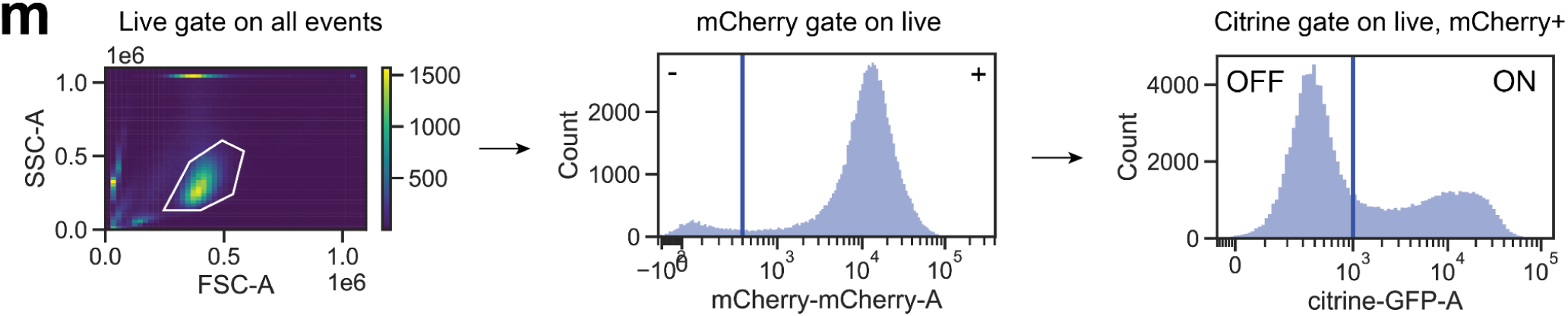
Supporting information for the RD-DMS screen and analysis. a. Distribution of RDs selected for deep mutational scans by protein family (**Methods: Domain classification and disorder prediction**). Fill values indicate number domains predicted to be structured (black), hybrid (blue), or disordered (red). b. Reporter intensity measured by citrine fluorescence of the pool of screened cells before and after the addition of doxycycline for 2 biological replicates, over 5 days. Cells were sampled on day 5 post-dox addition and immediately subjected to magnetic separation. c. Results of magnetic separation of “ON” cells (bead BOUND) and “OFF” cells (UNBOUND) for 2 biological replicates. One billion cells were subjected to magnetic separation per replicate. The fraction of OFF cells are indicated by the number in each plot. d. Alignment results for HT-recruit reads where the number of bp from the forward i5 read and number of allowed mismatches to the index were varied for a pilot subset of 50,000 reads (**Methods: Computing enrichments and hit thresholds**). The % of total reads aligned (left) and % ambiguous reads (right) are shown for each of the indicated conditions. Selected conditions are highlighted by red boxes. e. Diversity of BOUND and UNBOUND populations for both biological replicates of the screen shown as the distribution of counts per library element. f. Density plots showing the distribution of all negative and positive control elements. The repression threshold is shown as a red vertical line and the proportion of elements below (negative) or above (positive) the threshold for respective control types are indicated in red text. g. Scatter plots comparing the Log_2_(OFF:ON) values for homologous mutants belonging to different but overlapping tiles. The concordance correlation coefficient is denoted by CCC and the line with slope = 1 and y-intercept = 0 is drawn in black. The coefficient of determination is denoted by R^2^ and the best fit linear regression is shown in red. h. Density plots showing the distribution of citrine intensity as measured by flow cytometry for cell lines expressing individual RD tiles over time, pre- and post-addition of doxycycline. Results for wild-type and selected mutants of ZNF705D [2-82] are shown. i. Timecourse for individual ZNF705D [2-82] validations. showing the cell fraction OFF as measured by Citrine signal for each mutant vs. days post addition of doxycycline. j. Same as (**h**). Results for wild-types and selected mutants of TWIST1 and ID2 RDs are shown. k. Same as (**i**). Results for individual TWIST1 and ID2 RD validations. l. Expression levels for individual rTetR-RD-FLAG(3X) constructs were measured by Western blot with an anti-FLAG antibody (red). RDs are labeled in their respective lanes. Anti-histone H3 was used as a loading control for normalization (green). Shift from expected molecular weights is likely due to post-translational modifications. m. Gating strategy used for all flow cytometry data presented and analyzed in this manuscript.

**Supplementary Figure 2:**
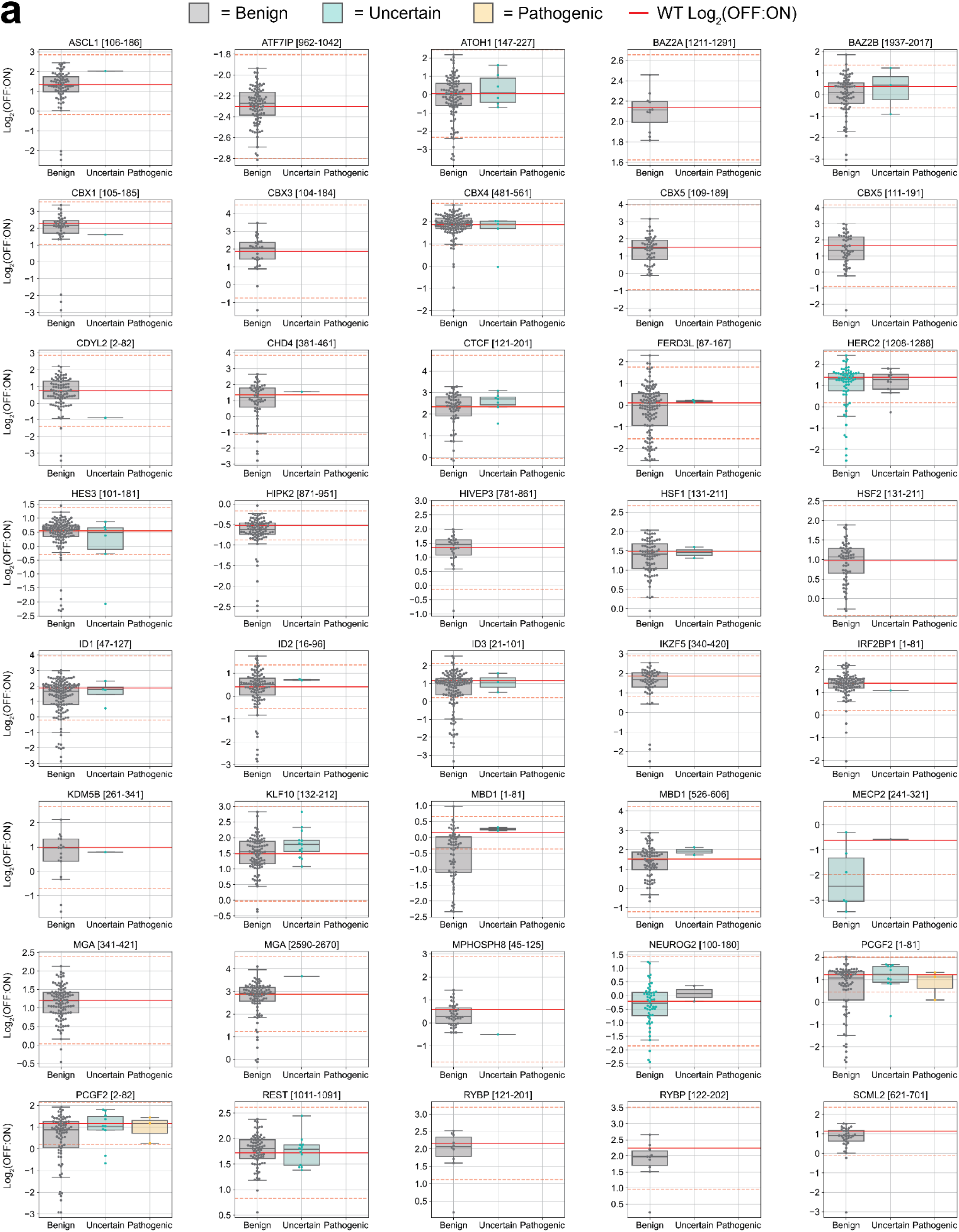
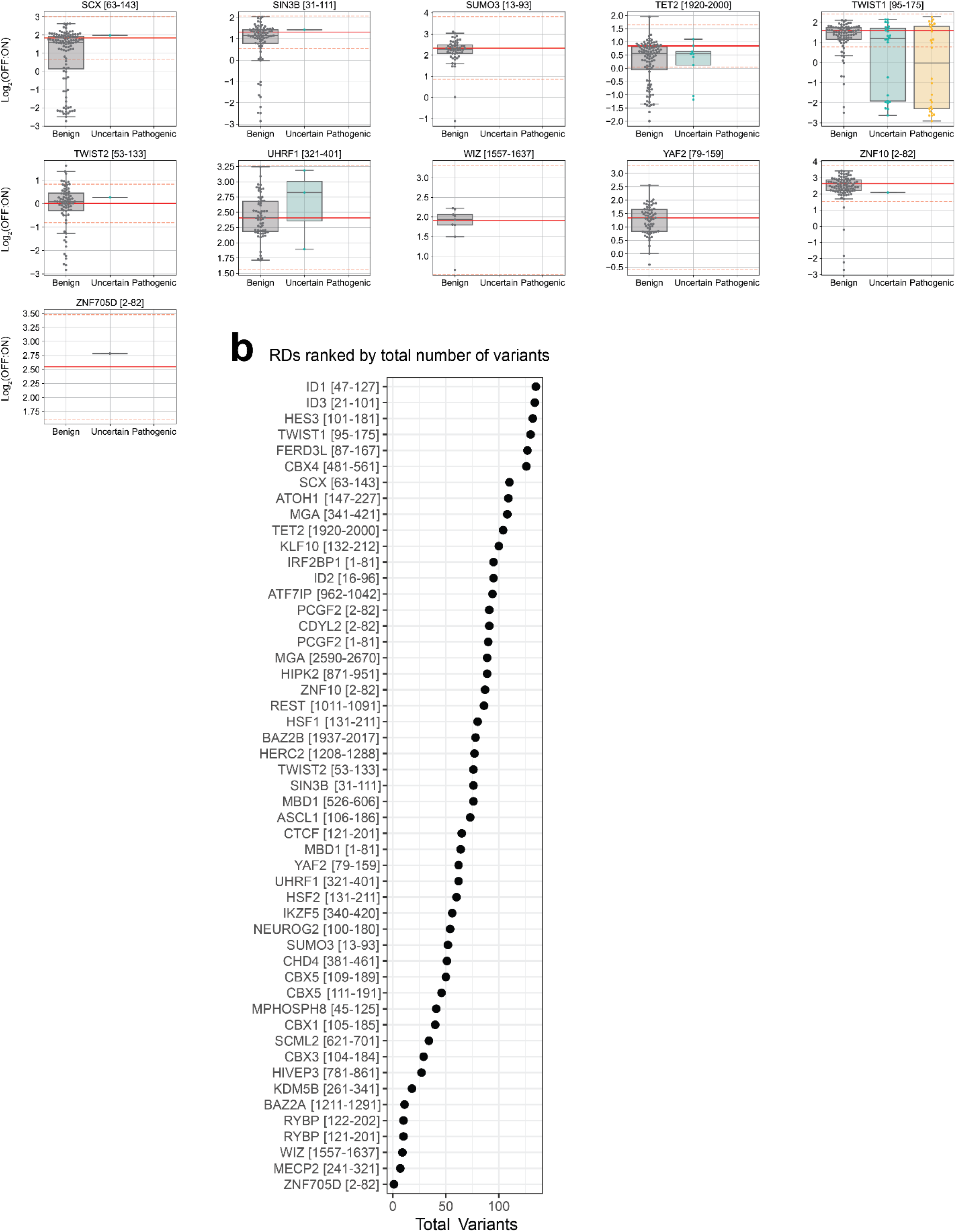
Additional characterization of the repressor activity of clinically-relevant coding variants in the RD-DMS screen. a. Distribution of Log2(OFF:ON) values for all missense variants measured in the RD-DMS screen, per tile. Variants are categorized into healthy (gray), uncertain (green), and pathogenic (yellow) groups based on reported clinical significance. The mean of the WT elements (up to 50) is indicated by a solid red line. Thresholds set three standard deviations above or below the mean of the WT elements are indicated by dashed red lines, and variants above or below these lines were considered to have significantly altered repressor activity. b. Total number of missense variants in all RDs screened in the RD-DMS library.

**Supplementary Figure 3:**
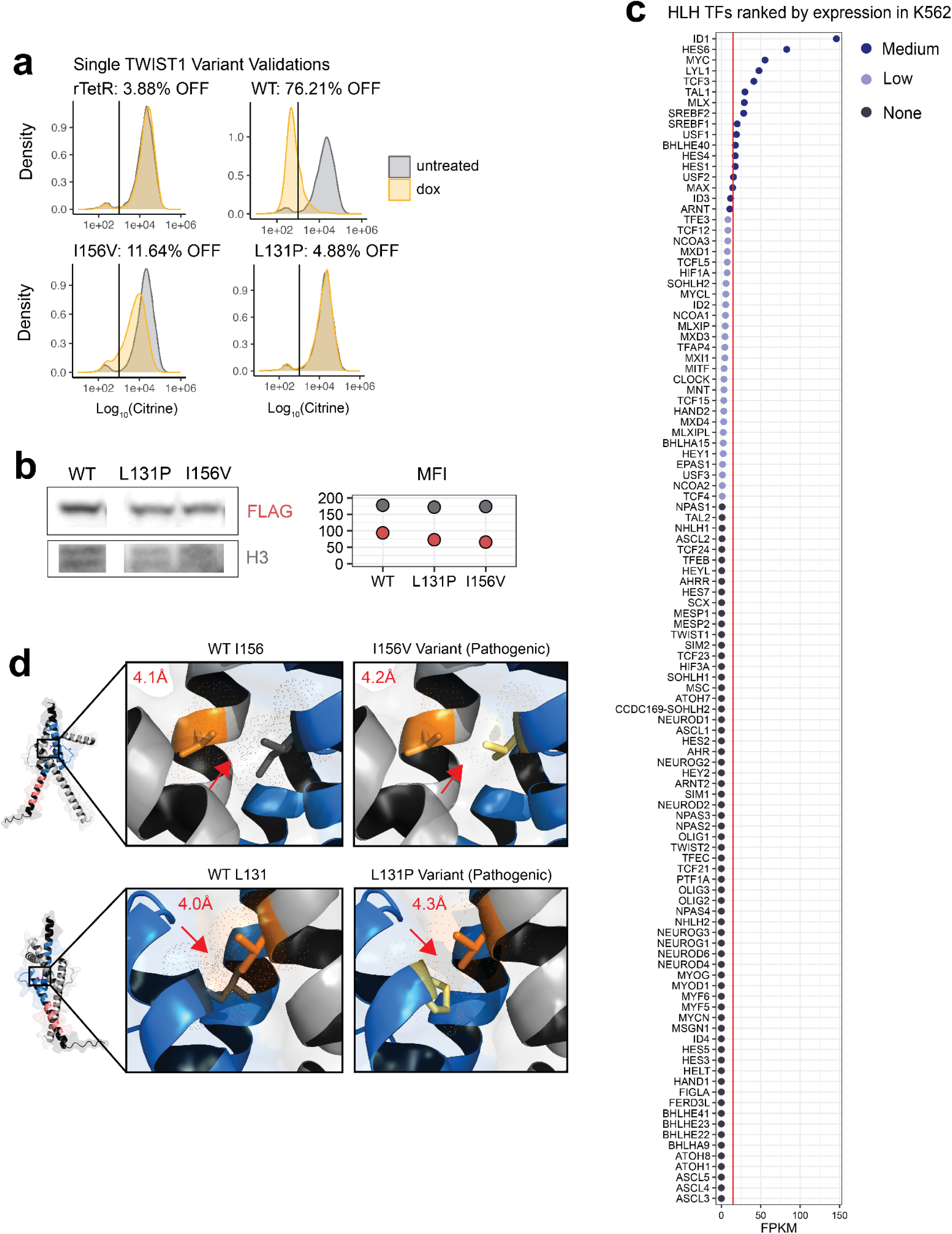
Supporting information for pathogenic LoF missense mutants in the HLH subdomain of TWIST1. a. Single validation data for I156V and L131P variants. Histograms show Citrine reporter signals after 5 days of dox recruitment for respective variants (colored) relative to untreated controls (gray). b. Zoom-in on FLAG Western blot (**Supplementary Fig. 1l**) measuring the stability of indicated TWIST1 bHLH recombinant variants. Median fluorescence intensity (MFI) was quantified for FLAG and H3 signal (left). c. Fragments Per Kilobase of transcript per Million mapped reads (FPKM) for all HLH family members in K562 cells. Genes are ranked from highest to lowest FPKM and are colored based on expression. The mean FPKM for all genes with non-zero counts is indicated in red. d. Location of highlighted pathogenic variants on the TWIST1 structure in complex with TCF3. Both I156V (top) and L131P (bottom) are located at the dimerization interface. Red arrows indicate the approximate location of the reported molecular distances measured between the indicated TWIST1 residue (dark gray or yellow) and the closest intermolecular residue on TCF3 (orange).

**Supplementary Figure 4:**
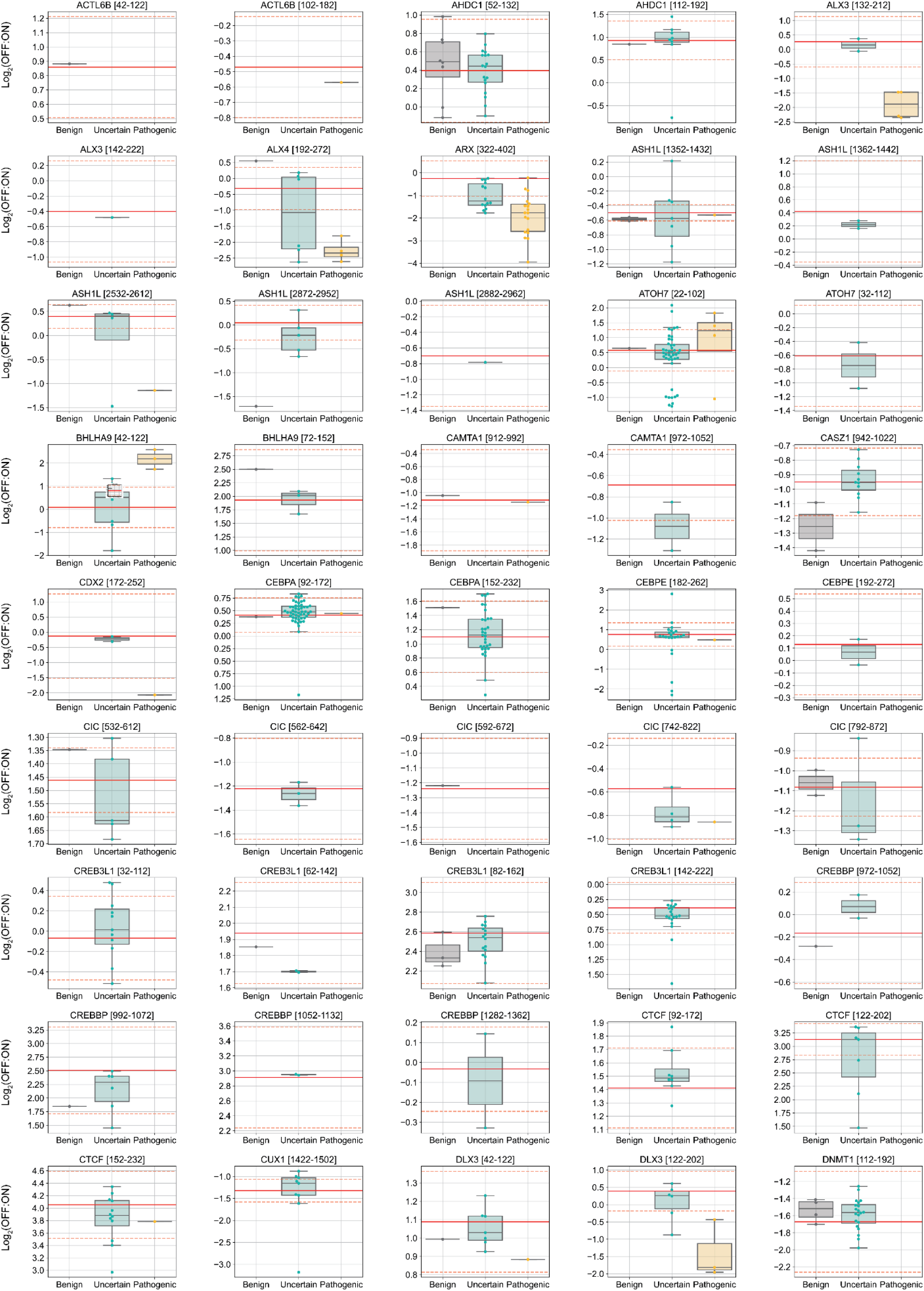
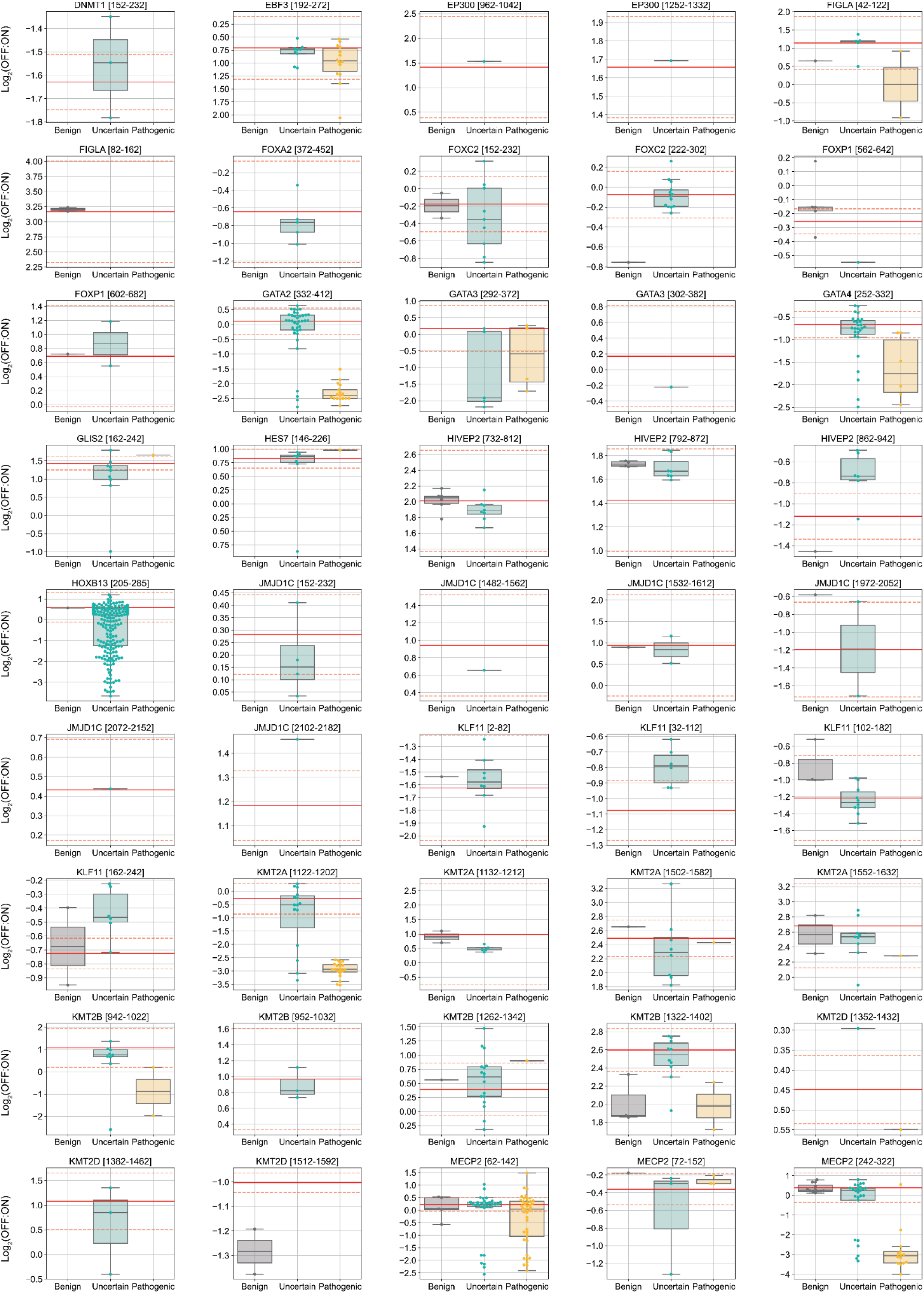
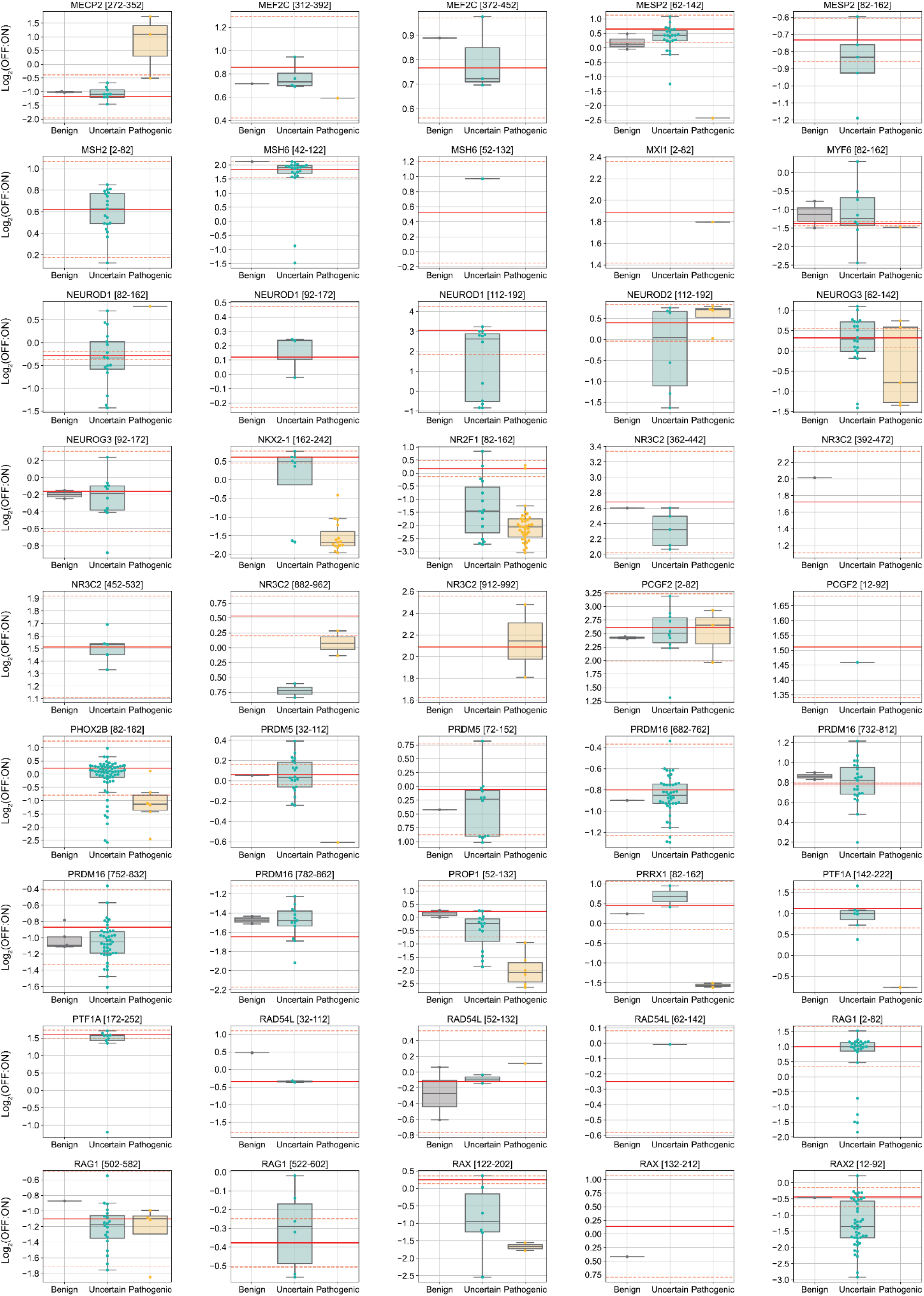
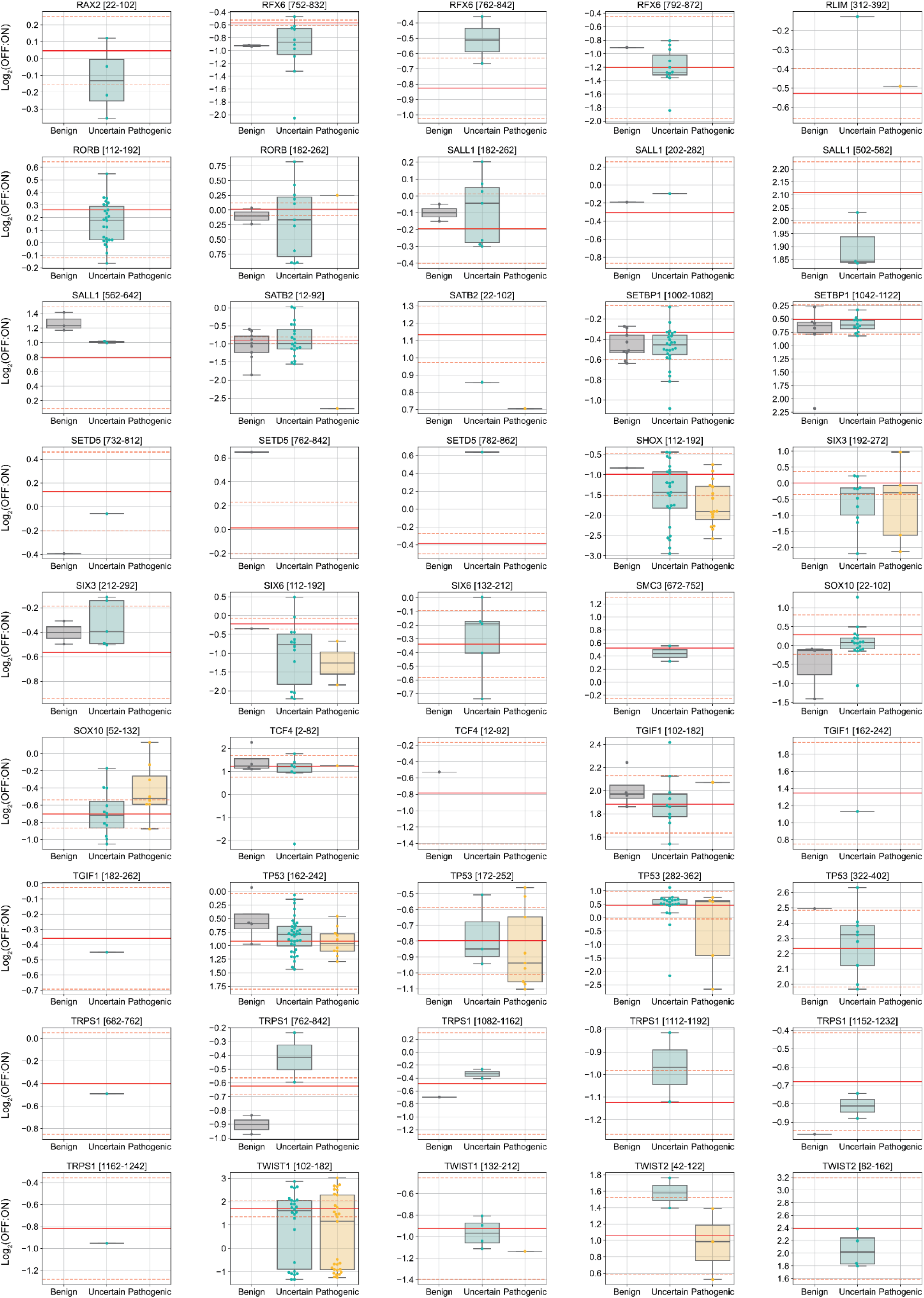
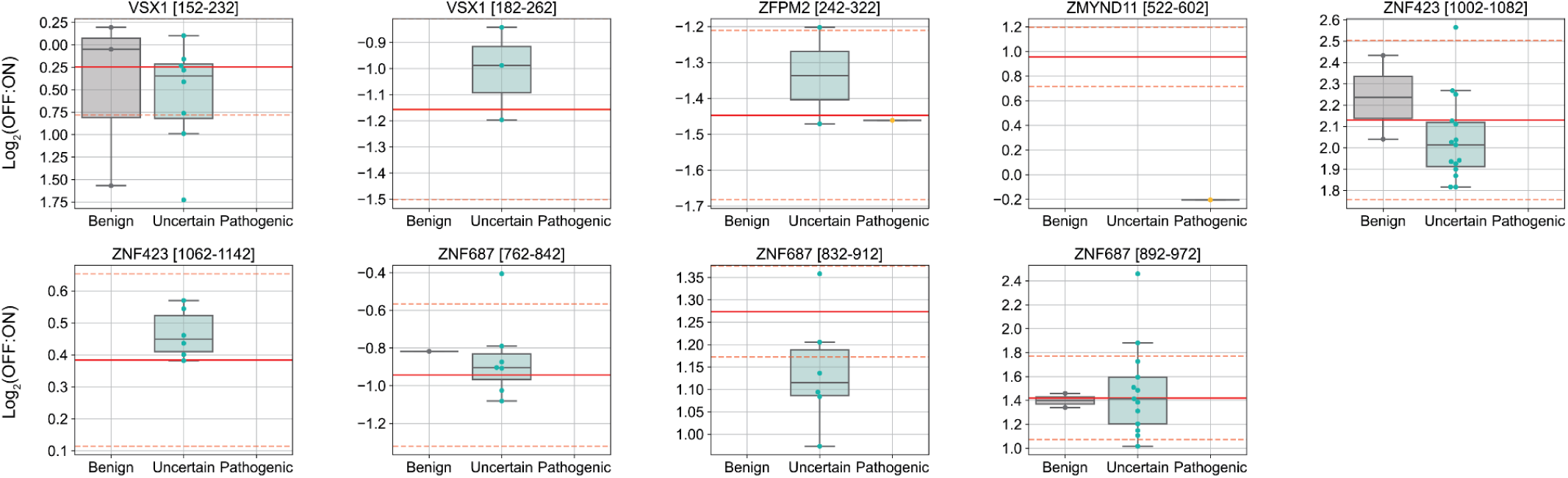
Characterization of the repressor activity of clinically-relevant coding variants in the CV-RD screen Distribution of Log_2_(OFF:ON) values for all missense mutants measured in the CV-RD screen, per tile. Variants are categorized into healthy (gray), uncertain (green), and pathogenic (yellow) groups based on reported clinical significance. WT means and standard deviations are set with up to 3 WT elements per tile and are indicated by a solid and dashed red lines, respectively. Thresholds set three standard deviations above or below the mean of the WT elements are indicated by dashed red lines, and variants above or below these lines were considered to have significantly altered repressor activity.

**Supplementary Figure 5:**
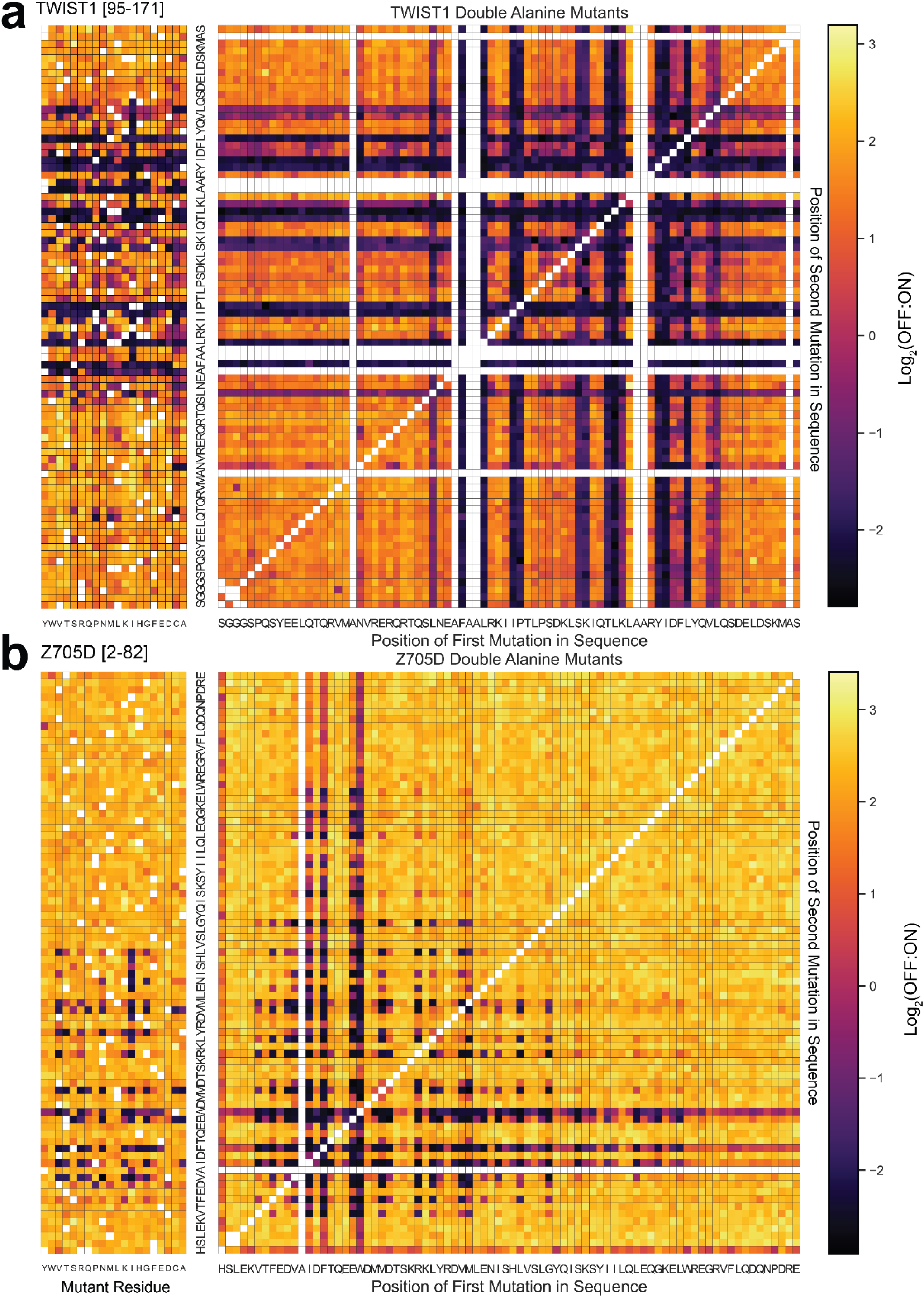
a. Heatmaps displaying Log2(OFF:ON) repression values for single substitutions (left) for all double Alanine substitutions in TWIST1 [95-171] (right). b. Heatmaps displaying Log2(OFF:ON) repression values for single substitutions (left) for all double Alanine substitutions in Z705D [2-82] (right).

**Supplementary Figure 6:**
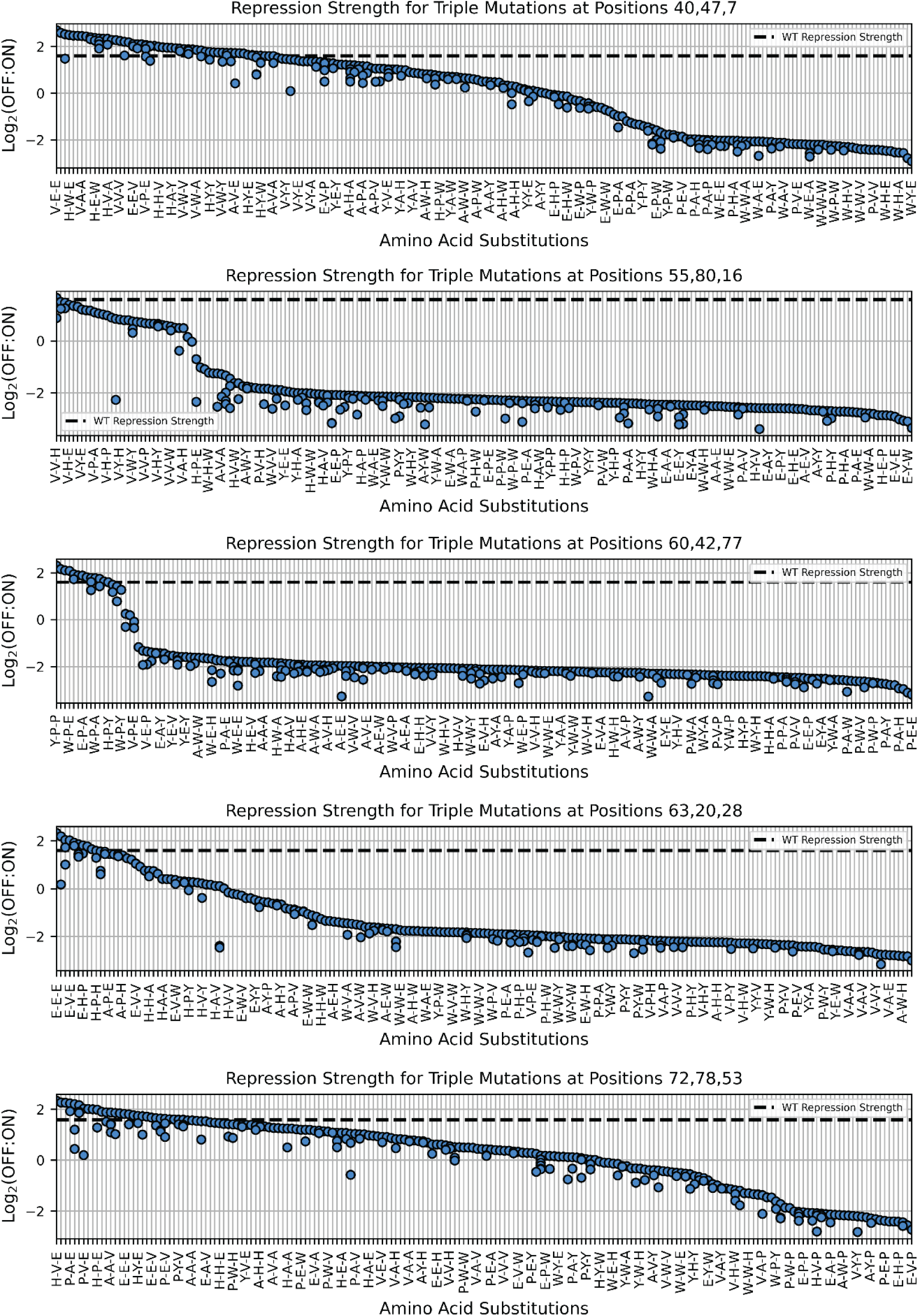
Repression strengths of selected TWIST1 [95-185] triple mutants Five representative samples of triple mutational data at specific positions are shown, with the mutant amino acids specified in order on the x-axis. The y-axis represents the log repression values, with a dashed line indicating the wild-type (WT) repression strength. These samples illustrate the effect of triple mutations on the TWIST1 protein within our library.

**Supplementary Figure 7:**
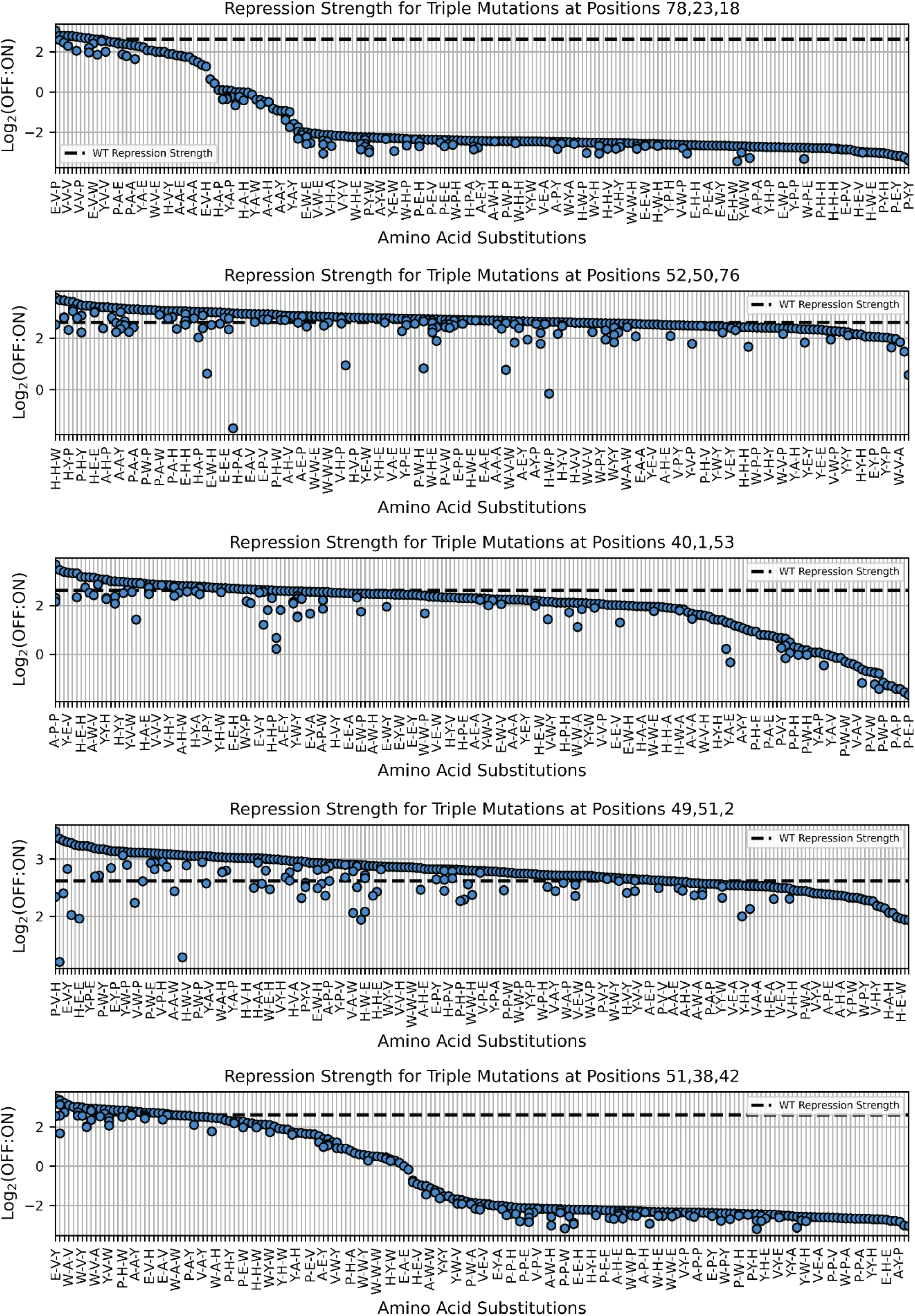
Repression strengths of selected ZNF705D [2-82] triple mutants Five representative samples of triple mutational data at specific positions are shown, with the mutant amino acids specified in order on the x-axis. The y-axis represents the log repression values, with a dashed line indicating the wild-type (WT) repression strength. These samples illustrate the effect of triple mutations on the ZNF705D protein within our library.

**Supplementary Figure 8:**
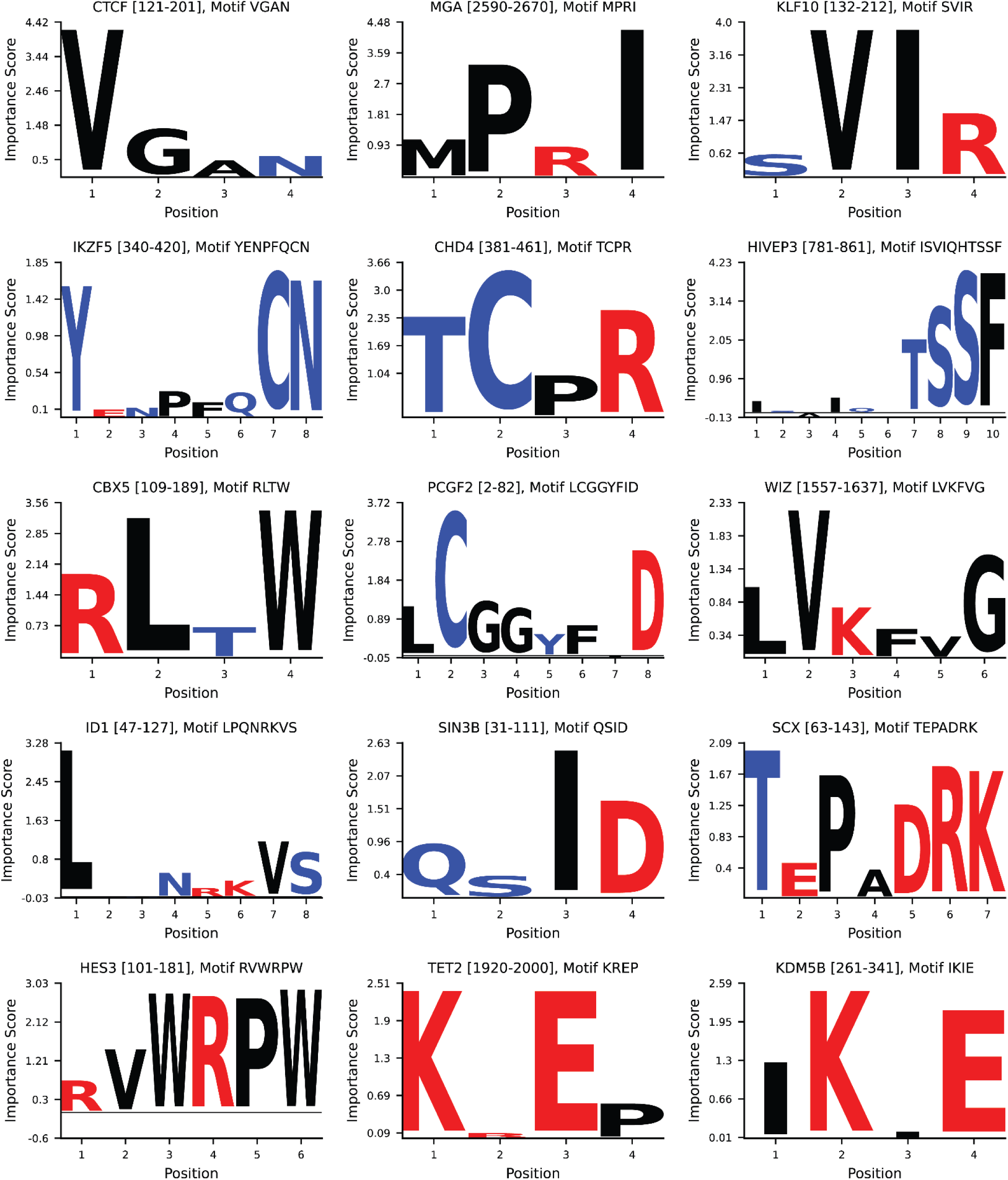
Top SLiMs with known nuclear ELM annotations identified in the RD-DMS screen Position weight matrices (PWMs) for the SLiMs with the highest importance score in indicated corresponding RDs. The importance score reflects the significance of individual AA in contributing to repression. High positive values indicate that mutations in these residues result in LoF, whereas low or negative values suggest that mutations in these residues do not significantly affect repression.

**Supplementary Figure 9:**
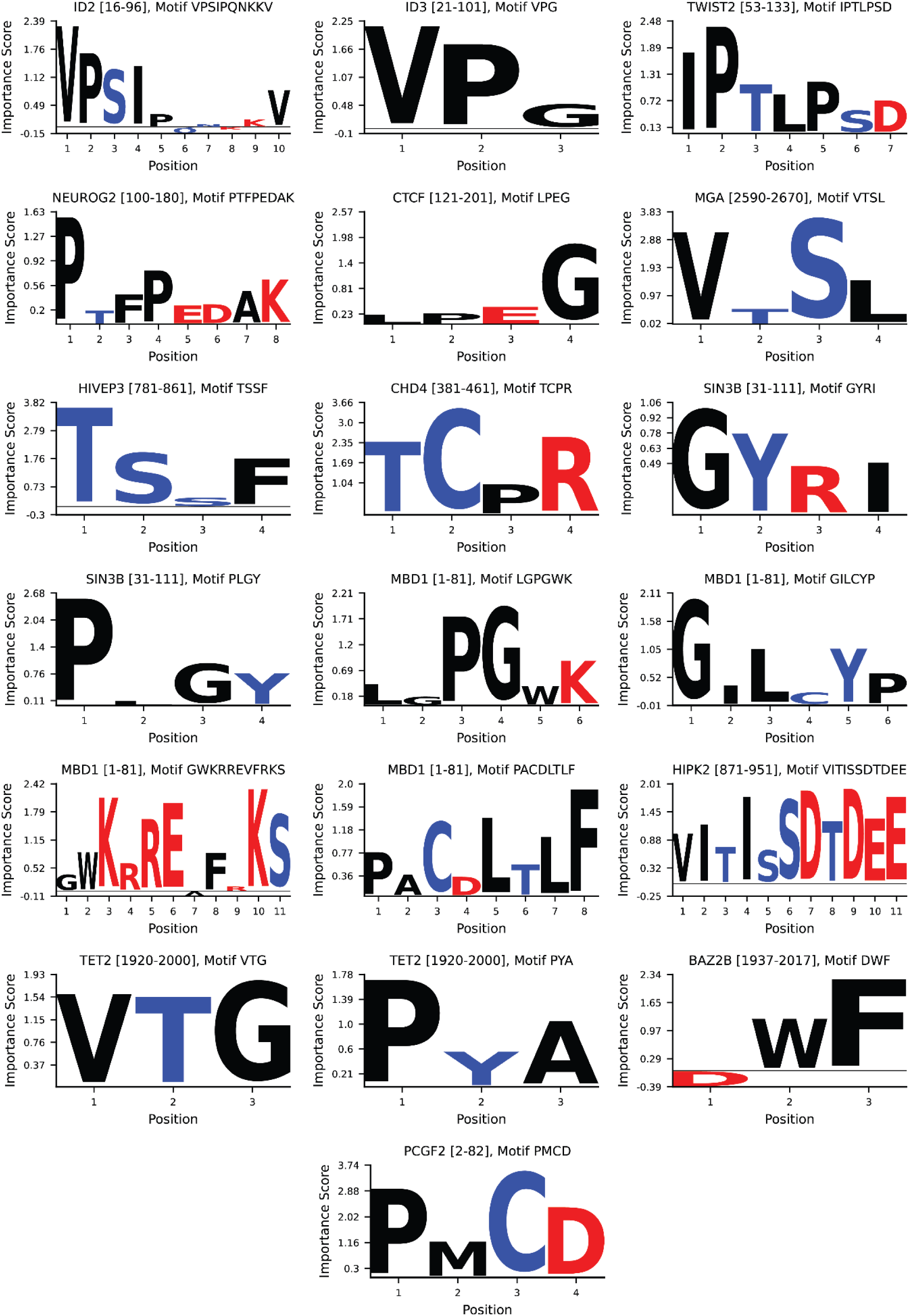
SLiMs without ELM annotations identified in the RD-DMS screen Position weight matrices (PWMs) for the SLiMs with no ELM annotation in indicated corresponding RDs. The importance score reflects the significance of individual AA in contributing to repression. High positive values indicate that mutations in these residues result in LoF, whereas low or negative values suggest that mutations in these residues do not significantly affect repression.

**Supplementary Figure 10:**
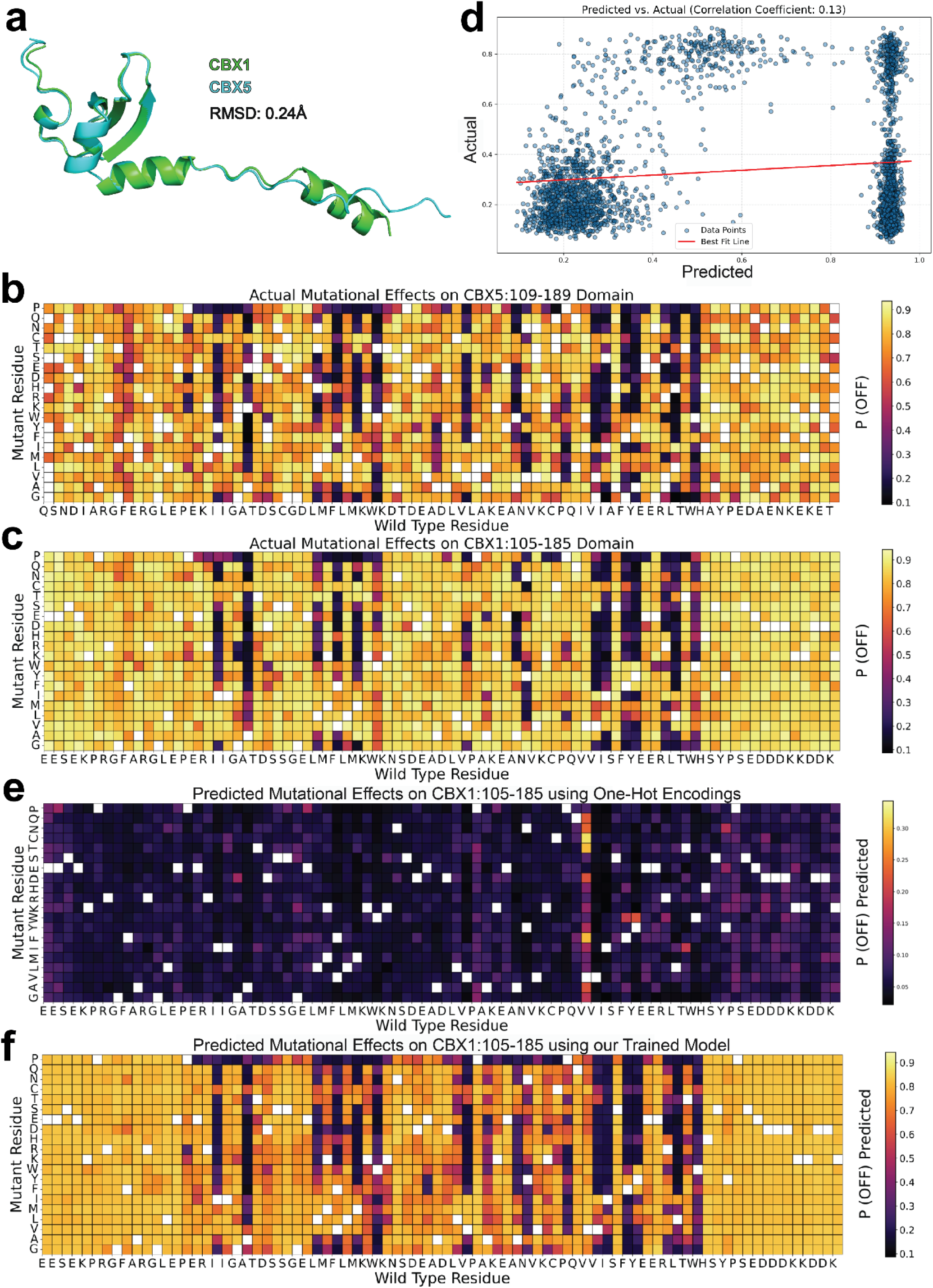
An architecture that considers sequence, structure, and biochemical features performs better than a neural network driven by one-hot encodings a. Superimposed structures of CBX1 (green) and CBX5 (cyan), exhibiting structural congruence with a root-mean-square deviation (RMSD) of 0.24Å. b. Heatmap of the measured mutational effects on the CBX5 [109-189] chromo domain, with color intensities indicating the P(OFF) values for each mutation. c. (**c**) for CBX1 [105-185]. d. Scatter plot showing predicted vs. actual P(OFF) values for CBX1 [105-185]: predictions were generated by a fully connected neural network leveraging one-hot encodings of AA sequences, with a training set that comprised of values for CBX5 [109-189], CBX3 [104-184], and CBX4 [481-561]. e. Heatmap of the predicted mutational effects on CBX1 [105-185] derived from one-hot encodings. f. Heatmap of the predicted mutational effects on CBX1 [105-185] from our trained model.

**Supplementary Figure 11:**
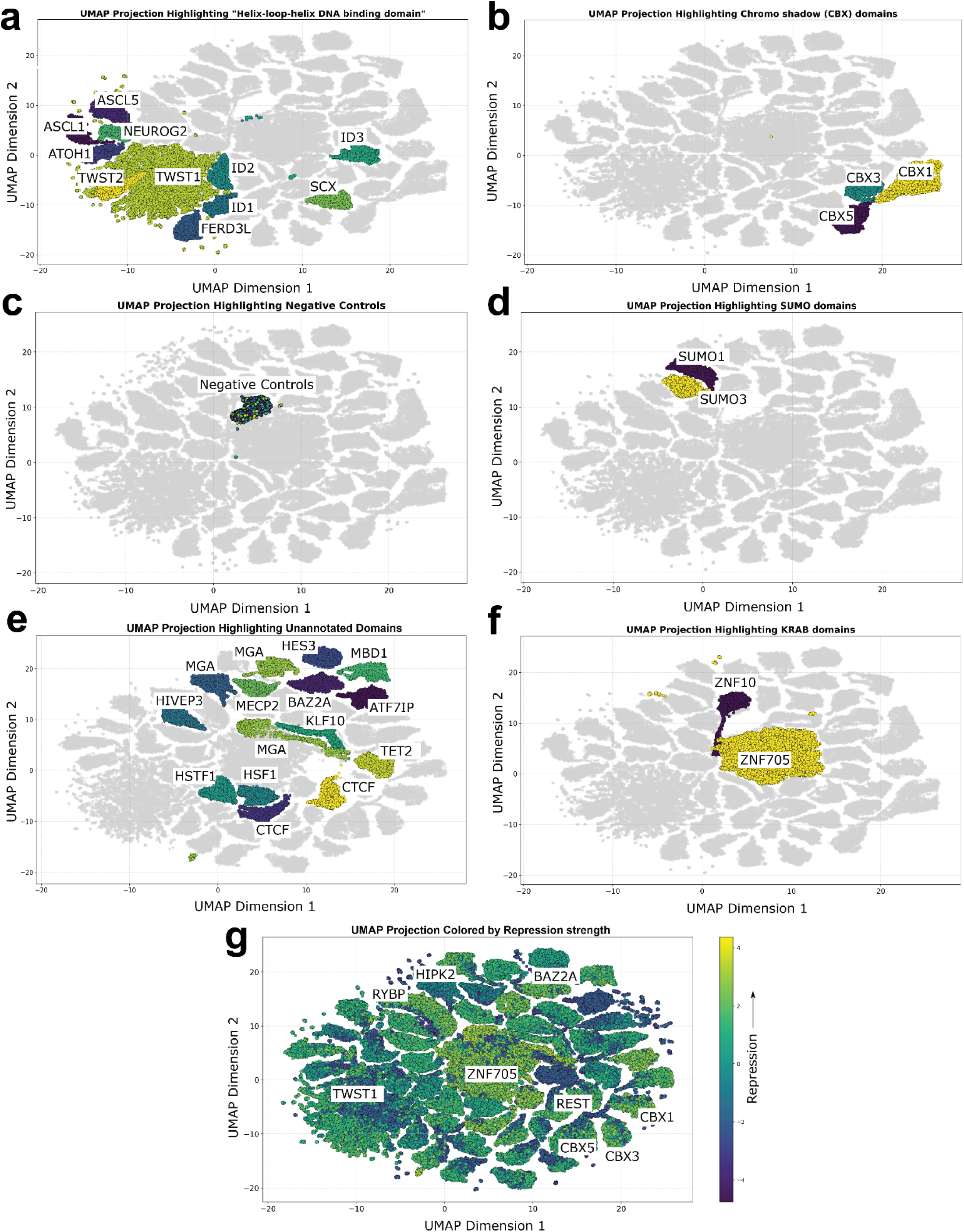
UMAP Projections of Protein Domains Using ESM-2 Embeddings. Panels A-F present 2D UMAP projections of various protein domains, with each panel accentuating a specific domain category through distinct color coding a. Helix-loop-helix DNA-binding domains are spotlighted, showcasing their clustering patterns. b. The Chromo shadow (CBX) domain family is highlighted, with an emphasis on its members. c. Negative control domains from our library are included for model validation. d. SUMO domains, especially SUMO1 and SUMO3, are illustrated. e. Unannotated domains from our collection are brought to the fore. f. KRAB domains are displayed, revealing their spatial distribution. g. The entire UMAP is color-mapped to signify repression strength, spanning from low (purple) to high (green), indicating poor clustering according to repression strength.

**Supplementary Figure 12:**
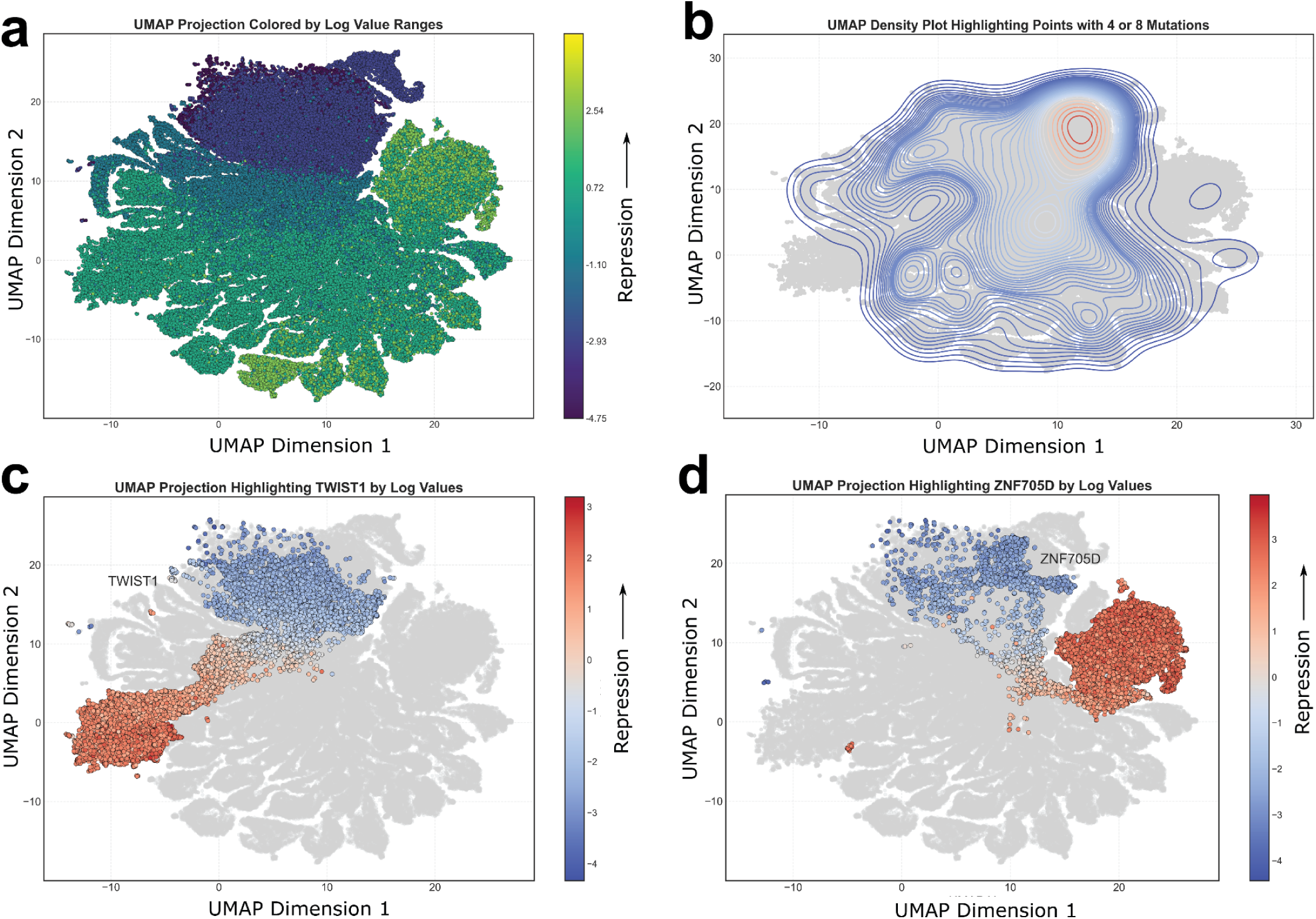
UMAP Projections of RDs using TENet embeddings. a. The UMAP projection colored by log value ranges, with the color gradient representing the degree of repression from low (purple) to high (green). b. A density plot on the UMAP projection highlights points with either 4 or 8 mutations, indicating regions of mutation concentration. c. The projection specifically highlights TWIST1 domains by log values, with the color scale indicating varying levels of repression. d. Similarly, ZNF705D domains are accentuated, with the color gradient reflecting repression intensity. This figure demonstrates the neural network’s capability to differentiate domain characteristics and mutation effects within the embedding space.

**Supplementary Figure 13.**
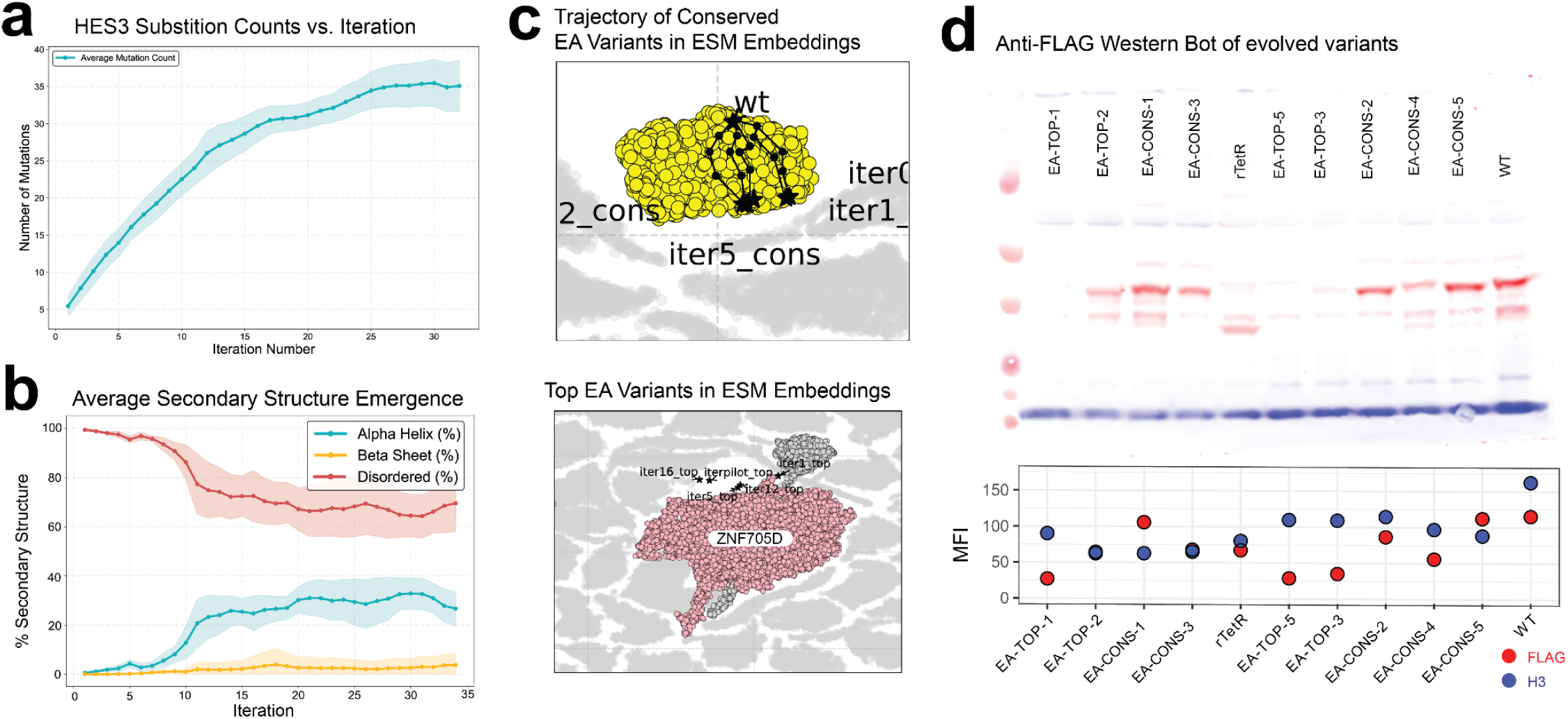
a. The number of total mutations accumulated with each evolutionary iteration. This value progressively increases as the EA adds more mutations. b. Average percentage over 5 trials of predicted alpha helix (turquoise), beta sheet (yellow), or disordered (red) AAs within the evolved populations at every iteration of the maximization task. c. Projected trajectory of evolved, relatively conserved (top) or top variants (bottom) of HES [101-181] in latent UMAP space derived from ESM embeddings. d. FLAG western (red) measuring expression of individual FLAG-tagged rTetR-HES3 variants in K562 reporter cells (top). H3 was measured as a housekeeping control (blue). Median fluorescence intensity (MFI) for FLAG and H3 signal of all evolved variants (bottom).

**Supplementary Figure 14.**
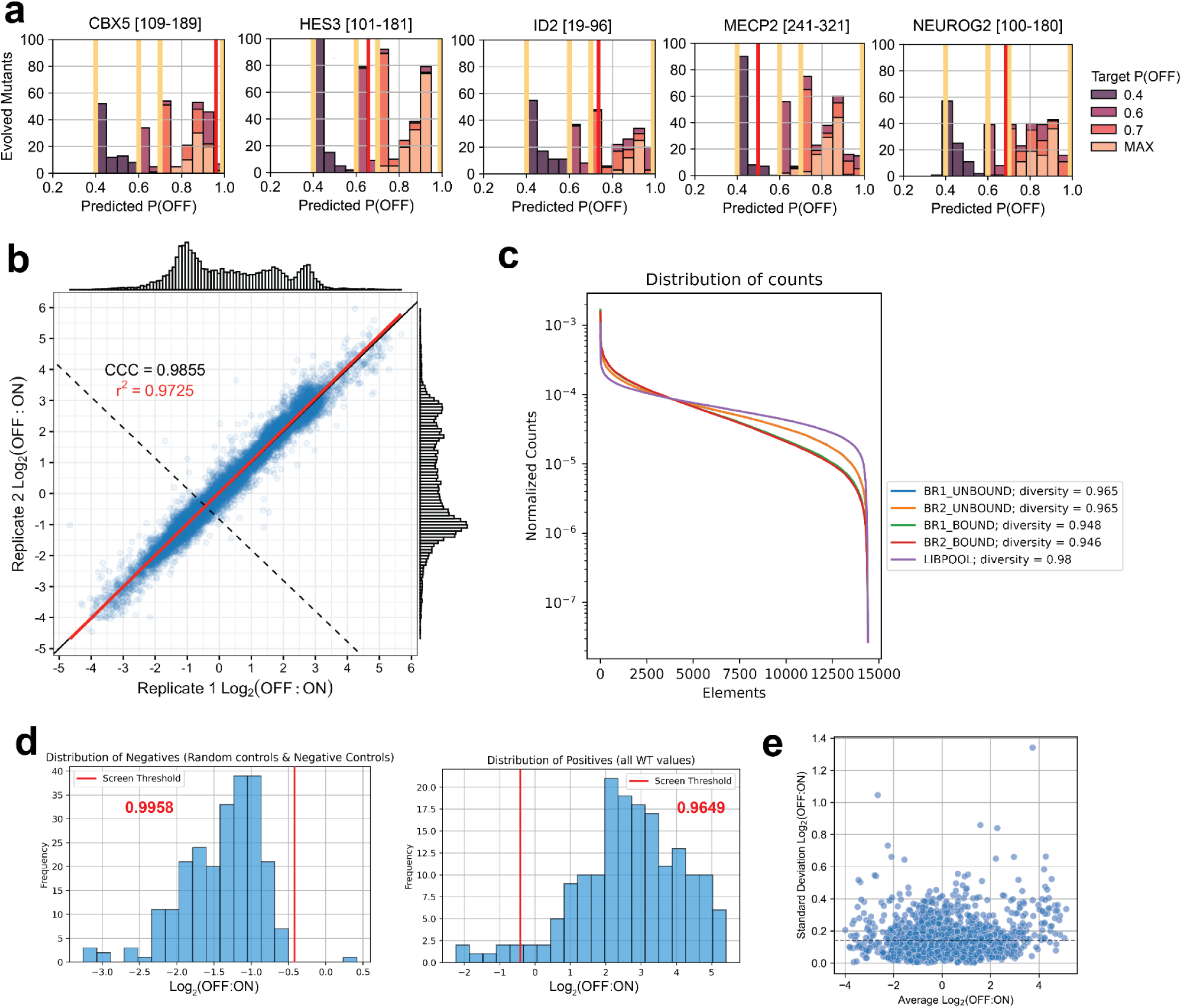
a. Distribution of predicted P(OFF) values for all RD-Design elements for each RD. A range of expected P(OFF) values were achieved by targeting 4 distinct target values (0.4, 0.6, 0.7, MAX). b. Reproducibility from two biological replicates of the joint CV-RD and EA-RD screens. The concordance correlation coefficient (CCC = 0.9855) is relative to the line with slope = 1 and y-intercept = 0 as shown in black. The coefficient of determination (r^2^ = 0.9725) is derived from the best fit linear regression as shown in red. The repressor threshold (dashed line) is set two SDs above the mean (**x̄ + 2σ**) of the negative controls. c. Recovery of RD-Design elements. Distribution of read counts for BOUND and UNBOUND samples from both BioReps compared to the LibPool. d. Confirmation of positive and negative element distributions in the RD-Design screen based on Log_2_(OFF:ON) ratio. The screen threshold (**x̄ + 2σ** of all negative controls) is indicated in red and the proportion of elements below (negative) or above (positive) the threshold for respective control types are indicated in red text. e. Average vs. Standard Deviation of Log_2_(OFF:ON) values for all AA sequences that were encoded by multiple degenerate codons in the RD-Design screen.

**Supplementary Figure 15.**
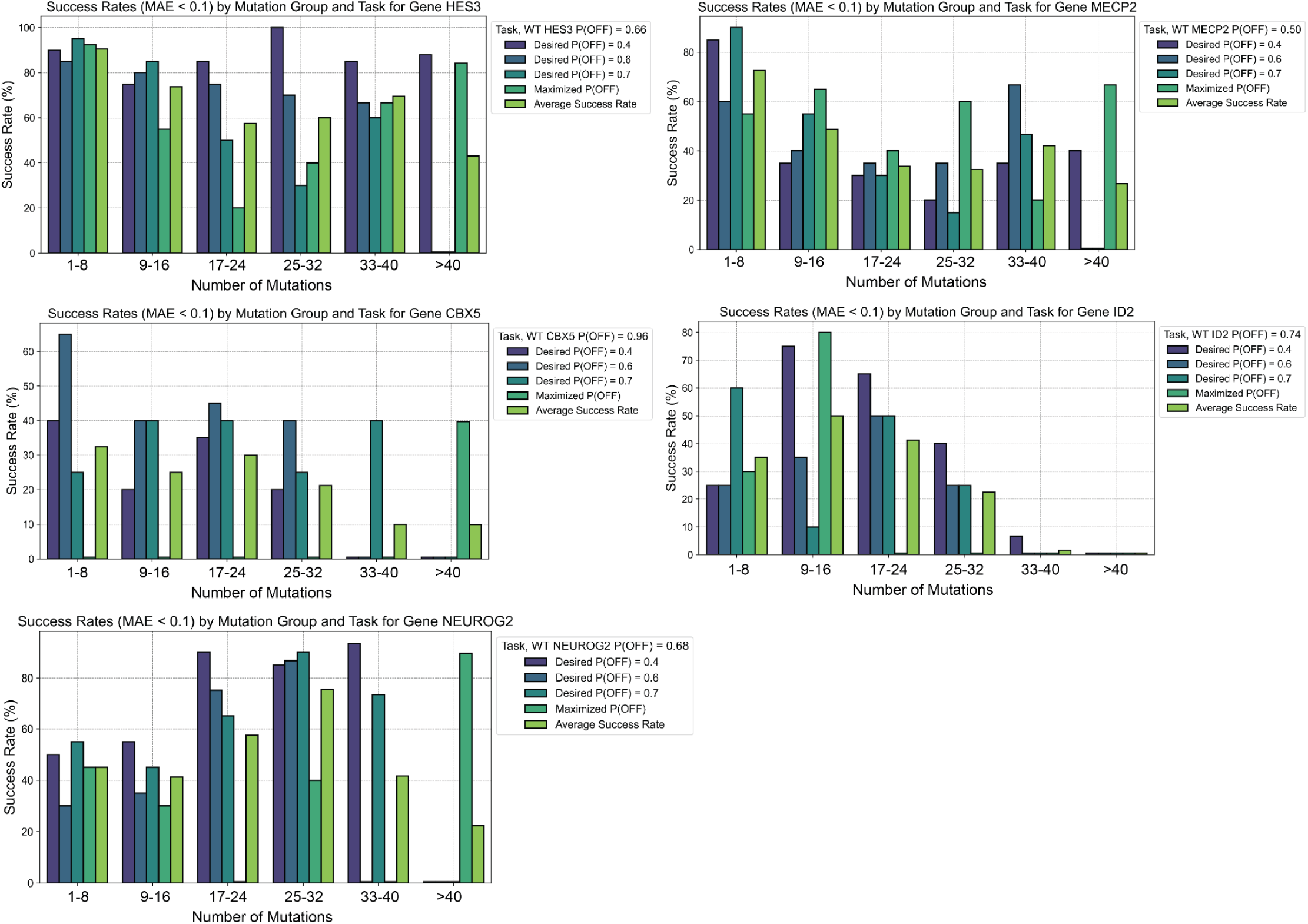
Success rates, defined by a mean average error (MAE) below 0.1, represent the comparison between the neural network-predicted performance and the measured values across various mutational intervals for each domain.

## Supplementary Tables

**Supplementary Table 1:** Repressor domains selected for deep mutational scanning and associated annotations.

**Supplementary Table 2:** Oligonucleotides libraries designed and ordered for this study.

**Supplementary Table 3:** Primer sequences used for library amplification, preparation, and sequencing. DNA fragment, vector, and primer sequences used for cloning constructs for individual domain validations.

**Supplementary Table 4:** HT-recruit processed analysis complete in this study.

**Supplementary Table 5:** Summary of all human missense variants that overlapped with measurements made by at least one sequence in our RD-DMS library. Functional LoF and significant variation are reported.

**Supplementary Table 6:** This table presents data on various functional motifs identified within disordered domains and their corresponding Eukaryotic Linear Motif (ELM) annotations. Each row corresponds to a motif instance with columns providing details on the motif sequence ("Slim"), the regular expression ("Regex"), the log value, and the importance score, which indicates the impact of each residue on the repression phenotype. The "DMS Label" specifies the domain and protein context, while "WT Seq" and "WT Value" describe the wild-type sequence and its associated repression value, respectively. The "Gene Names & Counts" column lists the different genes that the motif is found in, along with the number of occurrences, and the "ELM match?" column indicates whether an ELM match is present. Additional columns include "ELM Regex Matches" and "ELM Descriptions" for those instances with ELM matches, and "Resi Imp Scores," which detail the residue-specific importance scores. The "ELM Nuclear Regex Matches" and "ELM Nuclear Descriptions" columns only indicate the ELM annotations corresponding to the Nucleus cell compartment.

**Supplementary Table 7:** Comparative analysis of various models and their effectiveness in predicting protein repressor function. The table showcases modeling strategies including: (1) Cross-protein transfer amino acid descriptors (CPT AA Descriptors), (2) a six-layer architecture that utilizes one-hot encoding of the amino acid sequence (6 Layer One-Hot-Encoded FC Layers), (3) ESM embeddings averaged across positions (ESM Averaged), and (4) PADDLE, a 6-layer convolutional neural network. Our distinct method, which combines ESM embeddings, CPT amino descriptors, and ESMFold contact maps, is denoted as "TENet". The table presents both Pearson and Spearman Correlation coefficients across three mutation categories: Random Mutants, Single Mutants, and Windowed Mutants. All results pertain to the CBX1 domains, where no reads associated with this domain were used during training.

| MODEL | PEARSON CORRELATION |  |  | SPEARMAN CORRELATION |  |  |
| --- | --- | --- | --- | --- | --- | --- |
|  | RANDOM MUTANTS | SINGLE MUTANTS | WINDOWED MUTANTS | RANDOM MUTANTS | SINGLE MUTANTS | WINDOWED MUTANTS |
| CPT AA DESCRIPTORS | 0.12 | 0.84 | 0.79 | 0.16 | 0.50 | 0.77 |
| 6 LAYER ONE-HOT-ENCODED FC LAYERS | -0.09 | 0.13 | 0.36 | -0.00 | 0.12 | 0.38 |
| ESM AVERAGED | -0.09 | 0.36 | 0.62 | 0.10 | 0.43 | 0.70 |
| PADDLE | -0.13 | 0.06 | -0.02 | -0.18 | 0.08 | -0.05 |
| TENET | 0.53 | 0.88 | 0.90 | 0.49 | 0.56 | 0.75 |

**Supplementary Table 8:** Ablation study of the model developed for predicting repressive function. Four key features were explored to optimize the model: (1) ESM embeddings (labeled as ESM), (2) Self-attention maps from ESM Embeddings (denoted as Attention Maps), (3) Cross-protein transfer amino acid descriptors (indicated as CPT), and (4) Contact maps produced using ESMFold for structure prediction (labeled as ESMFold CM). Both graph convolutional networks (GCN) and graph attention networks (GAT) were evaluated as potential architectures for accurate repression function prediction. The table showcases the model’s performance on the CBX1 domain, for which no reads associated to this domain were shown during training.

|  | PEARSON CORRELATION |  |  |  | SPEARMAN CORRELATION |  |  |  |
| --- | --- | --- | --- | --- | --- | --- | --- | --- |
|  | FULL | RANDOM MUTANTS | SINGLE MUTANTS | WINDOWED MUTANTS | FULL | RANDOM MUTANTS | SINGLE MUTANTS | WINDOWED MUTANTS |
| ESM, CPT, CM & ATTENTION MAPS | 0.86 | 0.53 | 0.82 | 0.90 | 0.62 | 0.46 | 0.51 | 0.75 |
| **TENet (ESM, CPT & ESMFold CM)** | **0.91** | **0.53** | **0.88** | **0.90** | **0.65** | **0.49** | **0.56** | **0.75** |
| NO ATT. MAPS + NO ESMFOLD CM (ESM, CPT) | 0.83 | 0.34 | 0.78 | 0.88 | 0.62 | 0.33 | 0.52 | 0.77 |
| NO ATT. MAPS + NO ESM | 0.34 | 0.03 | -0.07 | 0.19 | 0.02 | 0.02 | 0.06 | 0.14 |
| GCN - COMPLETE GRAPH (ESM & CPT) | 0.81 | 0.10 | 0.74 | 0.84 | 0.60 | 0.03 | 0.50 | 0.70 |
| GCN - SPARSER (ESM & CPT) | 0.80 | 0.08 | 0.73 | 0.82 | 0.61 | 0.02 | 0.48 | 0.68 |
| GAT - COMPLETE GRAPH (ESM & CPT) | 0.79 | 0.07 | 0.73 | 0.82 | 0.60 | 0.02 | 0.48 | 0.67 |
| CPT AMINO ACID DESCRIPTORS | 0.80 | 0.12 | 0.84 | 0.79 | 0.58 | 0.16 | 0.50 | 0.77 |
| ESM + ESMFOLD CM (NO CPT) | 0.86 | 0.53 | 0.82 | 0.89 | 0.62 | 0.46 | 0.51 | 0.74 |

**Supplementary Table 9:** Pearson correlation coefficients (average ± standard deviation) for models trained on all data except all triple substitution data, and with or without double substitution data for ZNF707D or TWIST1, respectively. The models compare the effects of the presence or absence of double alanine mutations on the accuracy of predicting the triple mutant P(OFF) values for the indicated RD across 6 replicates.

| Effector Domain | Without Double Mutants | With Double Mutants |
| --- | --- | --- |
| ZNF705D [2-82] | 0.513 ± 0.081 | 0.609 ± 0.056 |
| TWIST1 [95-175] | 0.801 ± 0.002 | 0.813 ± 0.003 |

## Data Availability

The Illumina sequencing datasets generated in this study are available from the NCBI Sequence Read Archive (SRA) under BioProject ID PRJNA1147787 (https://www.ncbi.nlm.nih.gov/bioproject/1147787). All processed data and annotations are included as Supplementary Tables.

## Code Availability

The HT-recruit analysis software for processing the sequencing data for high-throughput recruitment assays is available on Github (https://github.com/bintulab/HT-recruit-Analyze). The kallisto software for processing the RNA-seq data is available on Github (https://github.com/pachterlab/kallisto). The deep learning model <u>T</u>ranscriptional <u>E</u>ffector <u>Net</u>work (TENet) for predicting repressor activity from amino acid sequence is available on GitHub (https://github.com/kundajelab/TENet): the repository contains the source code, detailed documentation, and instructions for running and analyzing results; custom scripts for data processing and analysis are provided within the same repository, including code for the evolutionary algorithm. Any additional custom code used for computational data processing and analysis are available from the authors upon request.

