## Extended Data for "Prediction and design of transcriptional repressor domains with large-scale mutational scans and deep learning"

**Extended Data:** Amino acid resolution data of the DMS for all protein tiles, including mapping to predicted structures and sequence conservation

a. Predicted percent disorder (blue) and AlphaFold 2 pLDDT values (orange) over the 80AA domain (**Methods: Domain classification and disorder prediction**). Residues with values over the disorder threshold (0.5) are predicted to be disordered within the context of the domain, with higher values indicating a higher probability of disorder.

b. Heatmaps filled by the Normalized $Log_2$(OFF:ON) values for each single substitution along the 80 AA domain. For normalization, the average $Log_2$(OFF:ON) is normalized to the mean of the WT values for each domain, such that WT is 0. The wildtype sequence is shown along the x-axis and corresponding substitutions are shown on the y-axis. Fill values correspond to the relative strengths where decreased strength (purple) are negative values and increased strength (yellow) are positive values. White boxes represent missing or wildtype data, while gray boxes represent values where data was missing for one BioRep.

c. Effect of all single substitutions on repressor strength where all single substitution data for a given residue is plotted in a box and whisker plot. Log2(OFF:ON) values have been normalized such that a value of 0 represents WT strength: positive values indicate increased repressor activity and negative values decreased activity.

d. Normalized $Log_2$(OFF:ON) values for all windowed mutants on repressor function. Locations of shuffled or reversed regions are indicated by the x-axis length of the bars (blue). The mean WT value is indicated in red with +/- 1 standard deviation shaded.

e. Relationship between observed conservation and the effect of single substitution on repression. ConSurf Conservation Scores report on the conservation of residues (red), where lower scores are given to more highly conserved AAs. Average $Log_2$(OFF:ON) values for all single substitutions of a given residue are superimposed (blue), where lower values indicate lower repressor activity. Pearson's R-squared and p-value (Wald Test) calculated from a linear least-squares regression between ConSurf Scores and $Log_2$(OFF:ON) values are indicated in the top-right corner.

f. The AlphaFold predicted structure for the 80 AA domain. Individual AAs are colored by mean $Log_2$(OFF:ON) values for all single substitutions at a given residue.

g. All screen data from both BioReps for a given domain. The WT values are shown in blue and the random and negative controls are shown in red and orange, respectively. The spread of all other mutant data is shown in gray.

h. (**b**) Heatmaps displaying repressor strength as calculated P(OFF) values are shown on the left. White boxes indicate missing data in one or both BioReps or WT data. On the right, the predicted P(OFF) values for each single substitution by the best NN model, which is trained using WT and Windowed-shuffle data, are shown.

### ASCL1 [106-186] Helix-loop-helix DNA binding domain

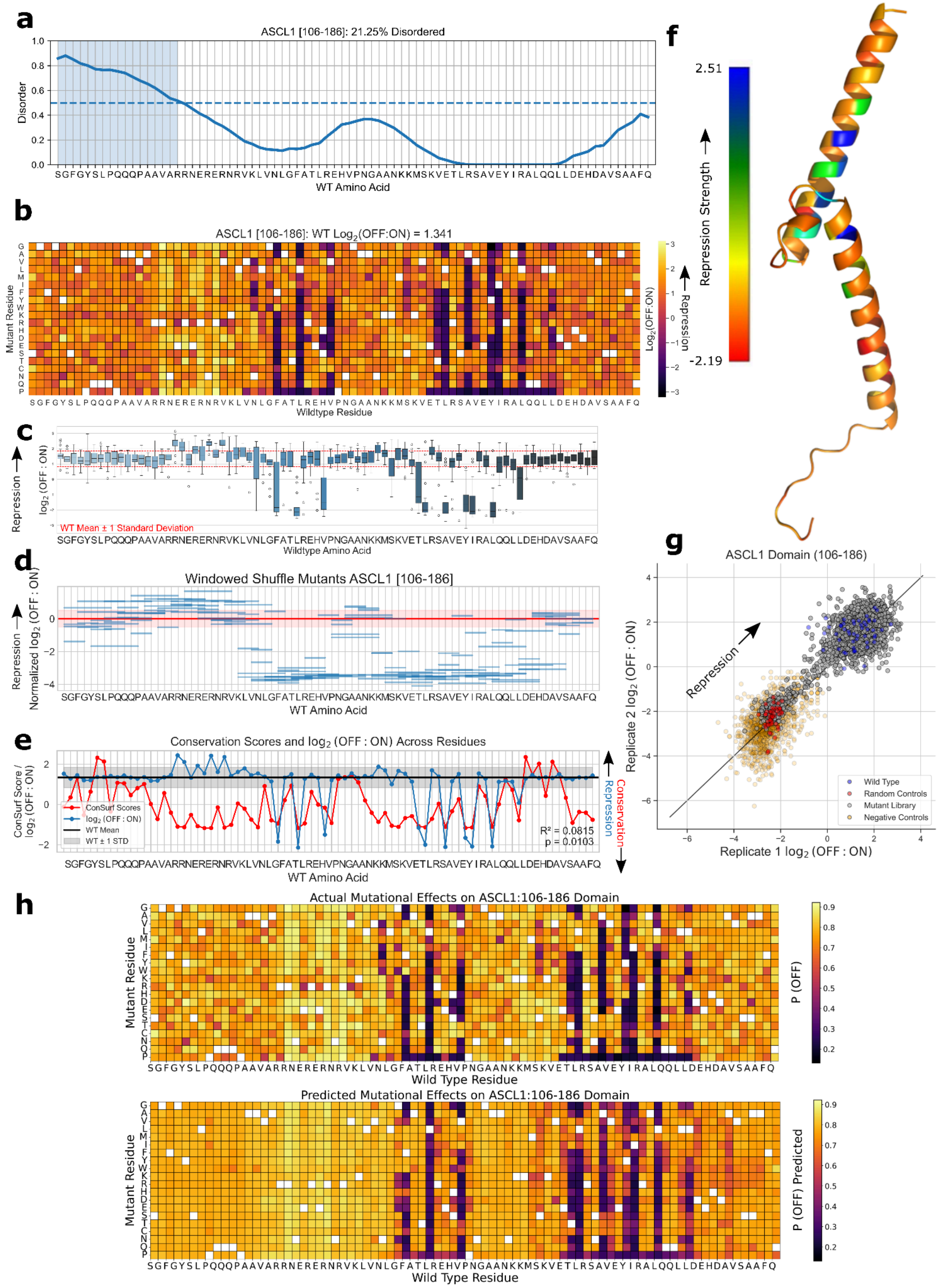

### ASCL5 [143-223] Helix-loop-helix DNA binding domain

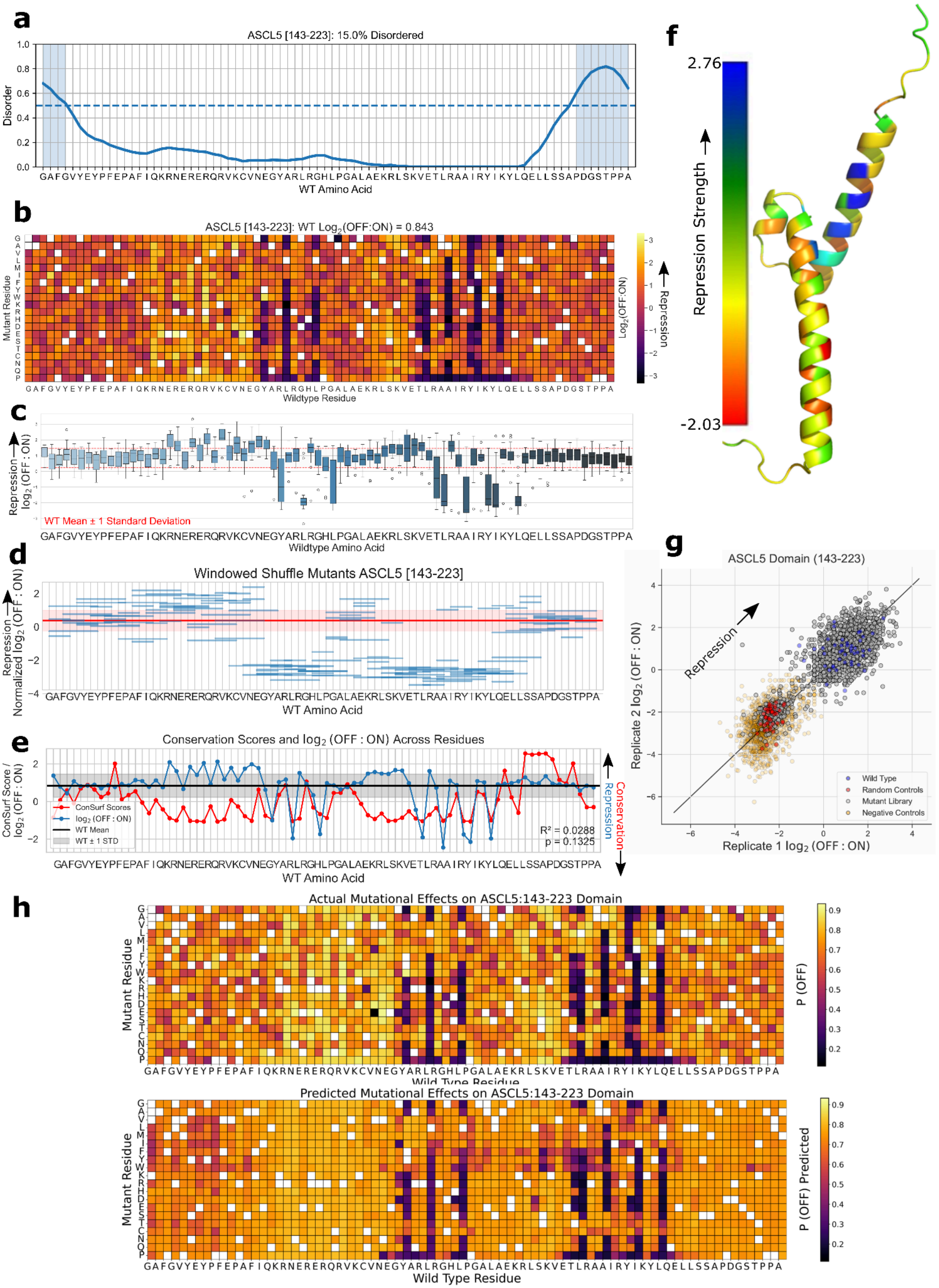

### ATF7IP [962-1042] Unannotated domain

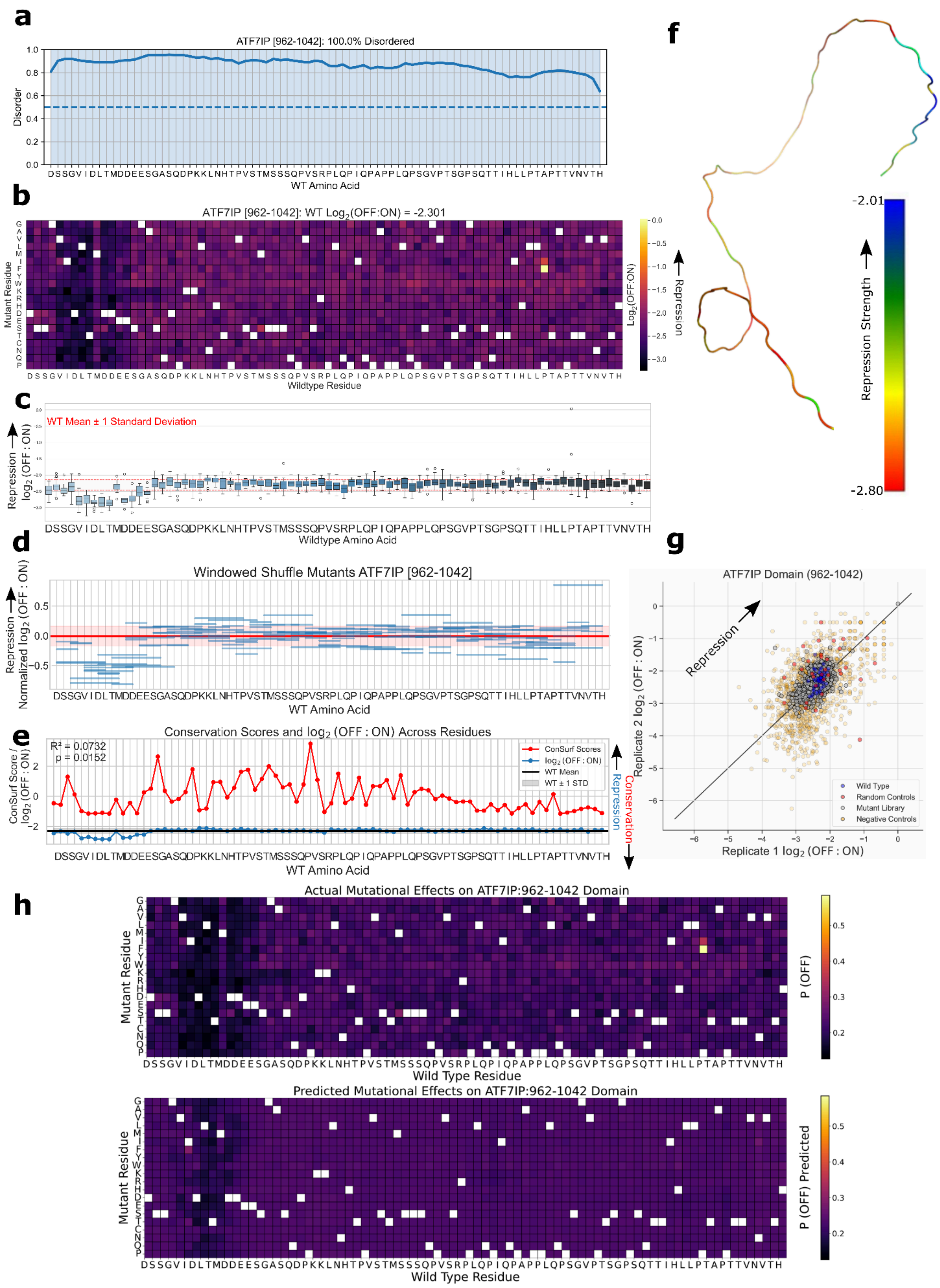

### ATOH1 [147-227] Helix-loop-helix DNA binding domain

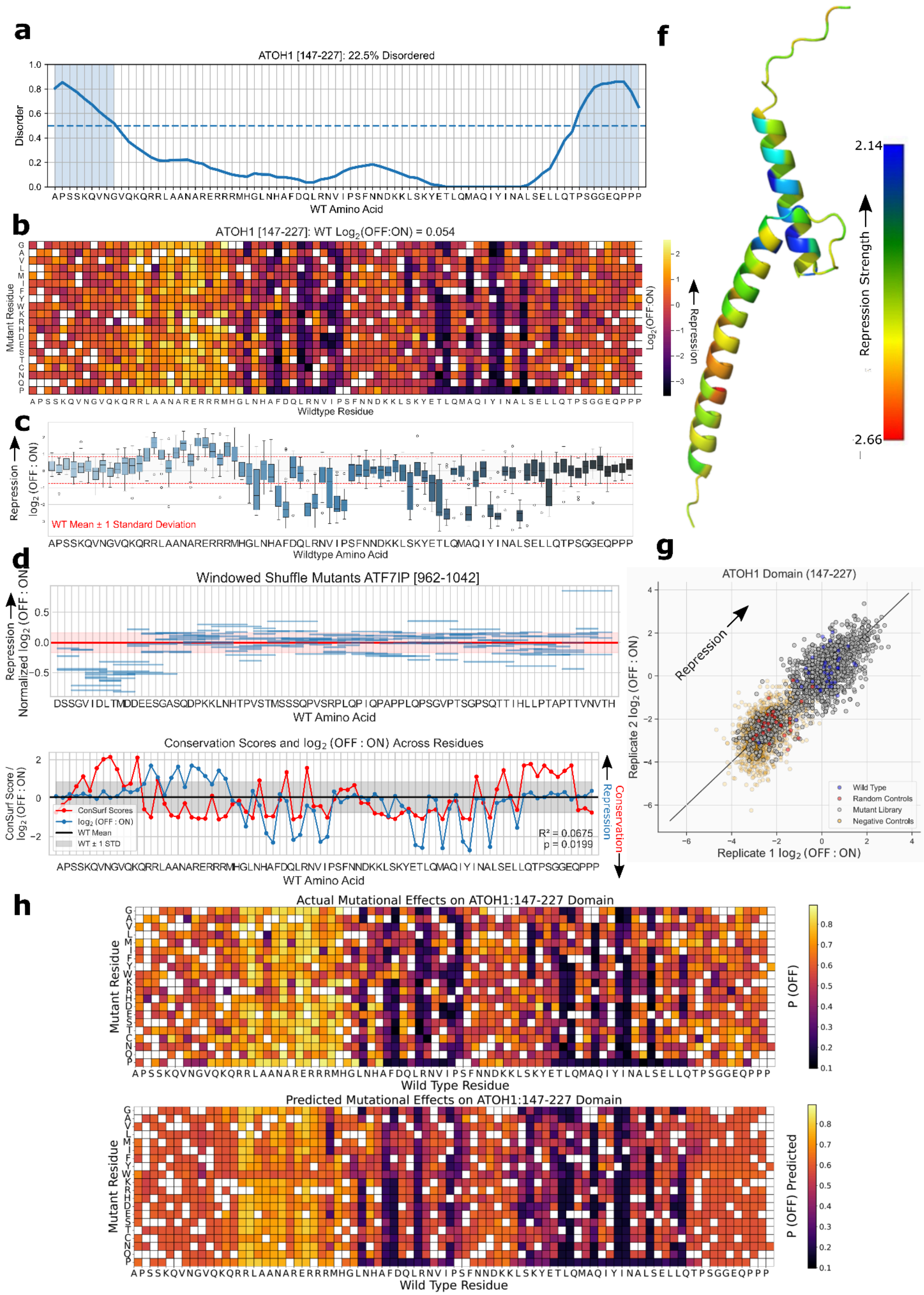

### BAZ2A [1211-1291] Unannotated domain

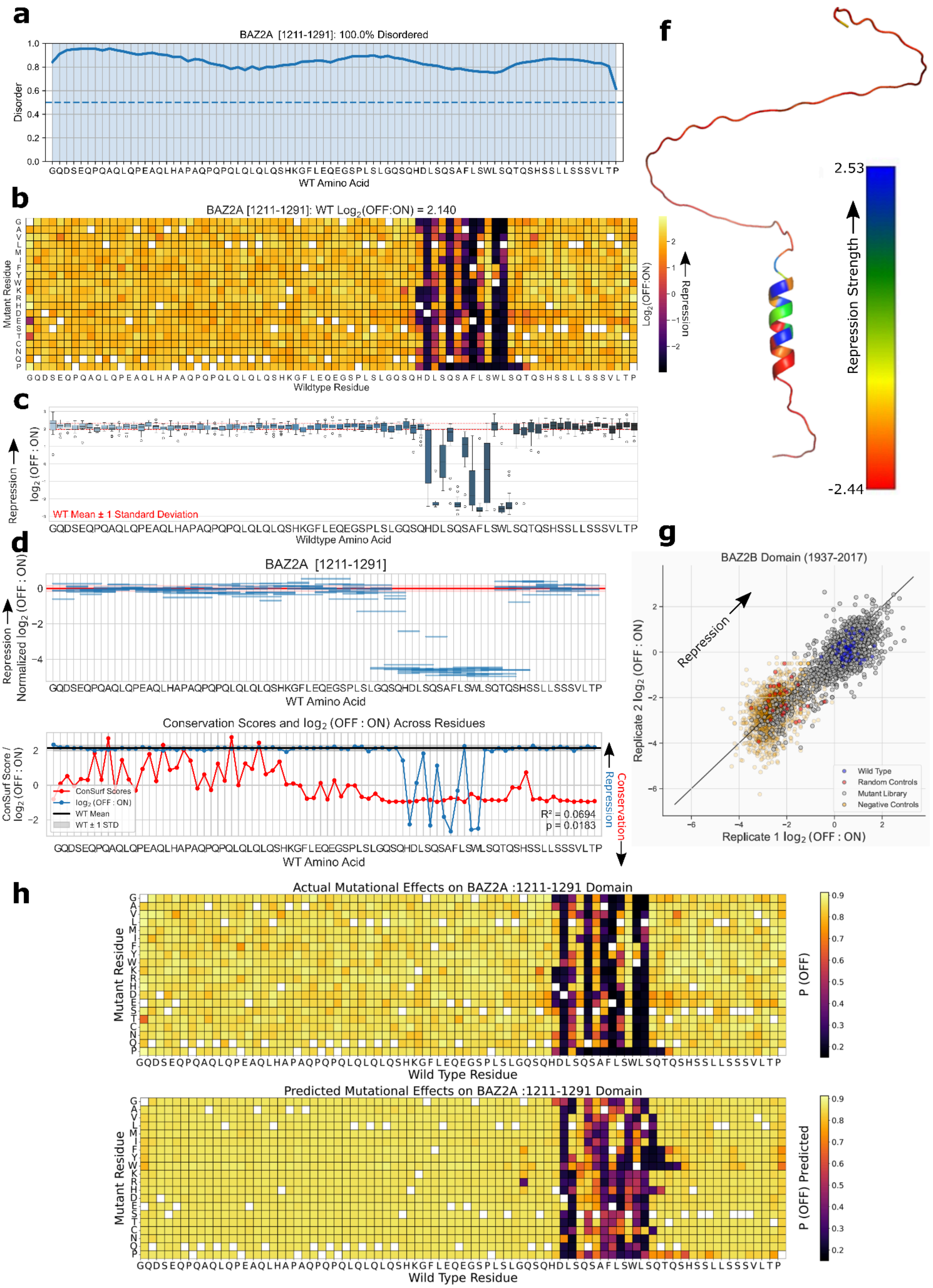

### BAZ2B [1937-2017] PHD-finger domain

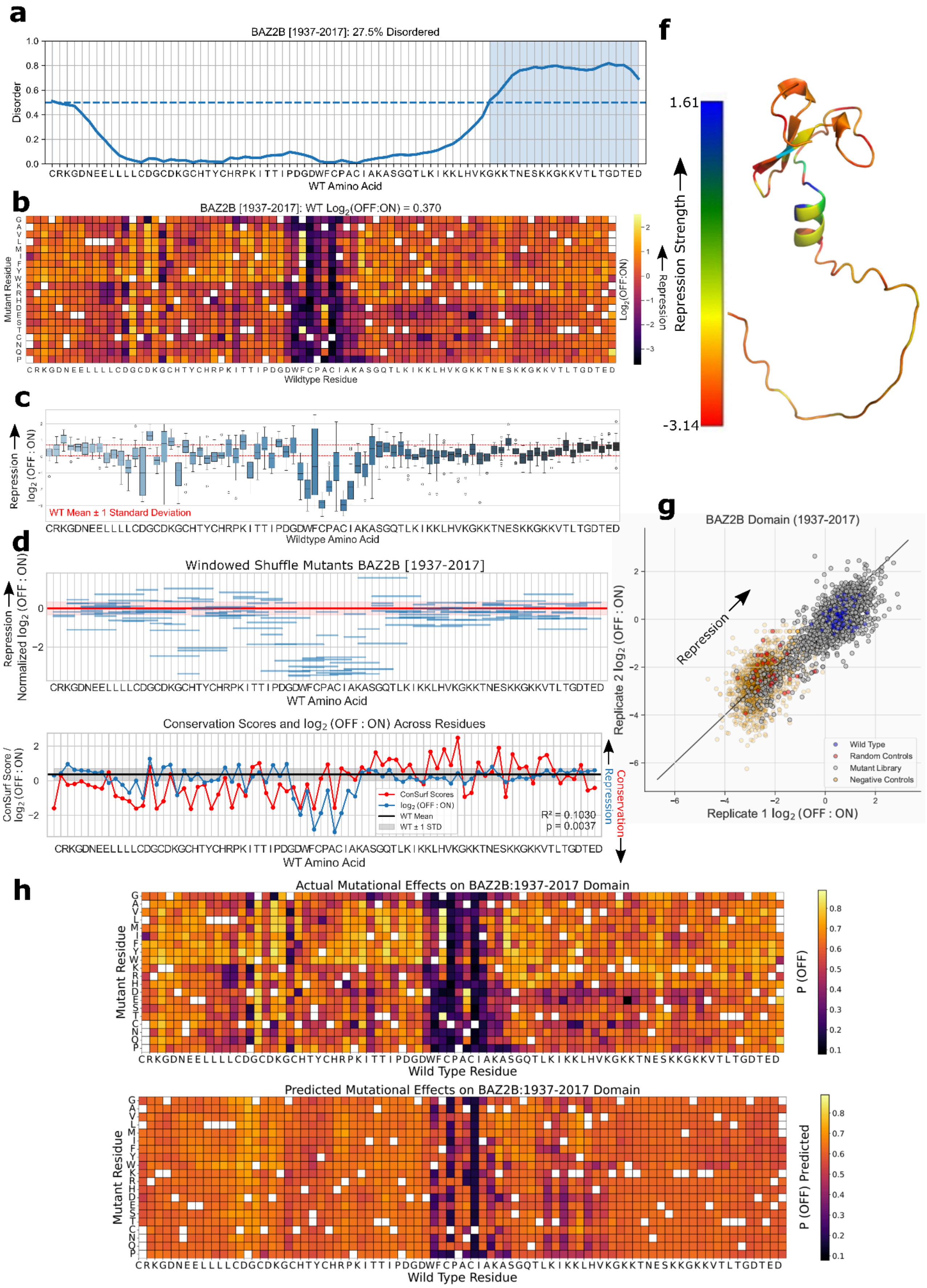

### CBX1 [105-185] Chromo shadow domain

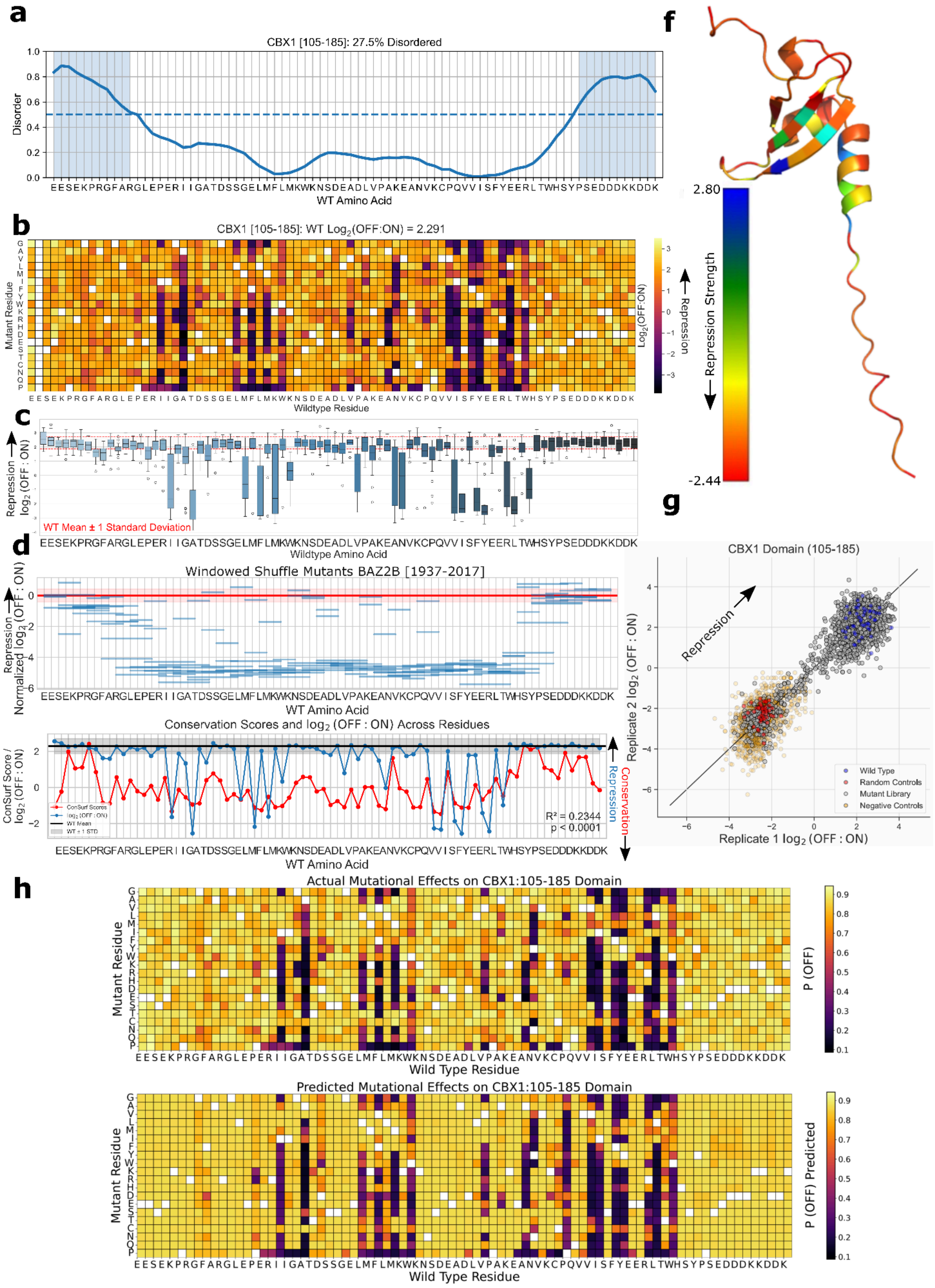

### CBX3 [104-184] Chromo shadow domain

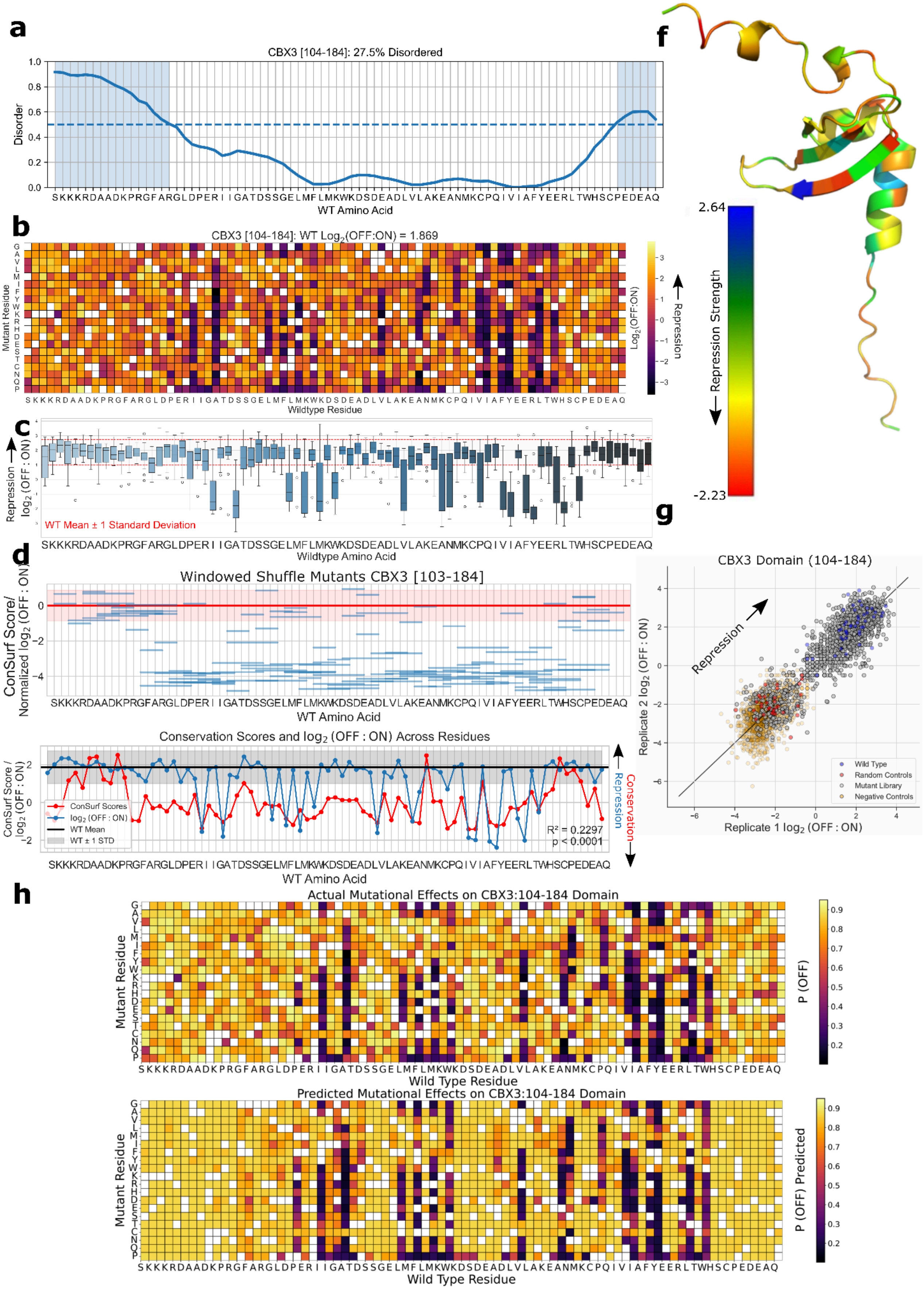

### CBX4 [481-561] CBX family C-terminal motif

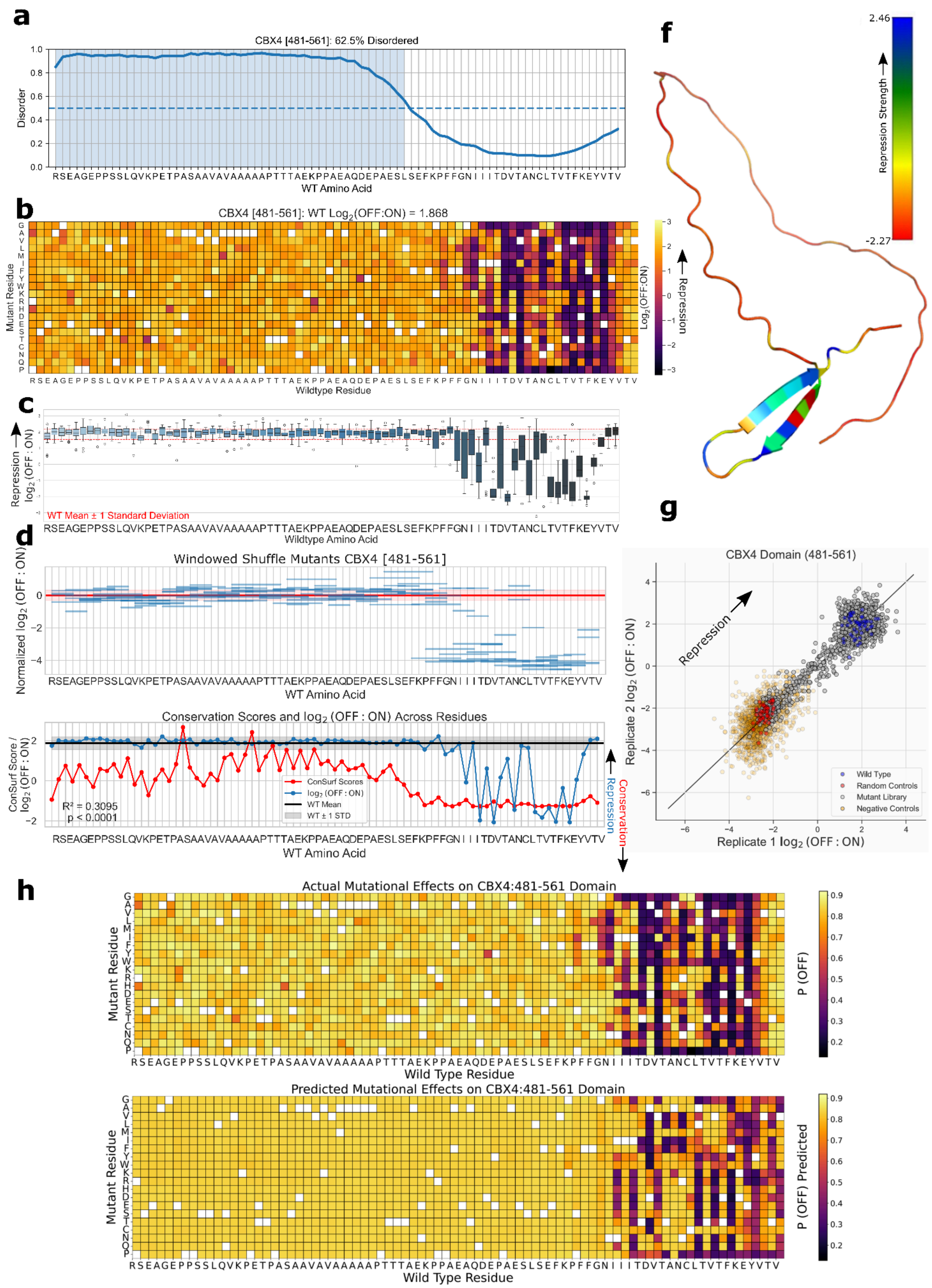

### CBX5 [109-189] Chromo shadow domain

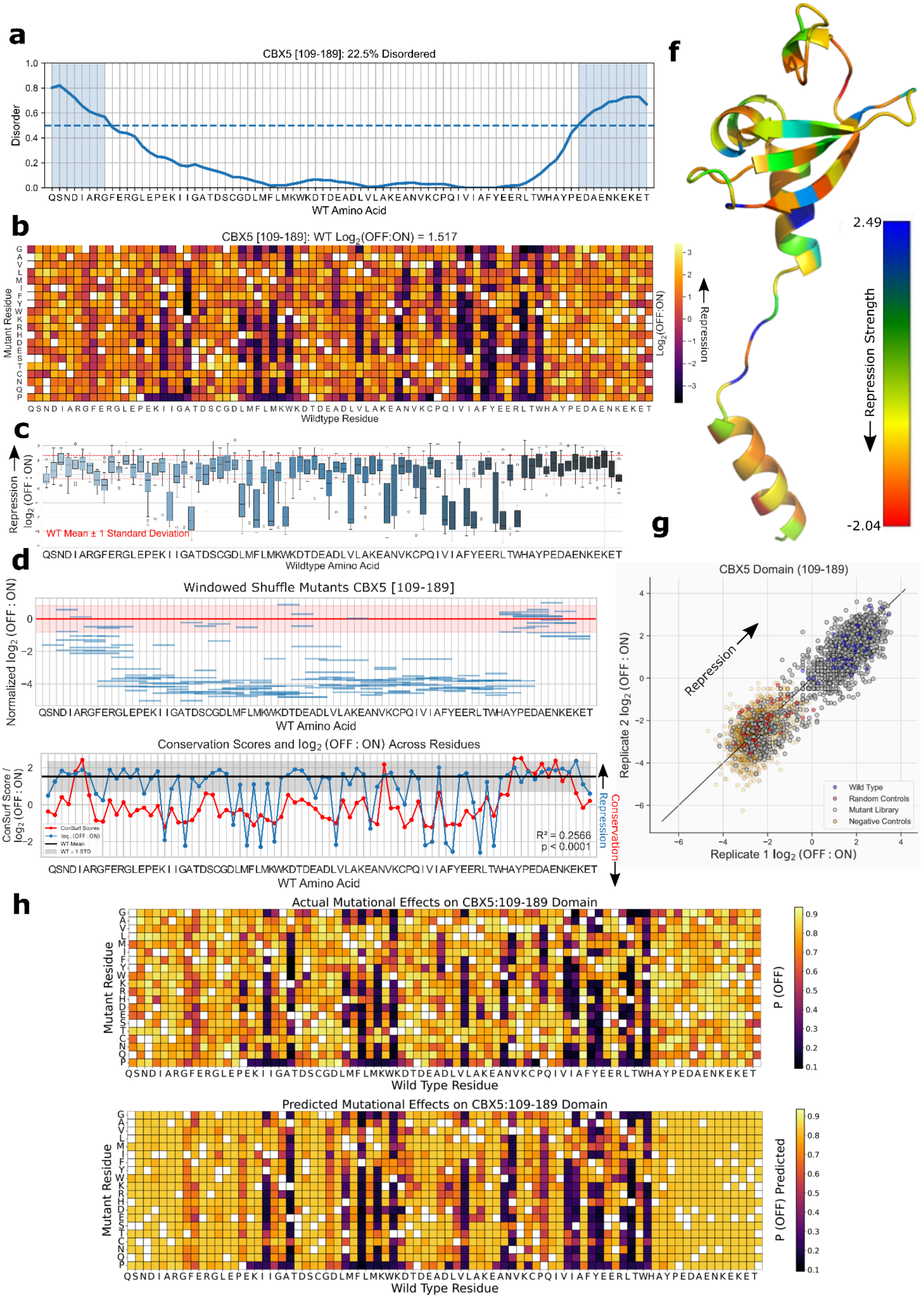

### CBX5 [111-191] Chromo shadow domain

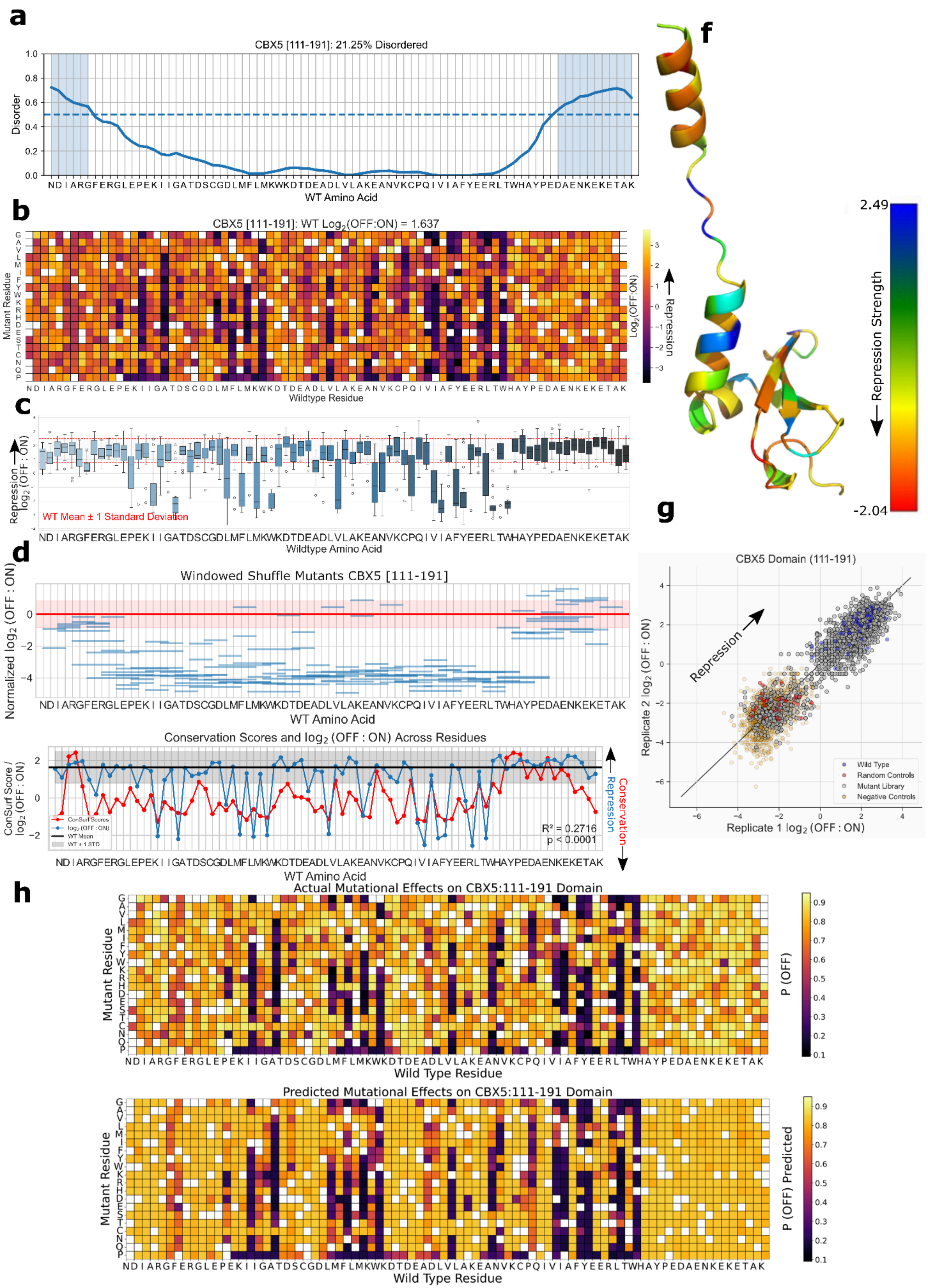

### CDY2A [2-82] Chromo (CHRromatin Organisation MOdifier) domain

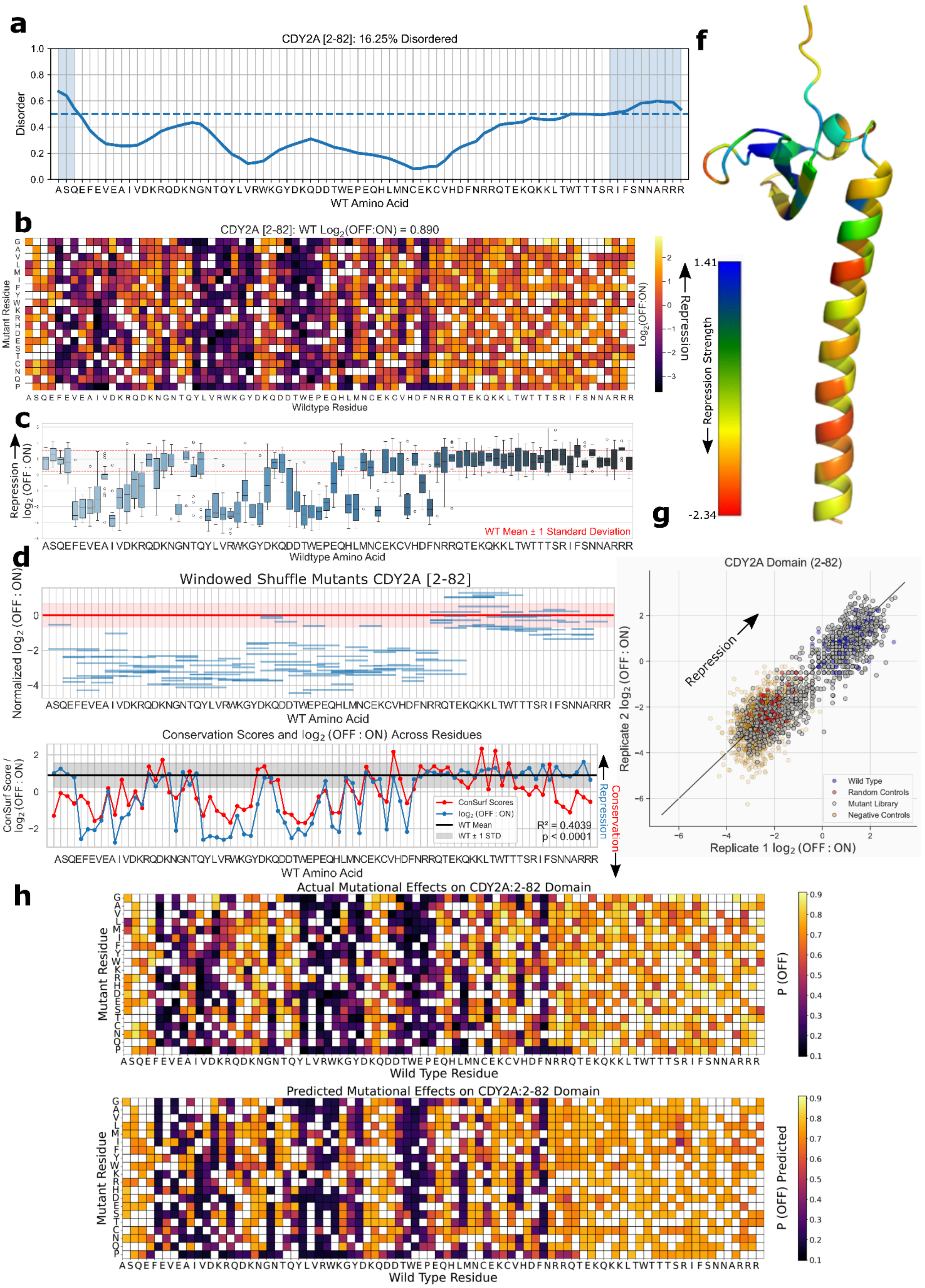

### CHD4 [381-461] PHD-finger domain

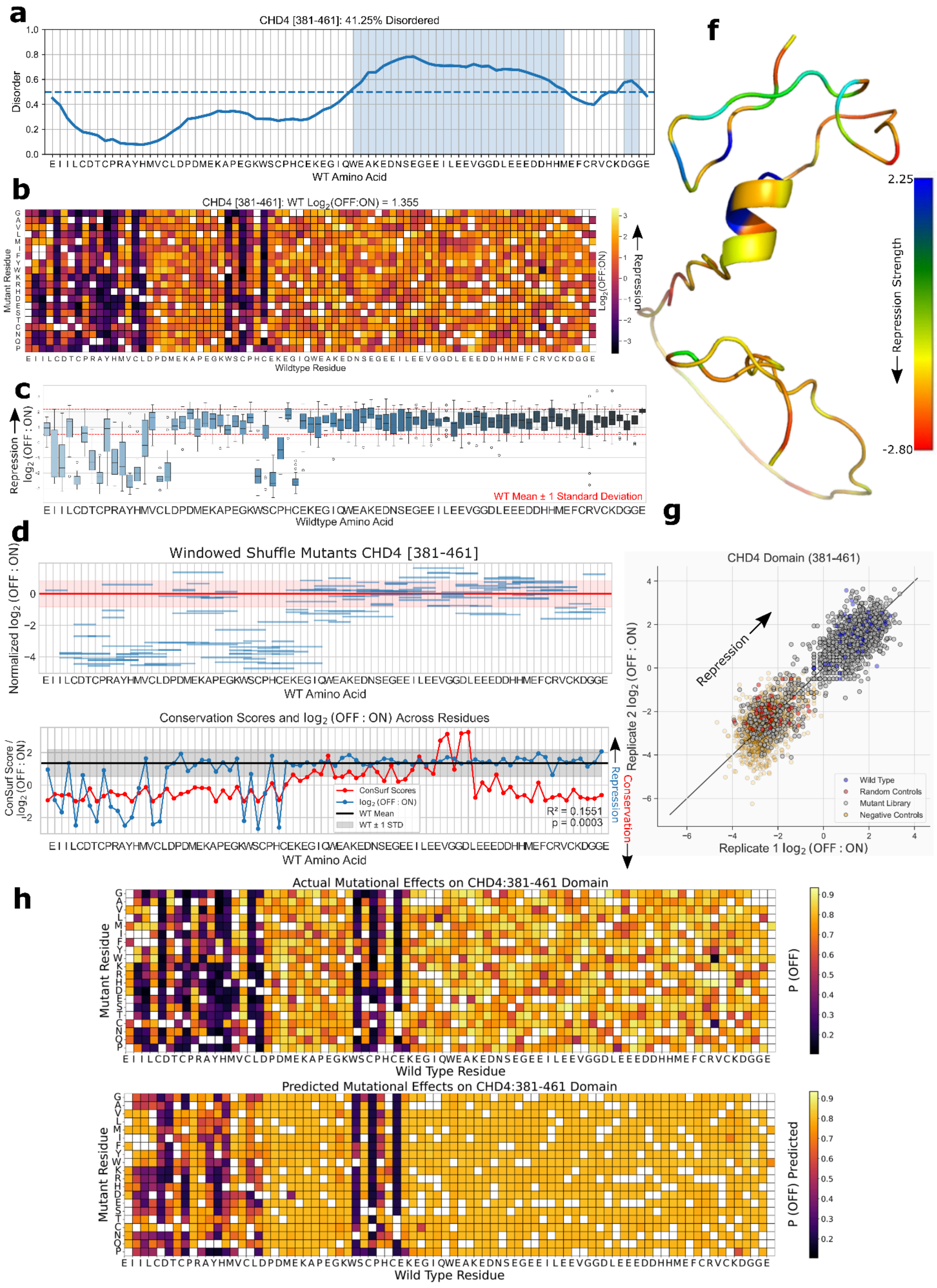

### CTCF [121-201] Unannotated domain

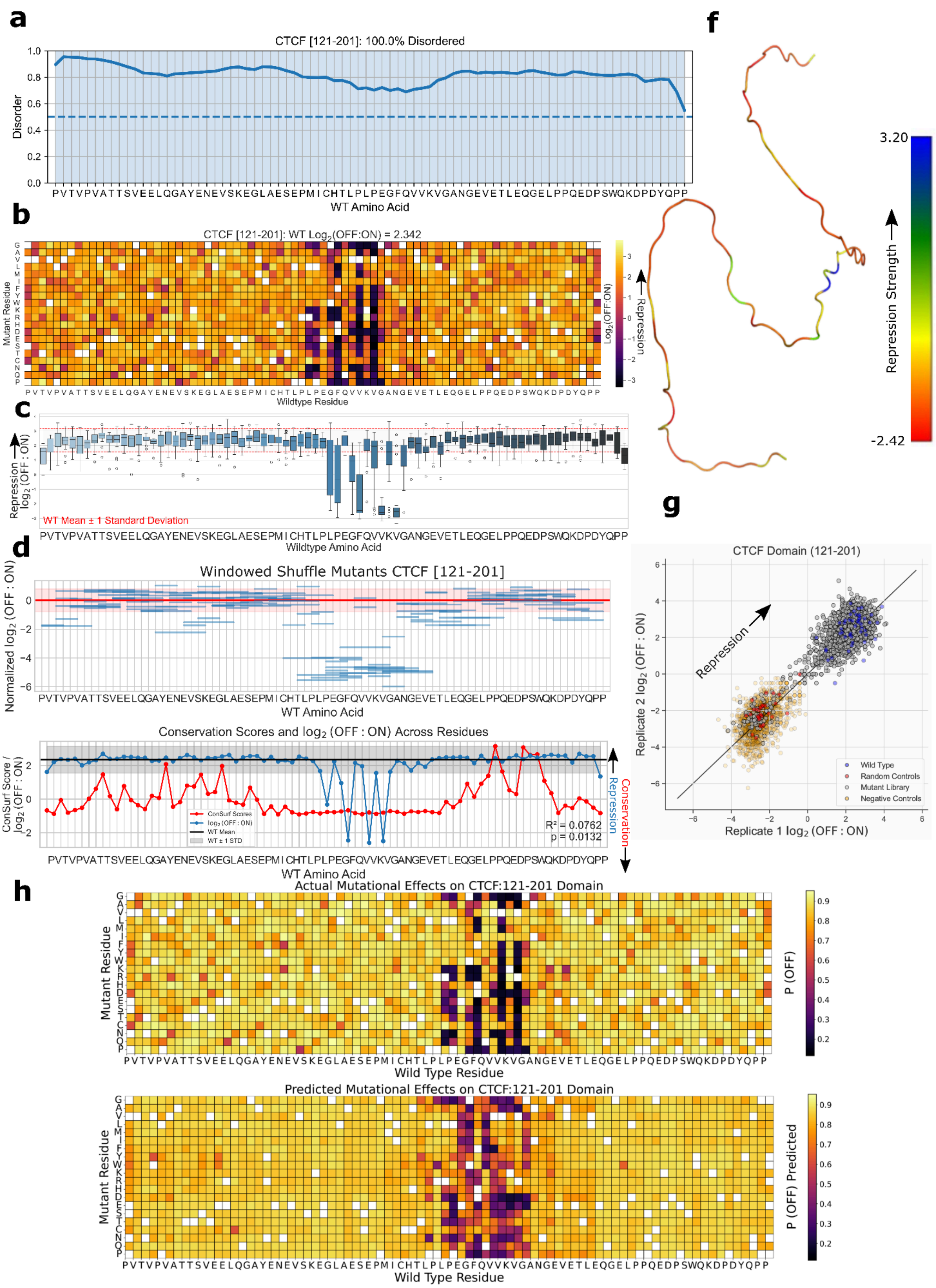

### FERD3L [87-167] Helix-loop-helix DNA binding domain

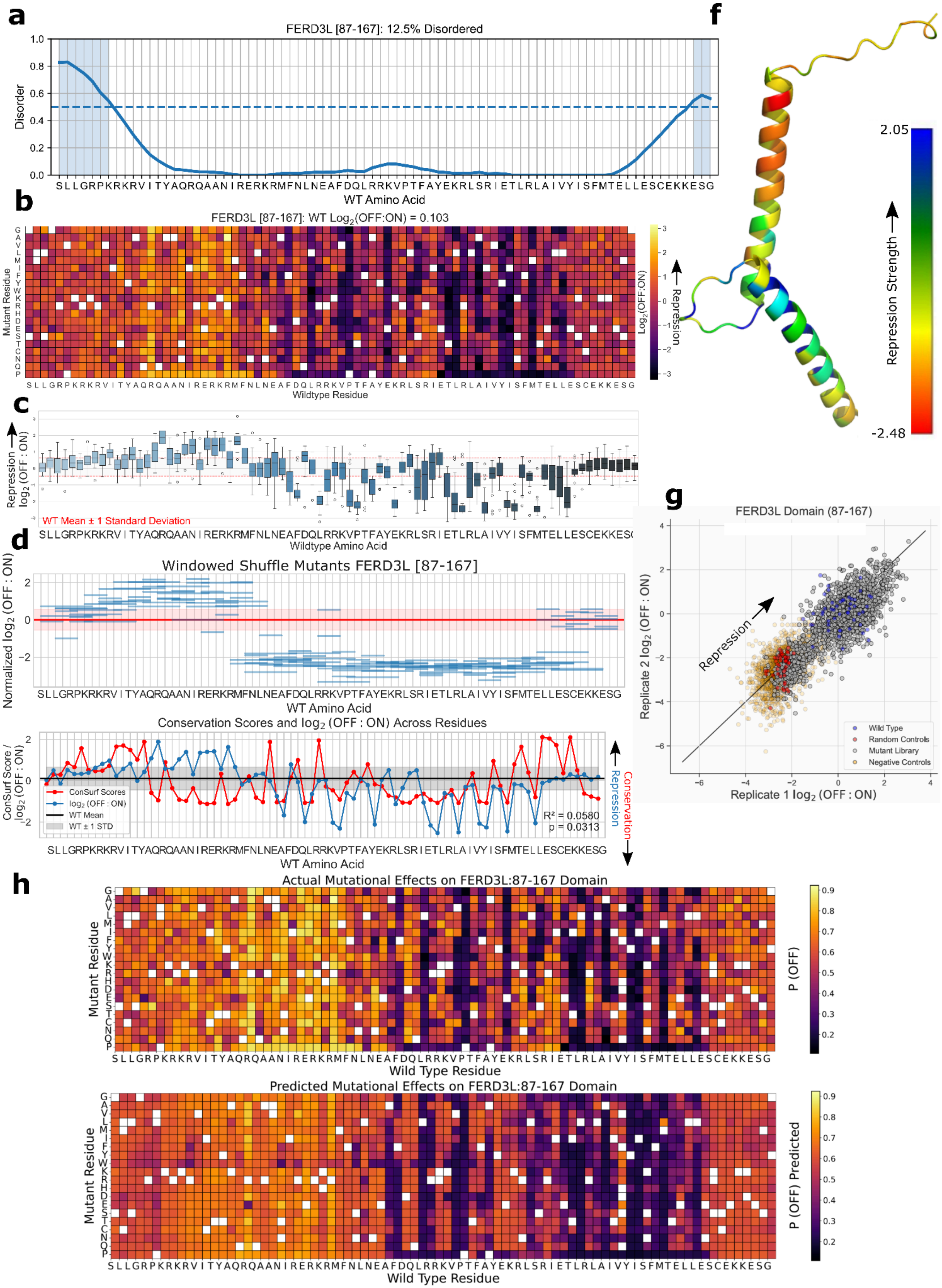

### HES3 [101-181] Unannotated domain

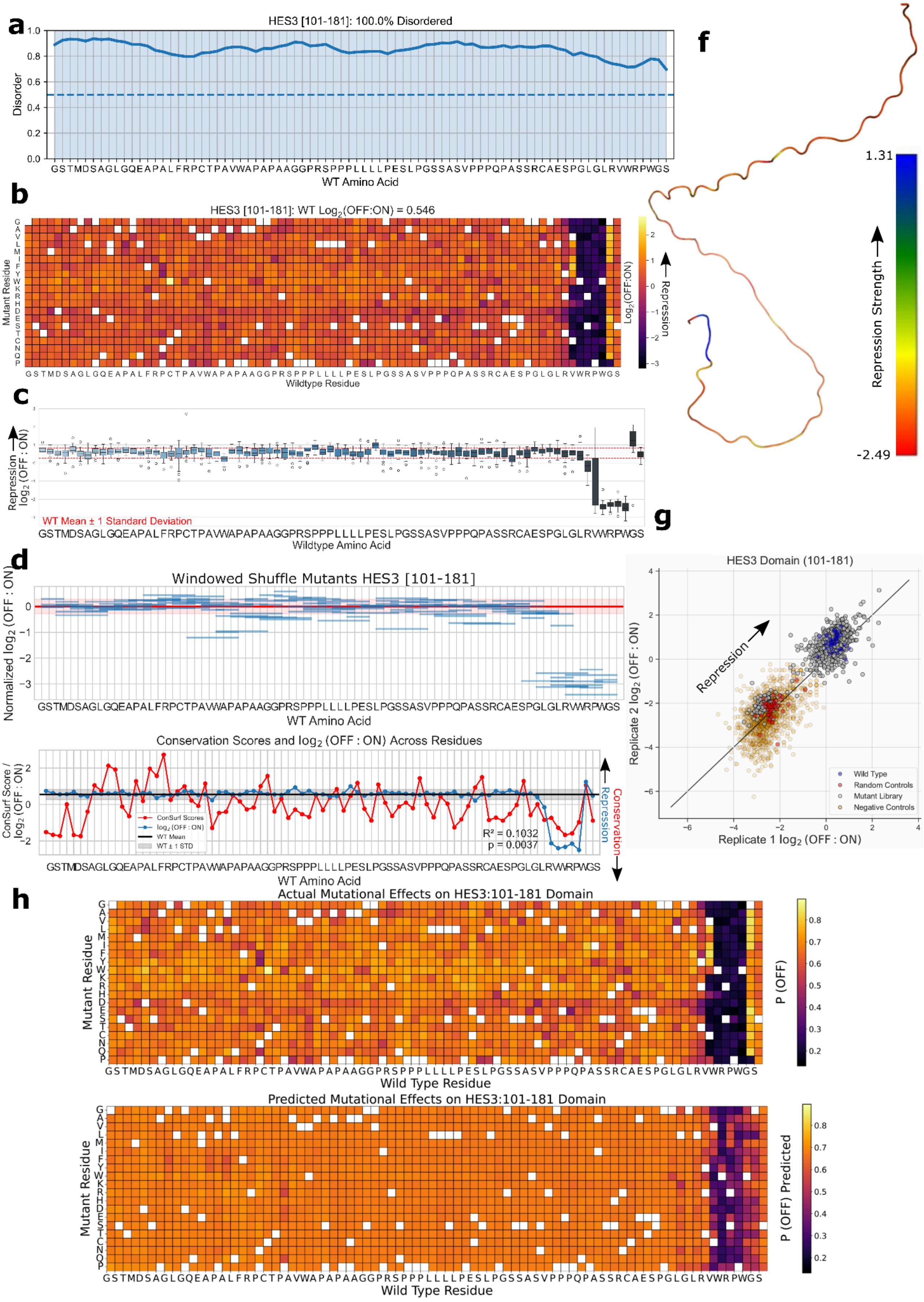

### HIPK2 [871-951] Unannotated domain

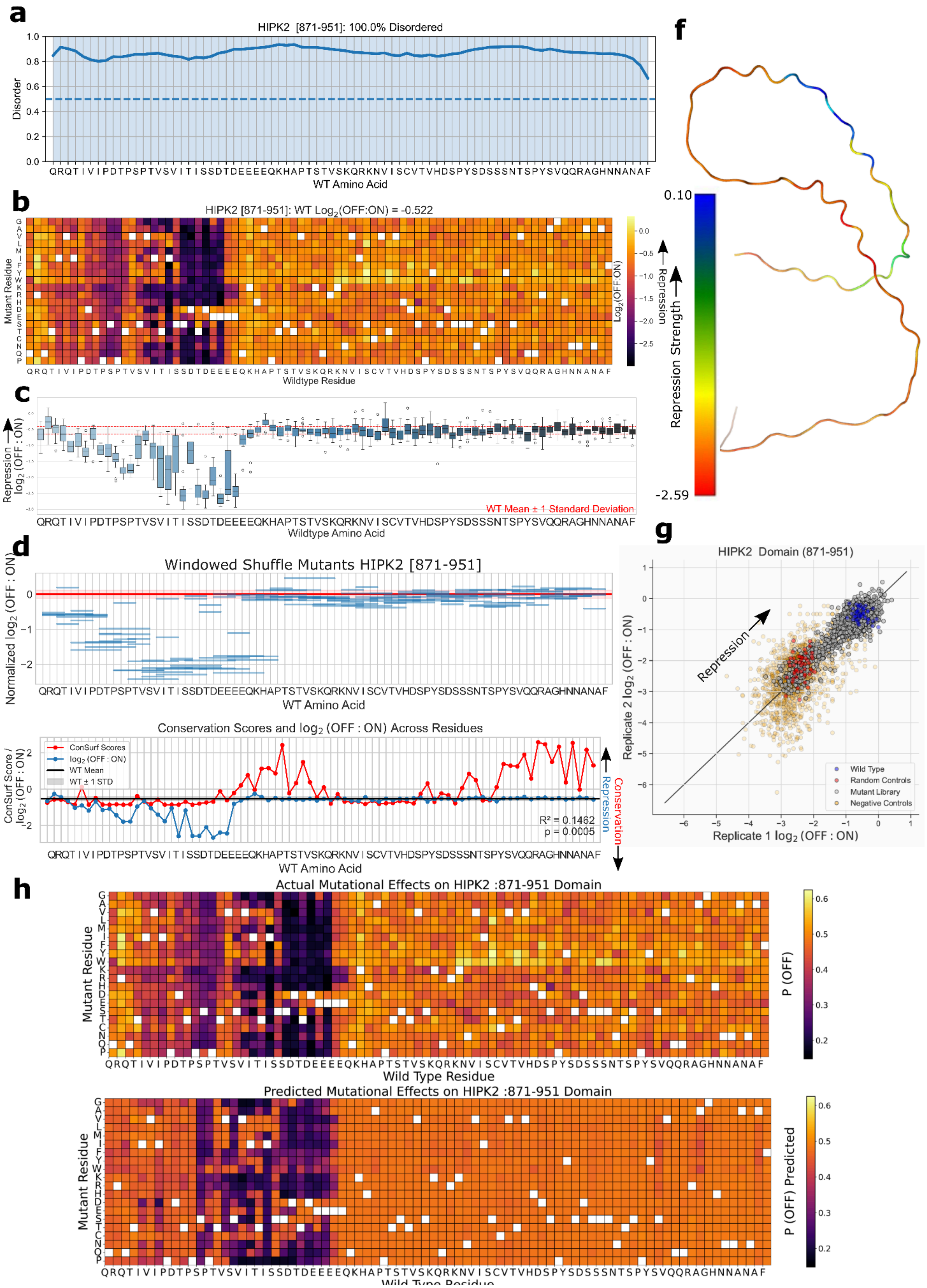

### HIVEP3 [781-861] Unannotated domain

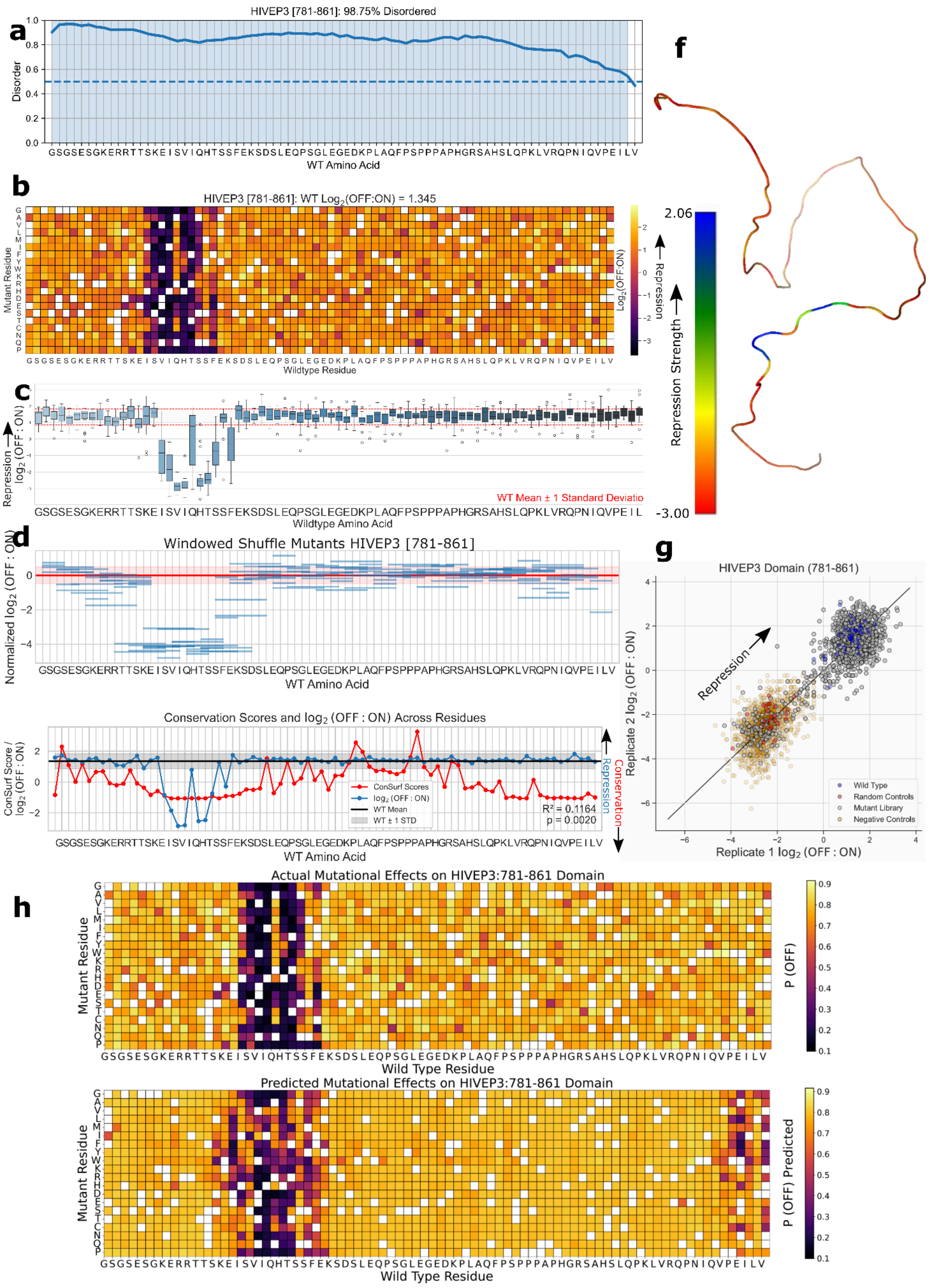

### HSF2 [131-211] Unannotated domain

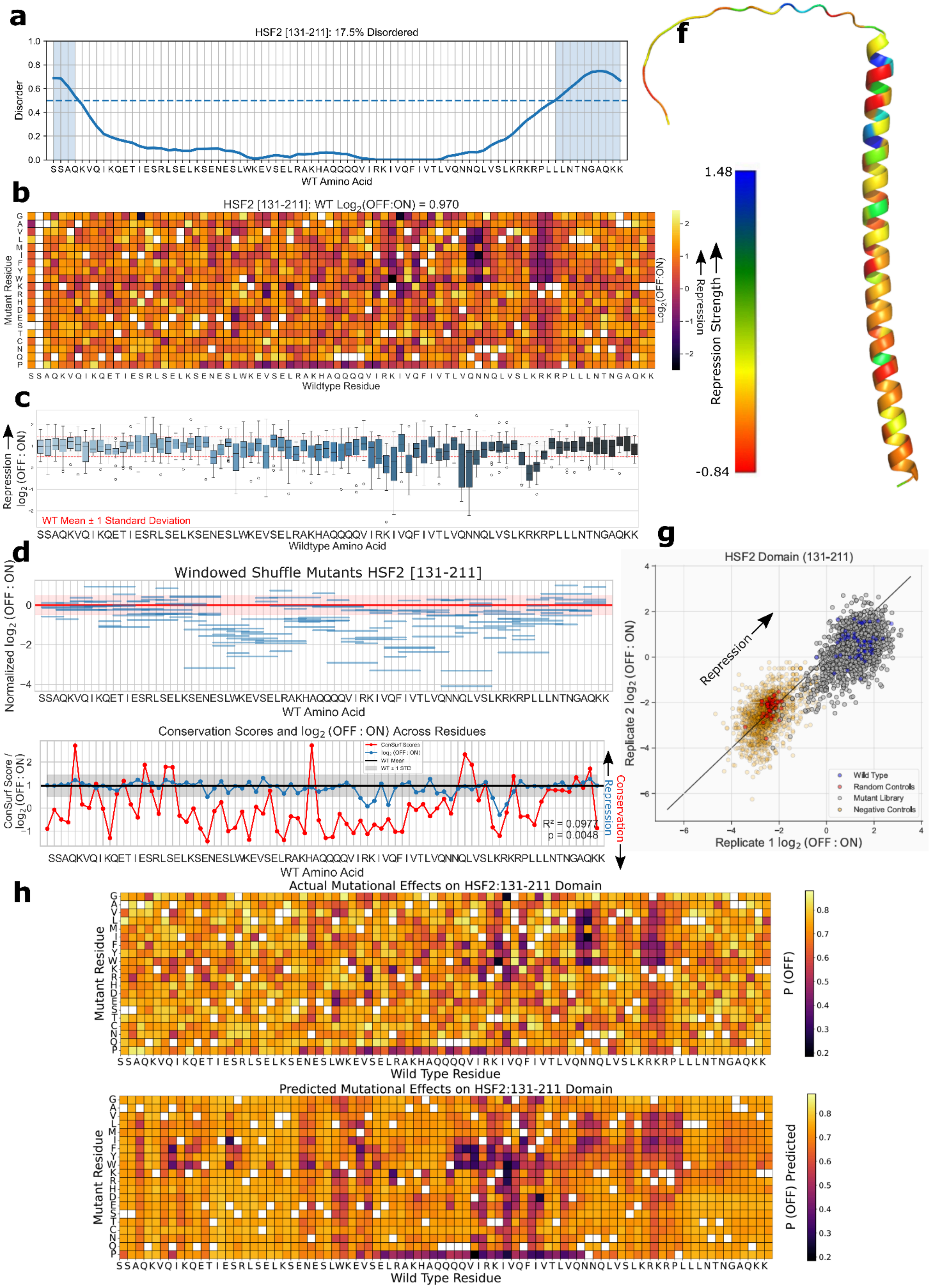

### HSTF1 [131-211] Unannotated domain

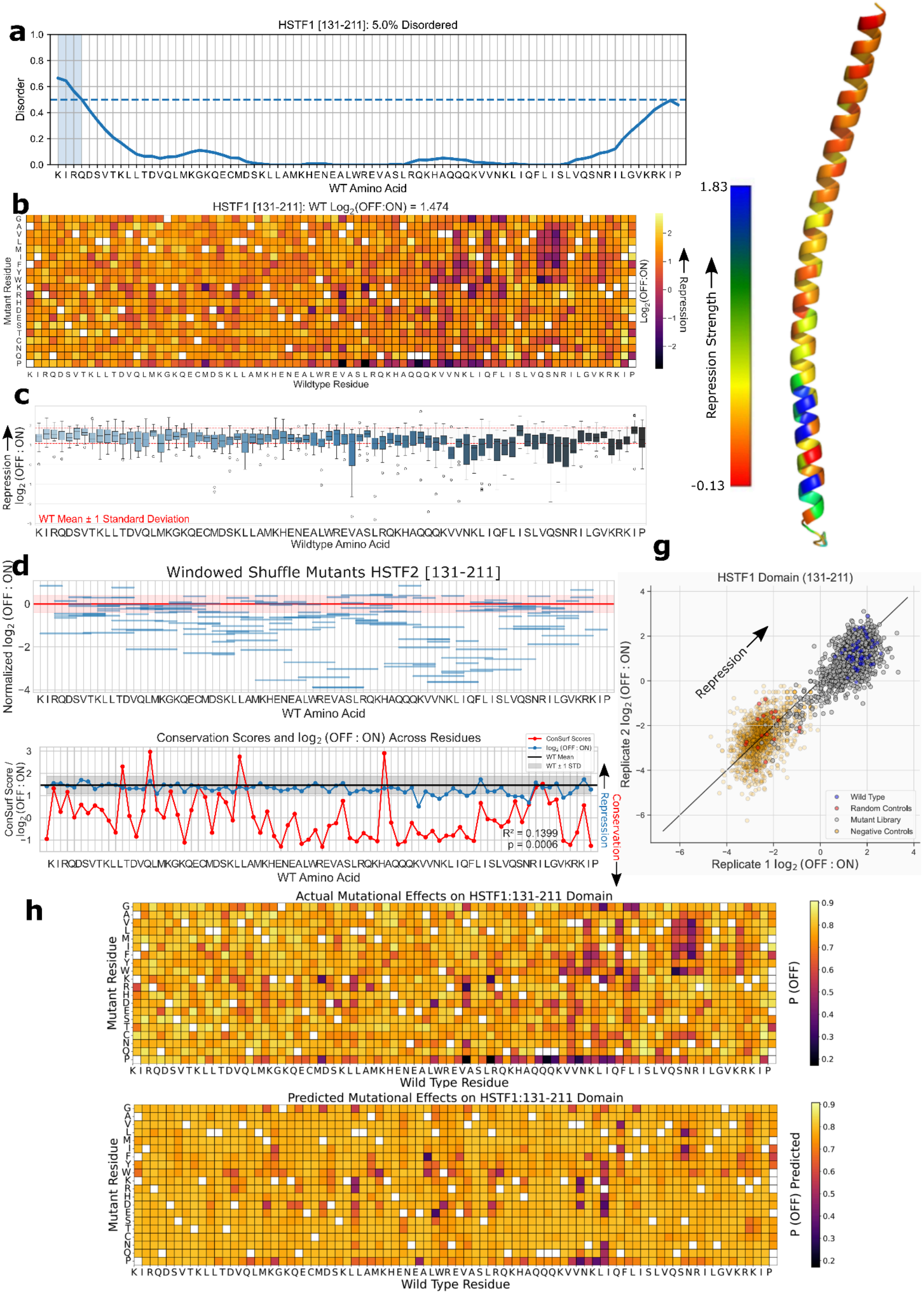

### ID1 [47-127] Helix-loop-helix DNA binding domain

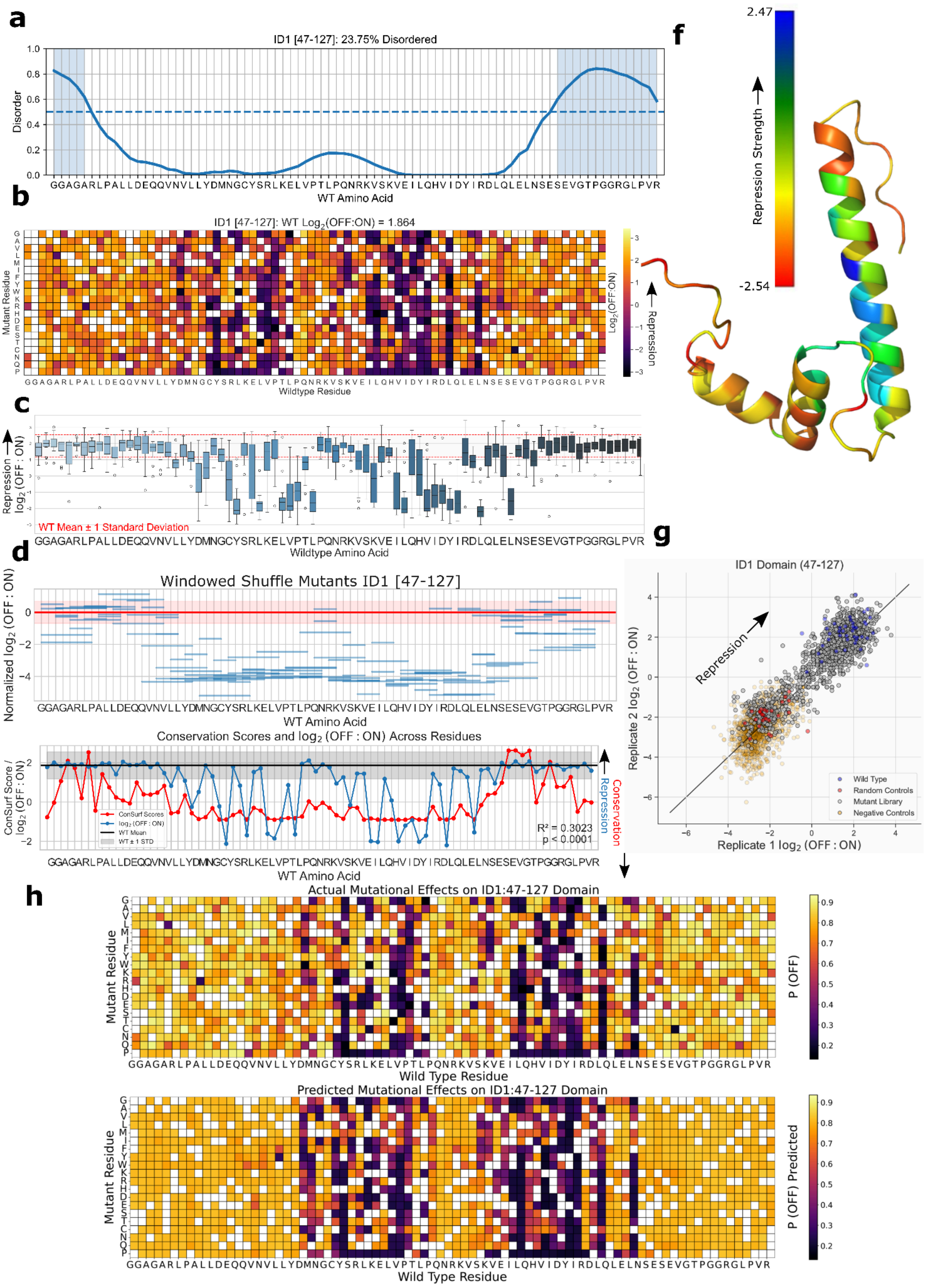

### ID2 [16-96] Helix-loop-helix DNA binding domain

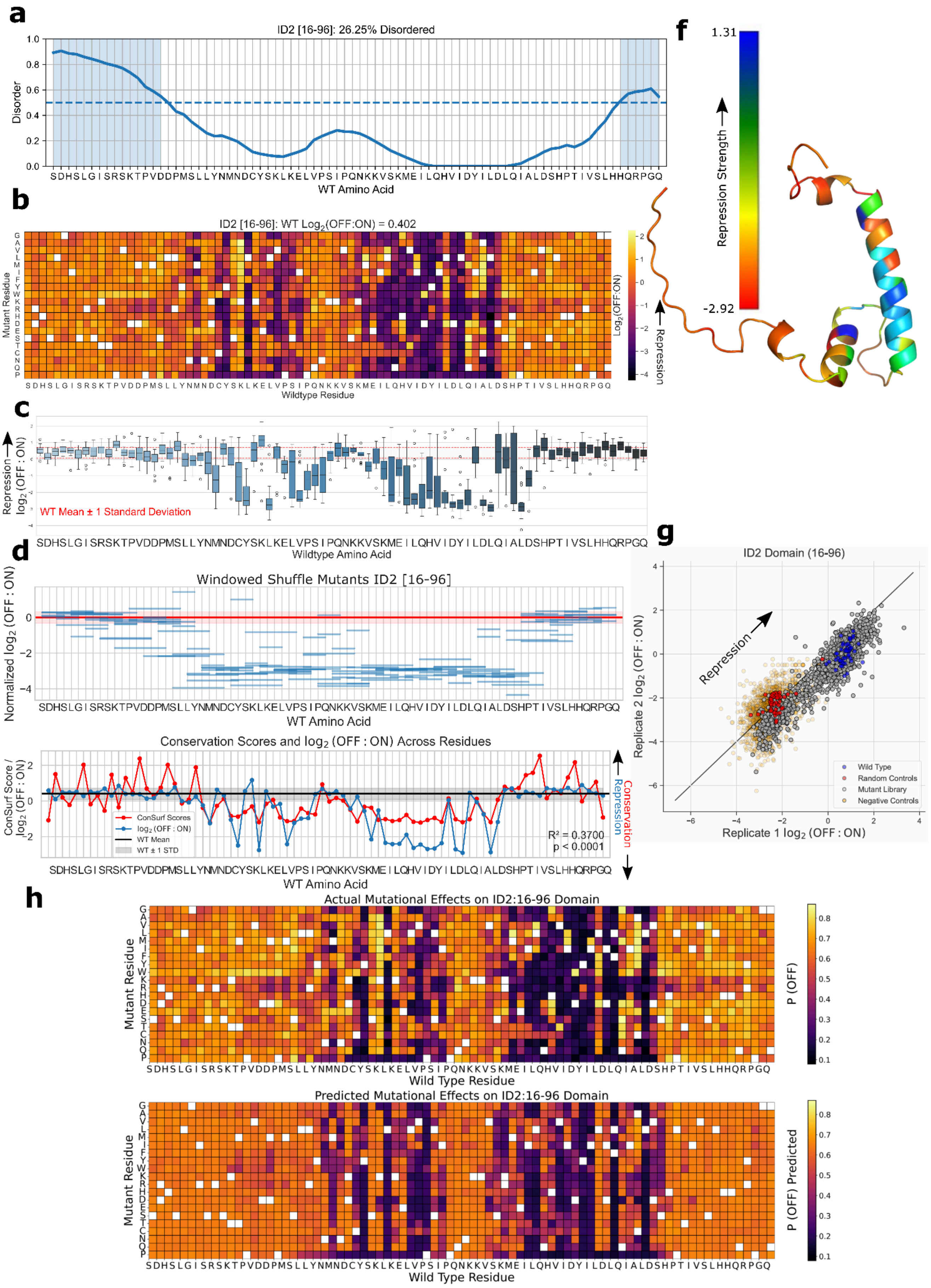

### ID3 [21-101] Helix-loop-helix DNA binding domain

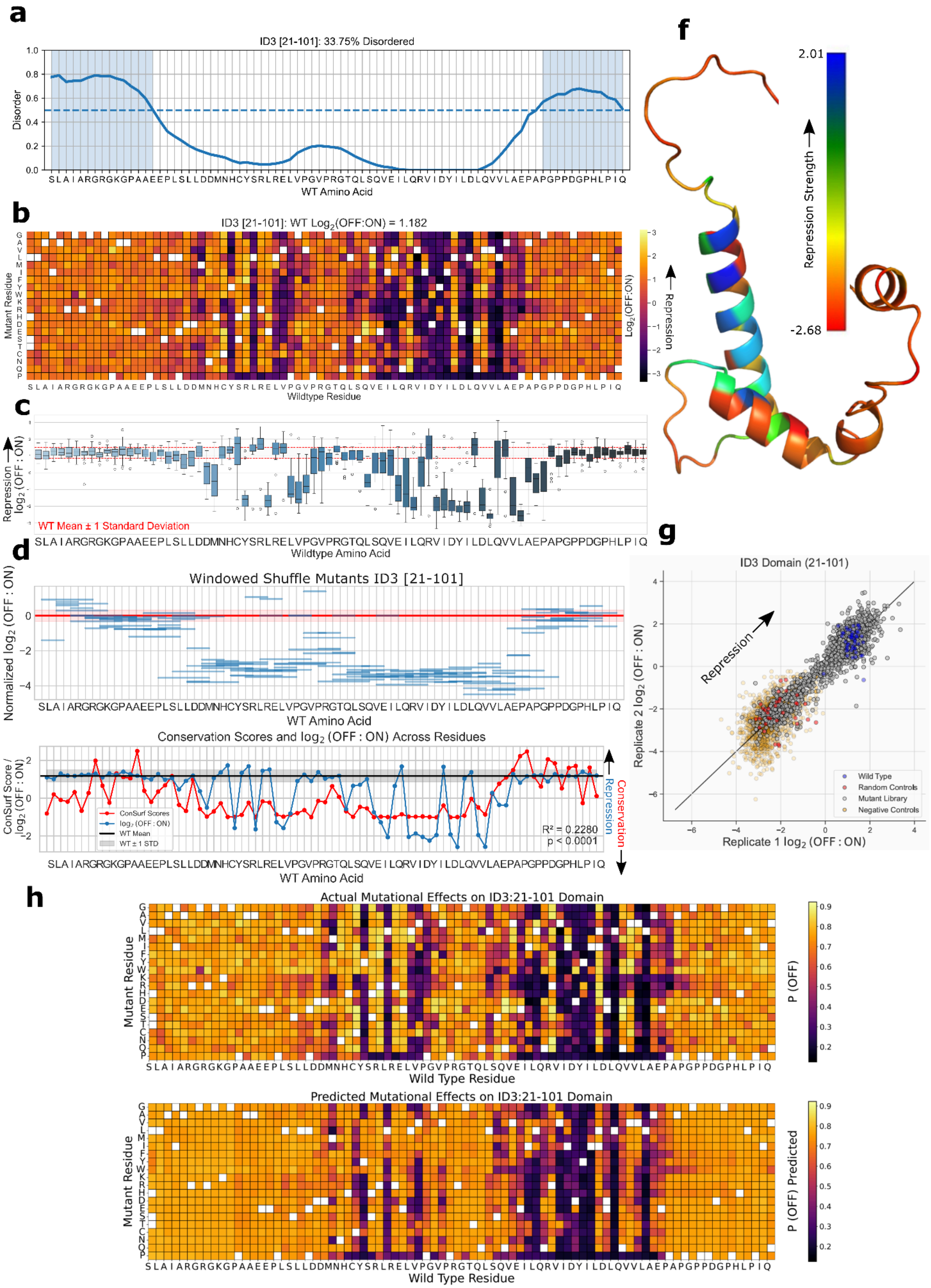

### IKZF5 [340-420] Zinc finger C2H2 type domain profile

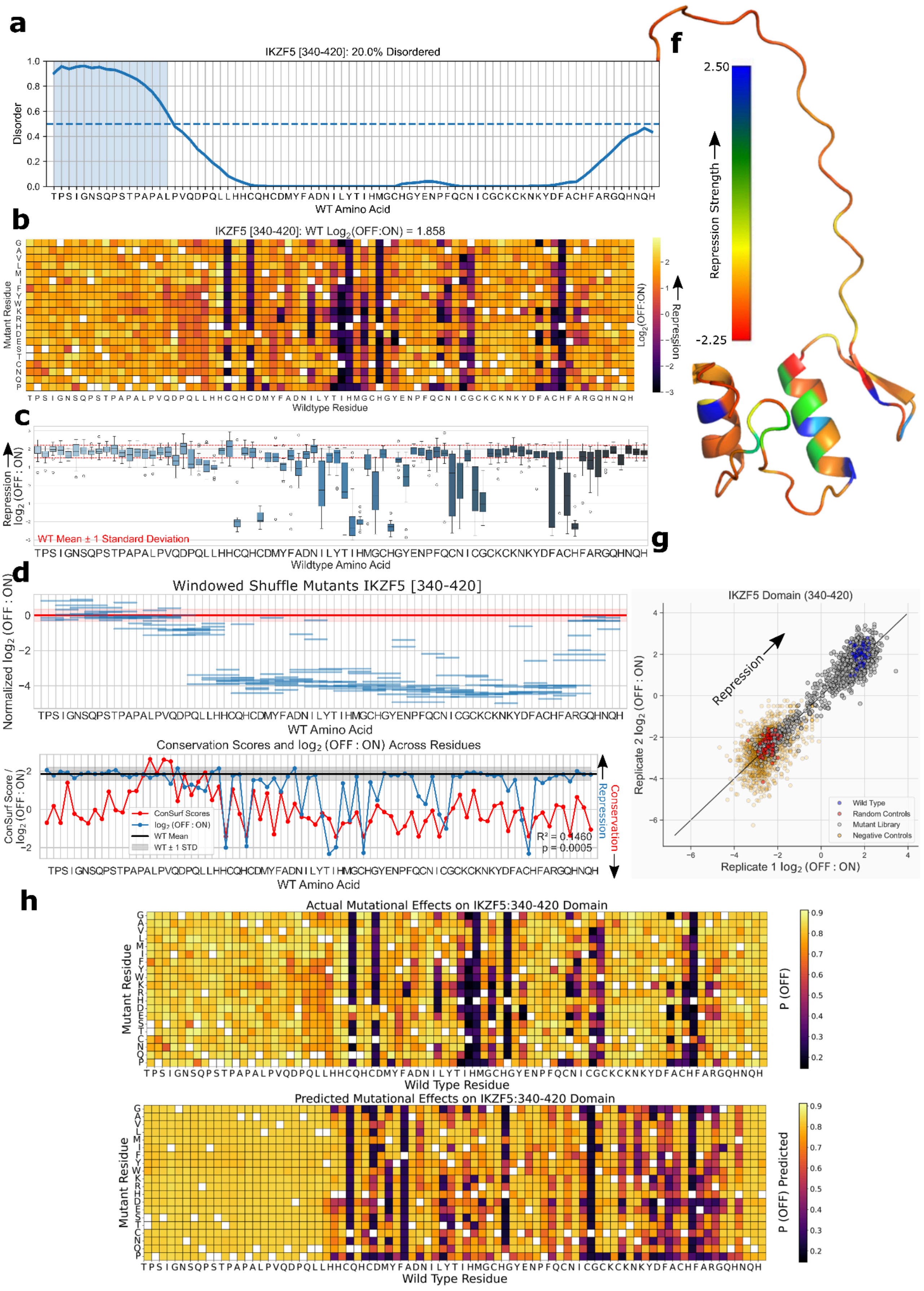

### IRF2BP1 [1-81] Interferon regulatory factor 2-binding protein zinc finger

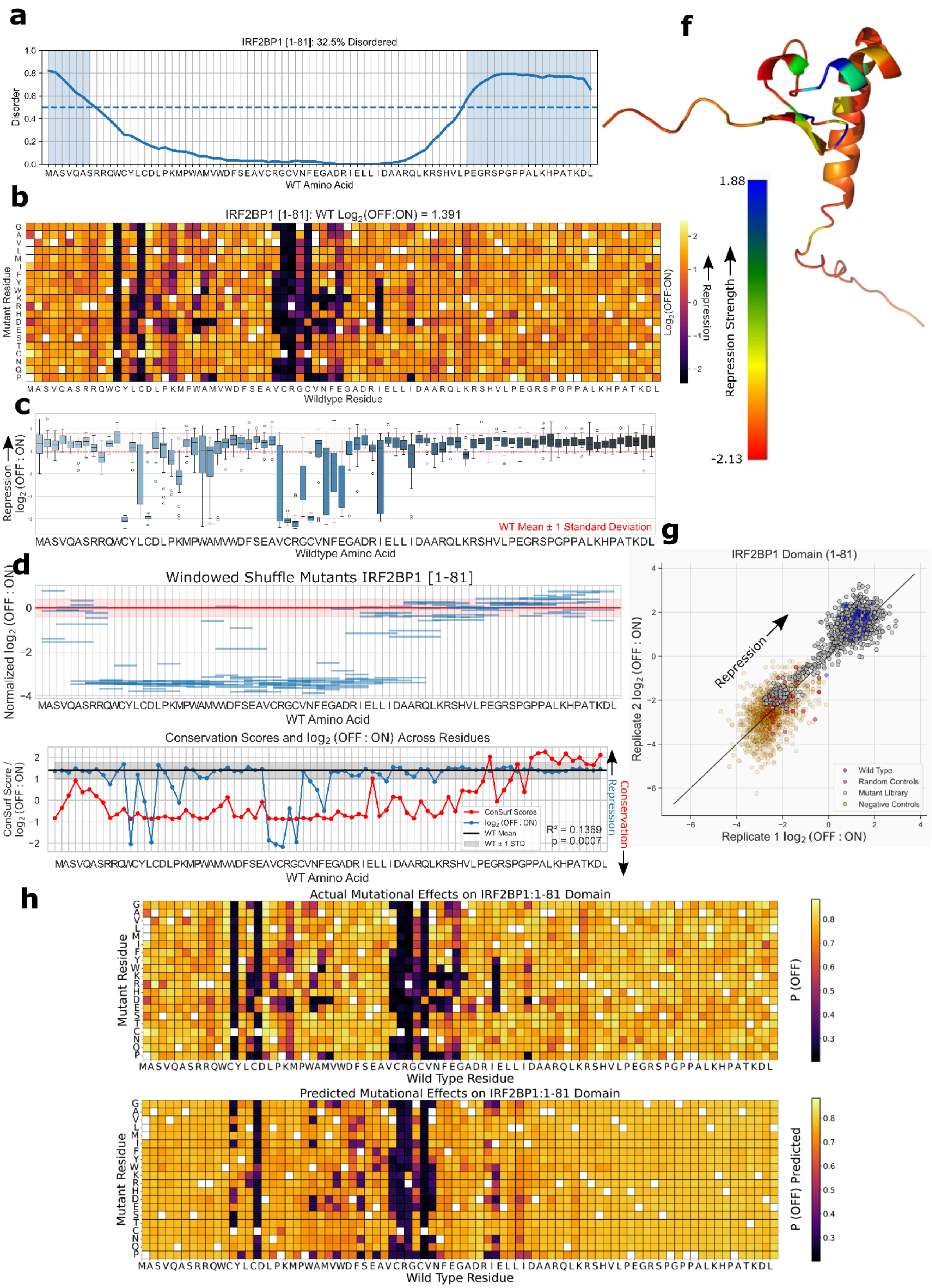

### KDM5B [261-341] PHD-finger domain

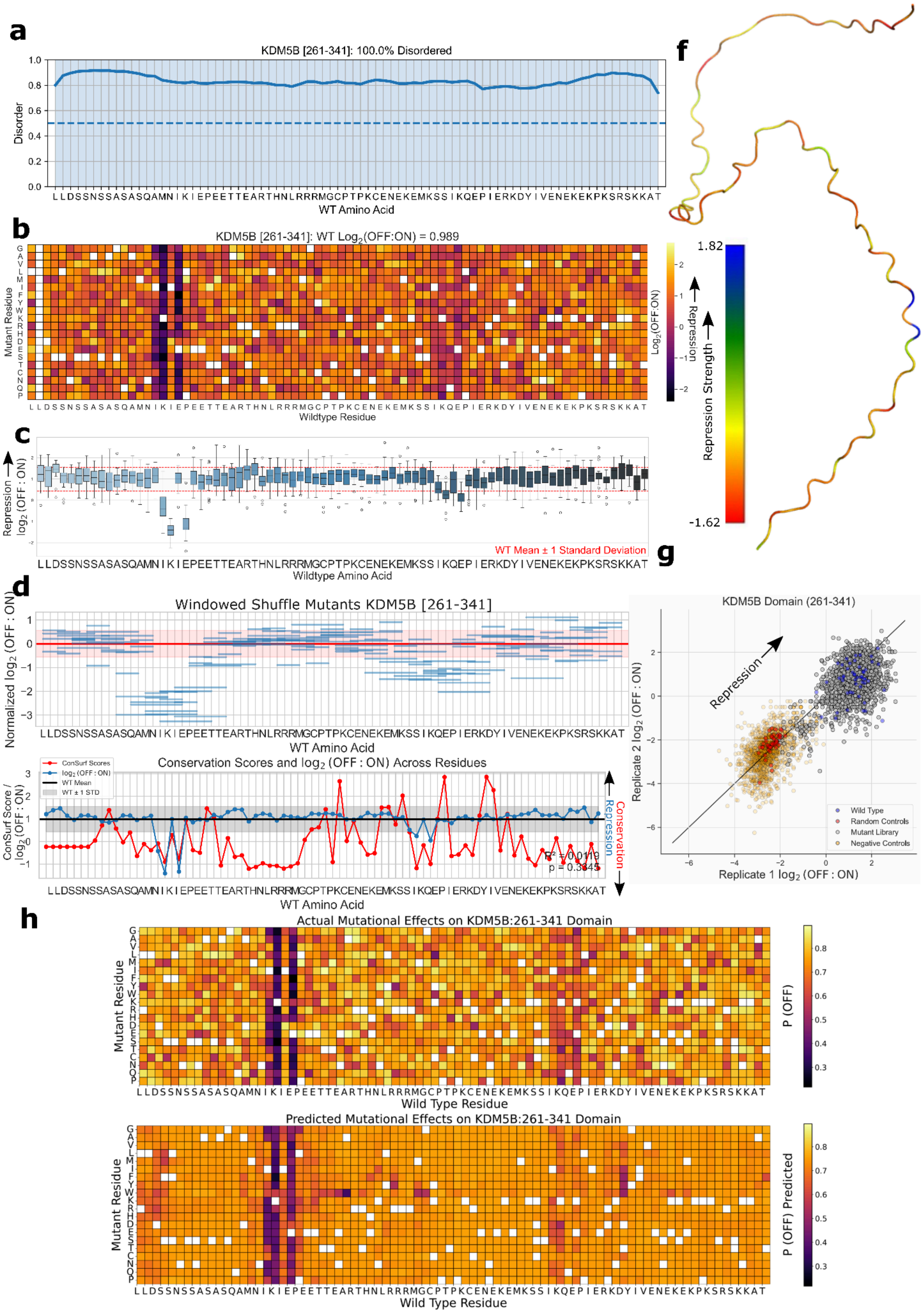

### KLF10 [132-212] Unannotated domain

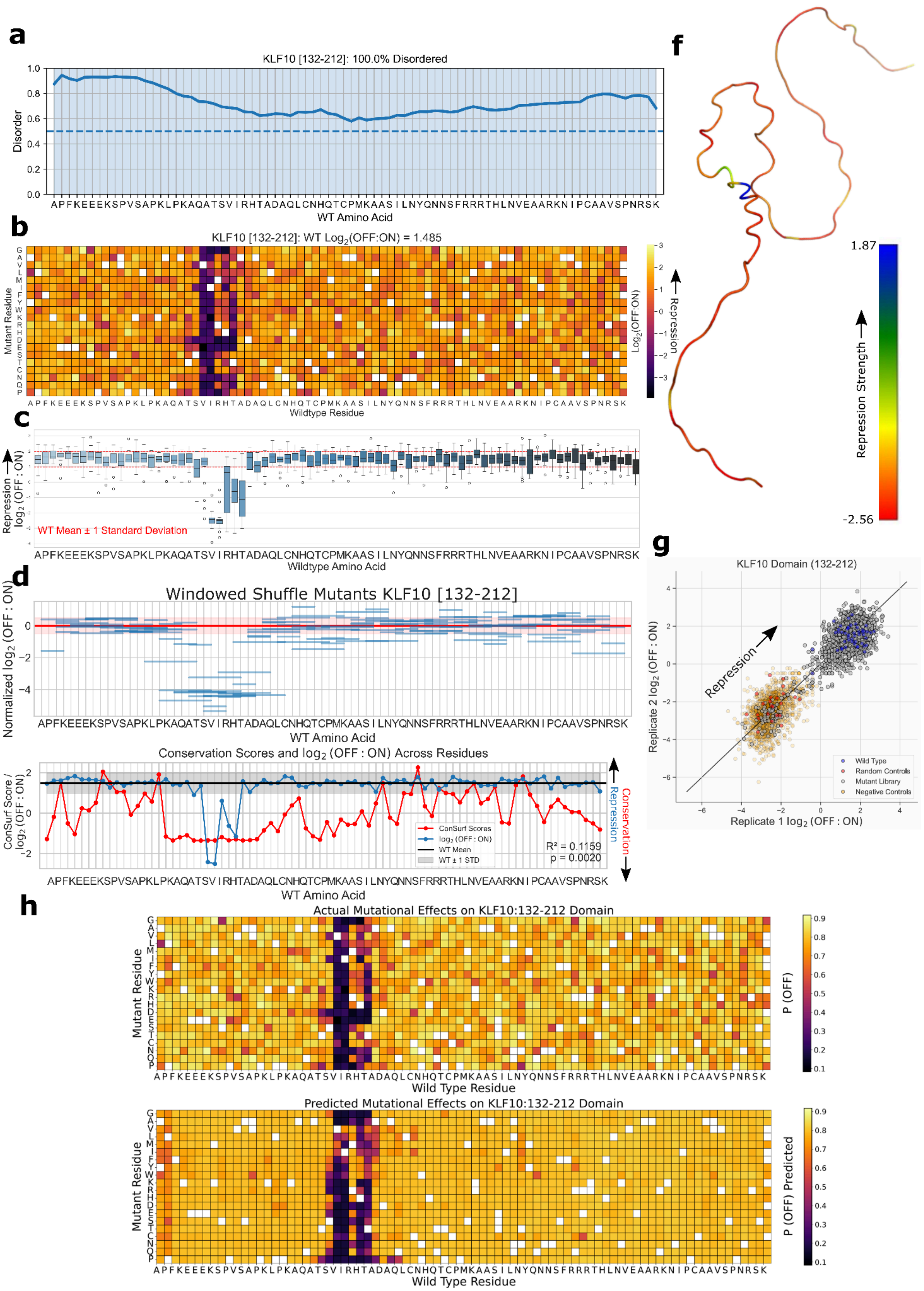

### MBD1 [1-81] Methyl-CpG binding domain

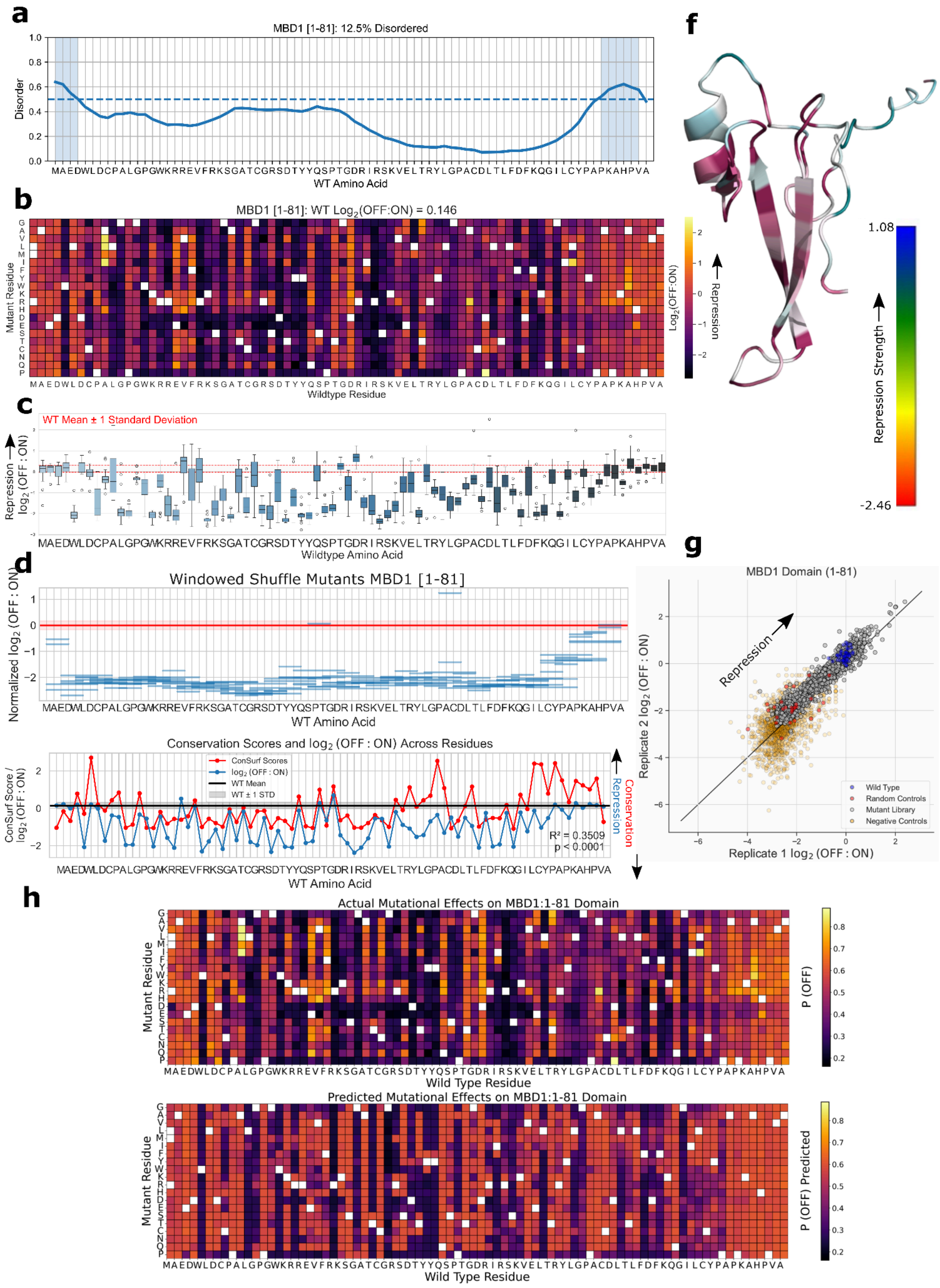

### MBD1 [526-606] Unannotated domain

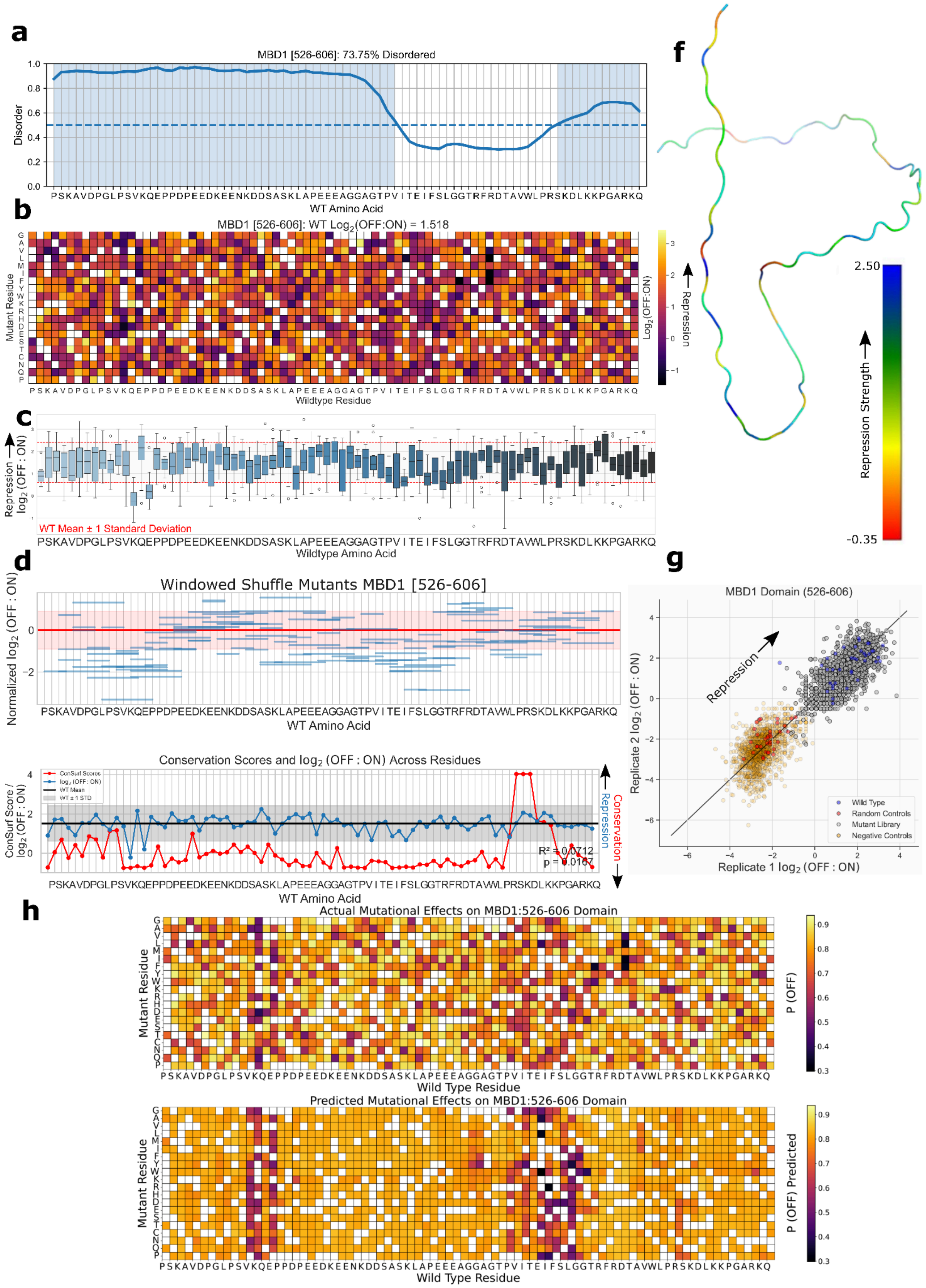

### MECP2 [241-321] Unannotated domain

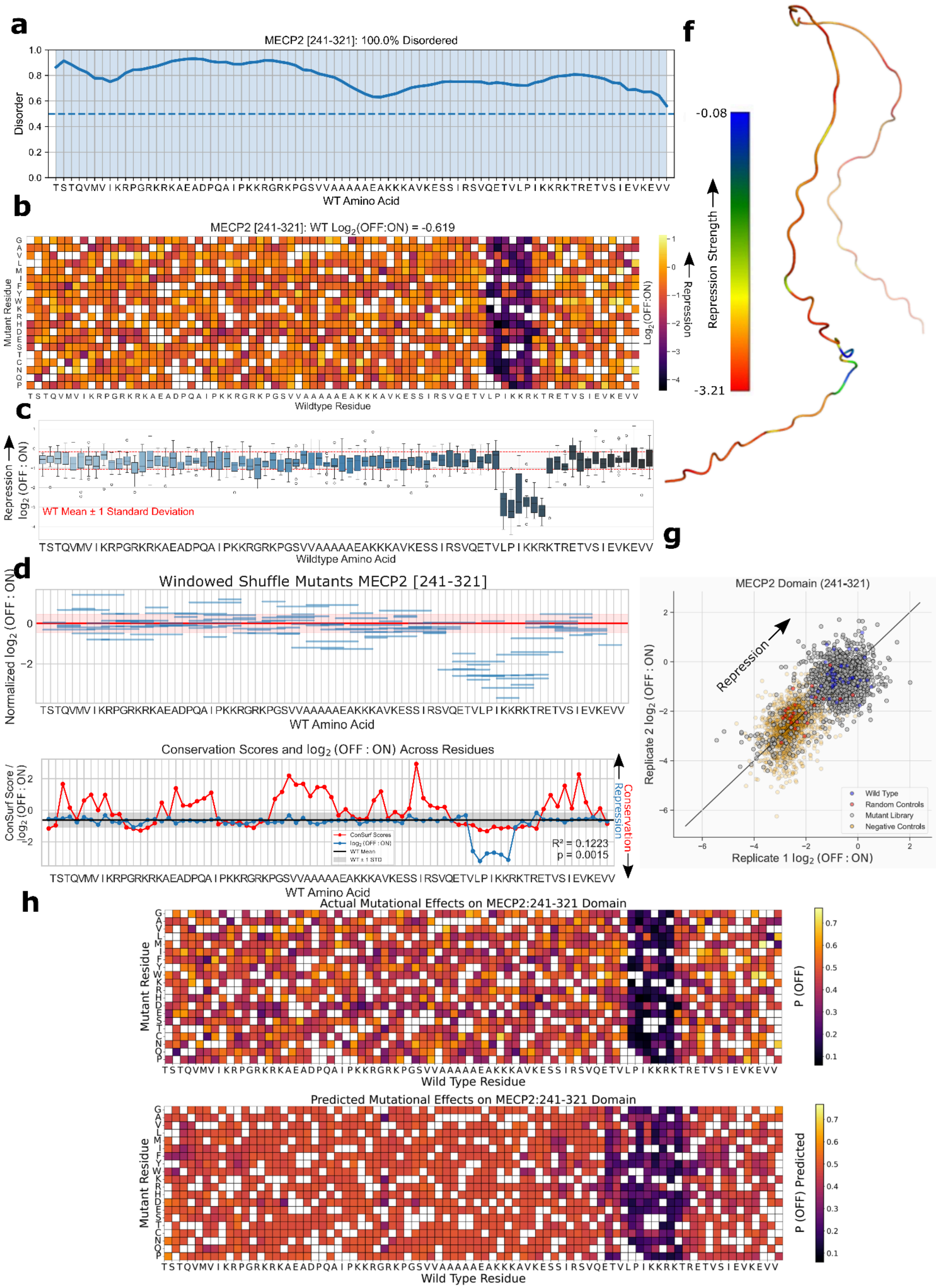

### MGA [341-421] Unannotated domain

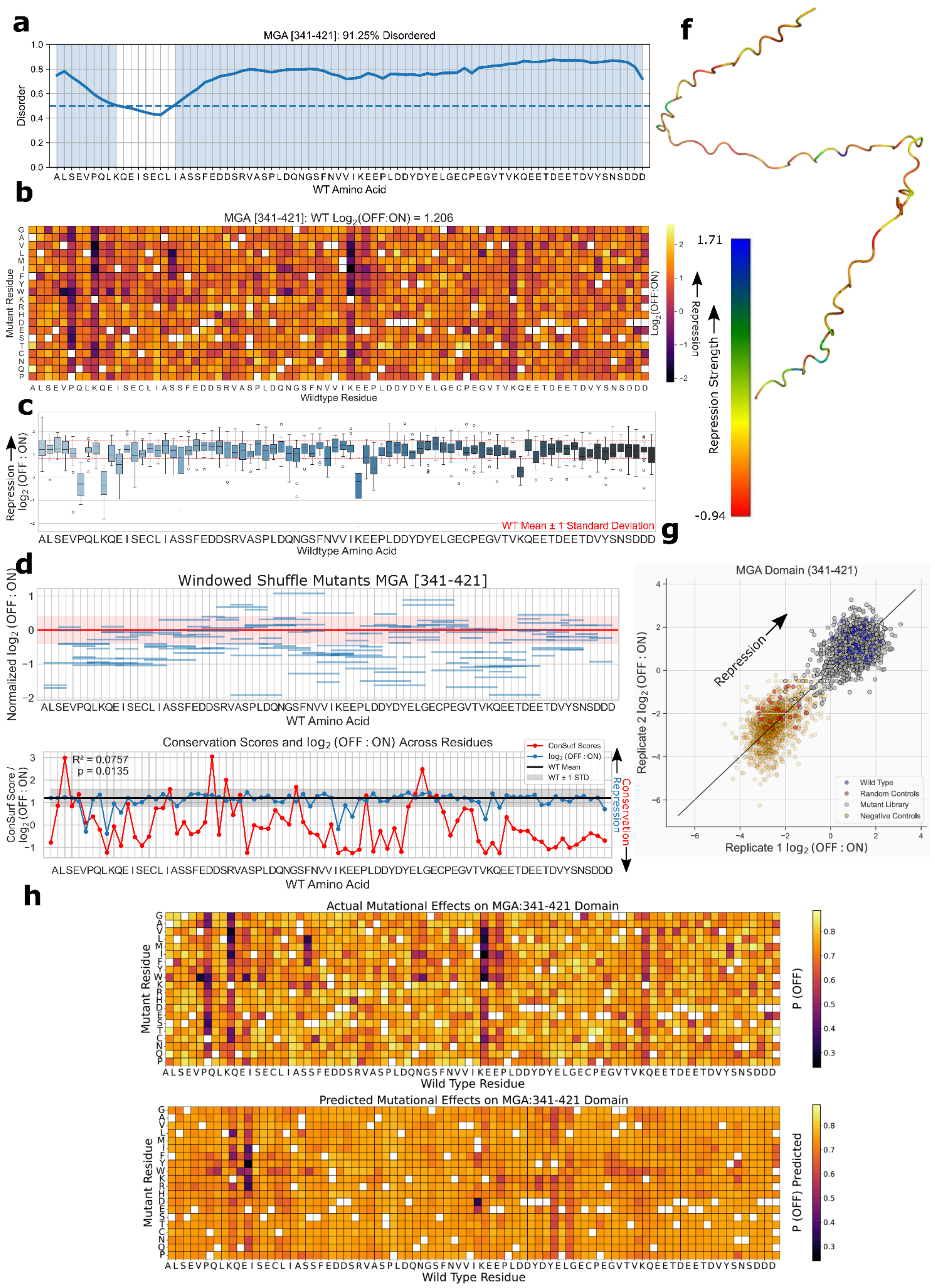

### MGA [2590-2670] Unannotated domain

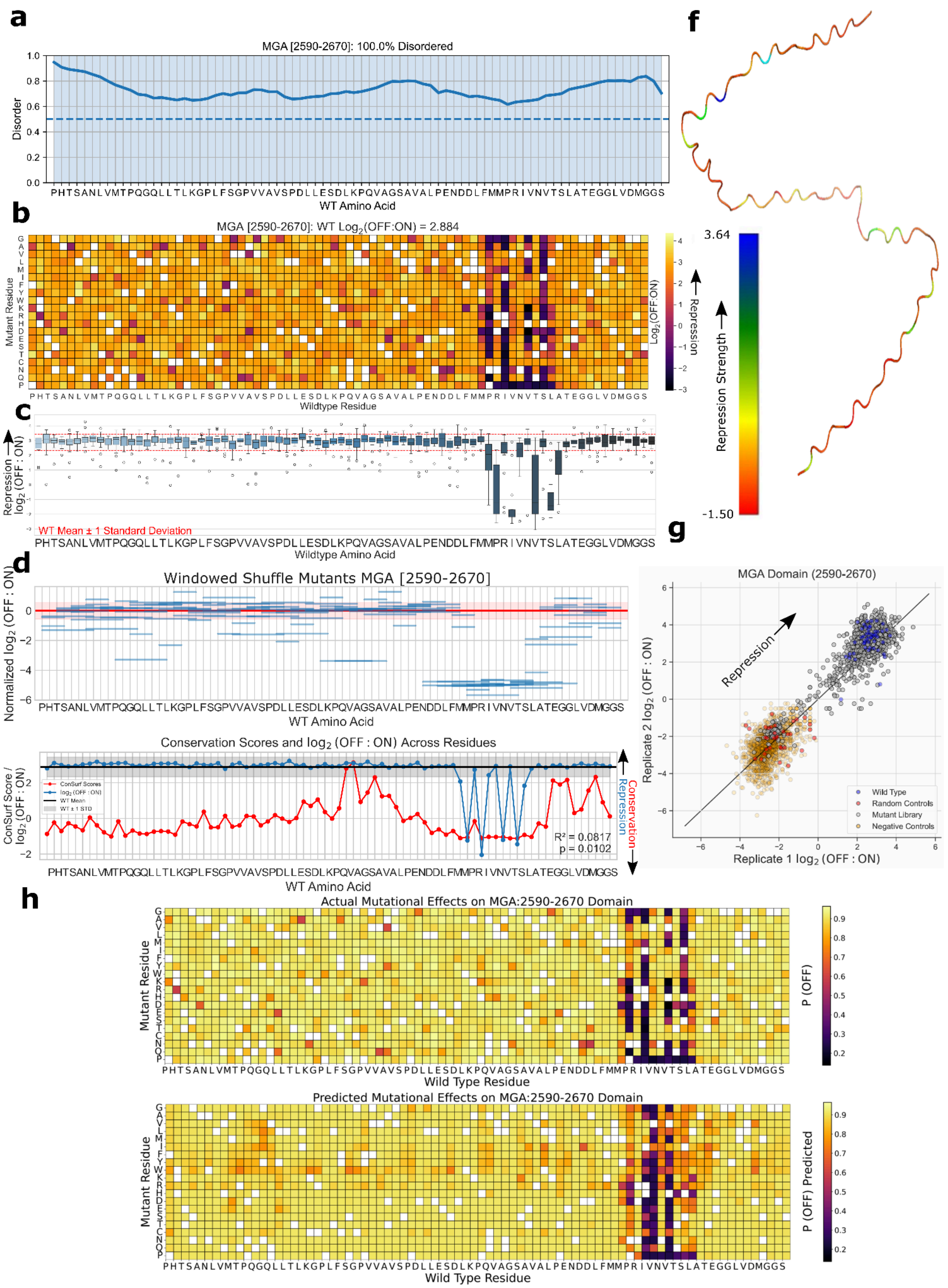

### MPHOSPH8 [45-125] Chromo (CHRromatin Organisation MOdifier) domain

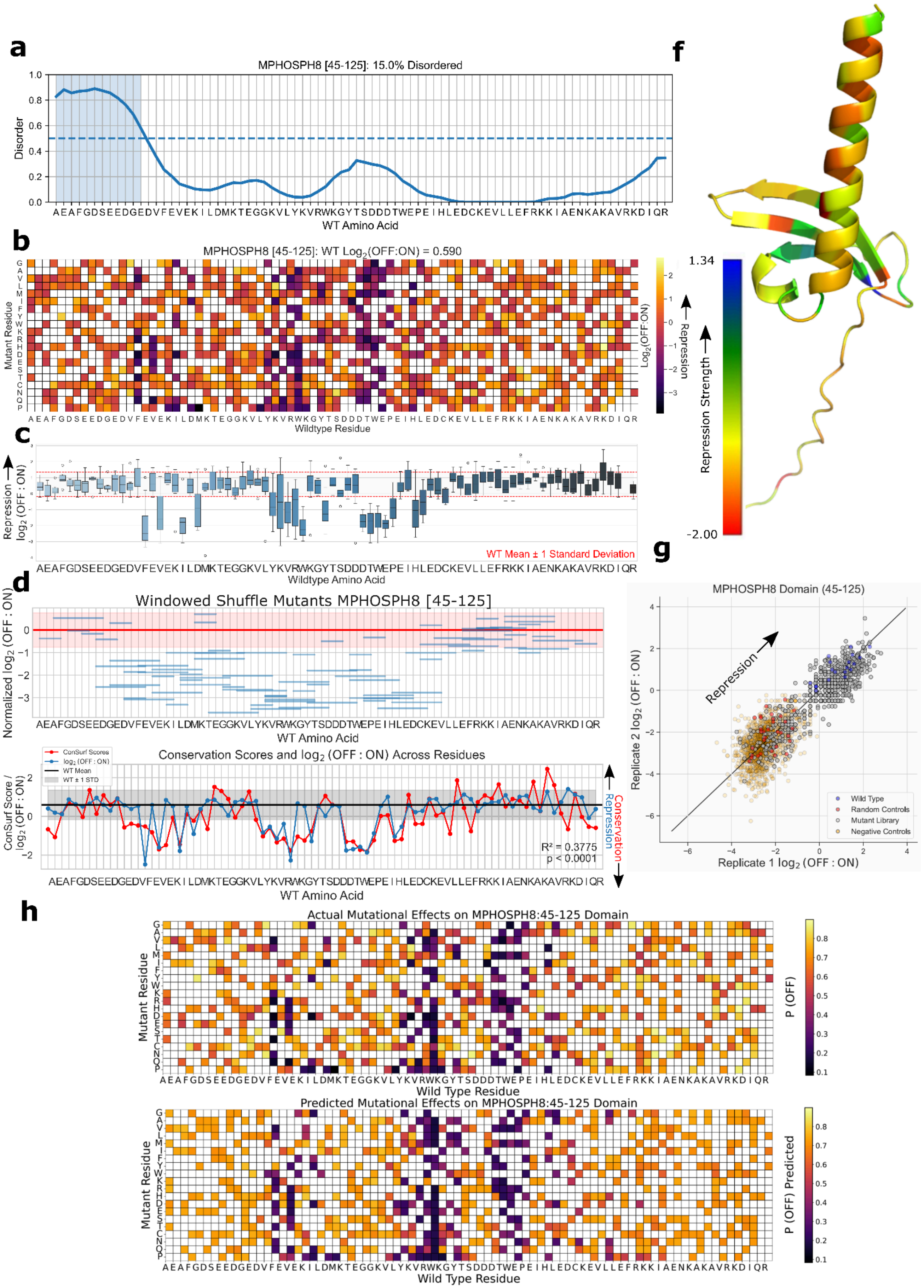

### NEUROG2 [100-180] Helix-loop-helix DNA binding domain

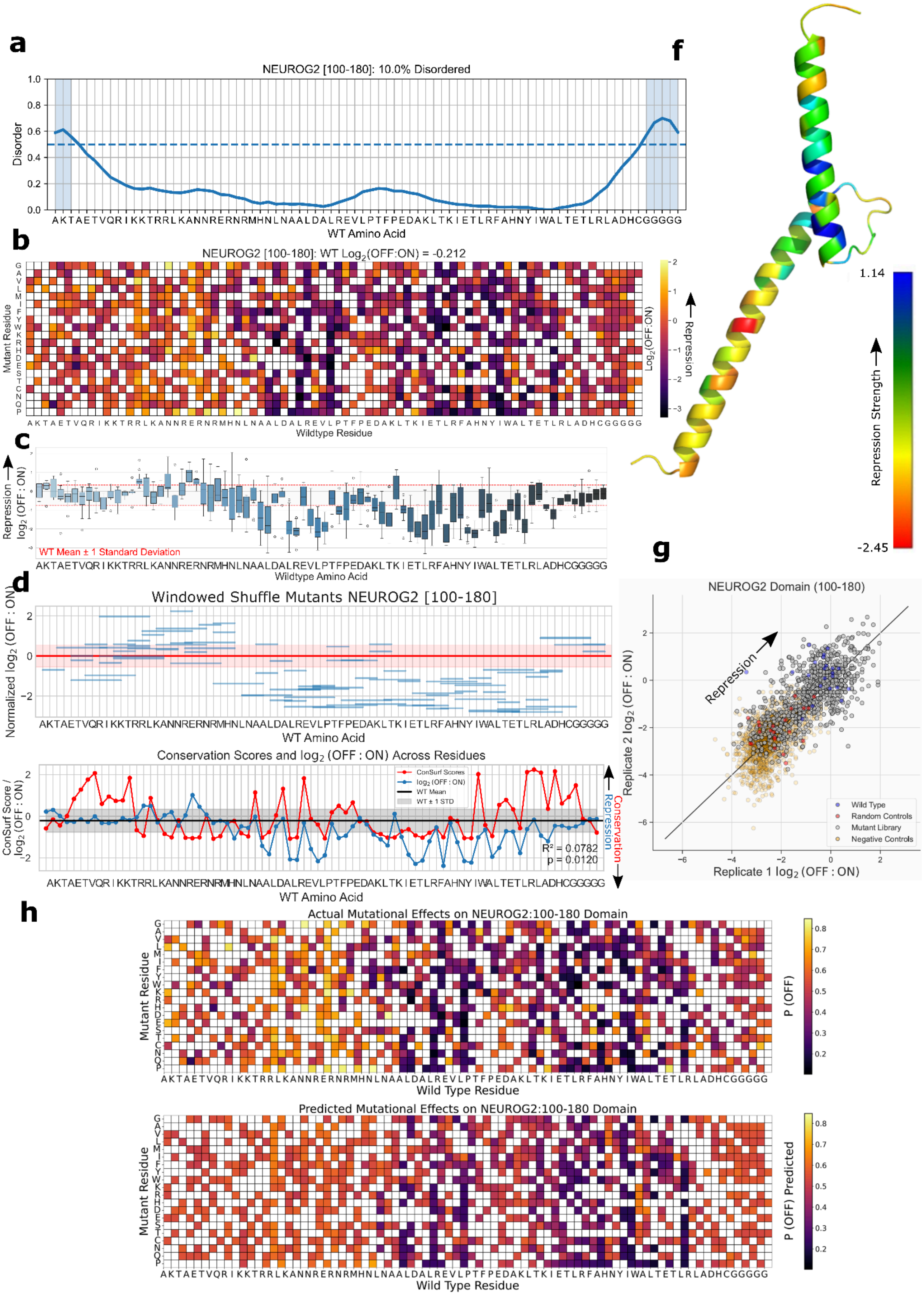

### PCGF2 [2-82] Zinc finger RING-type profile domain

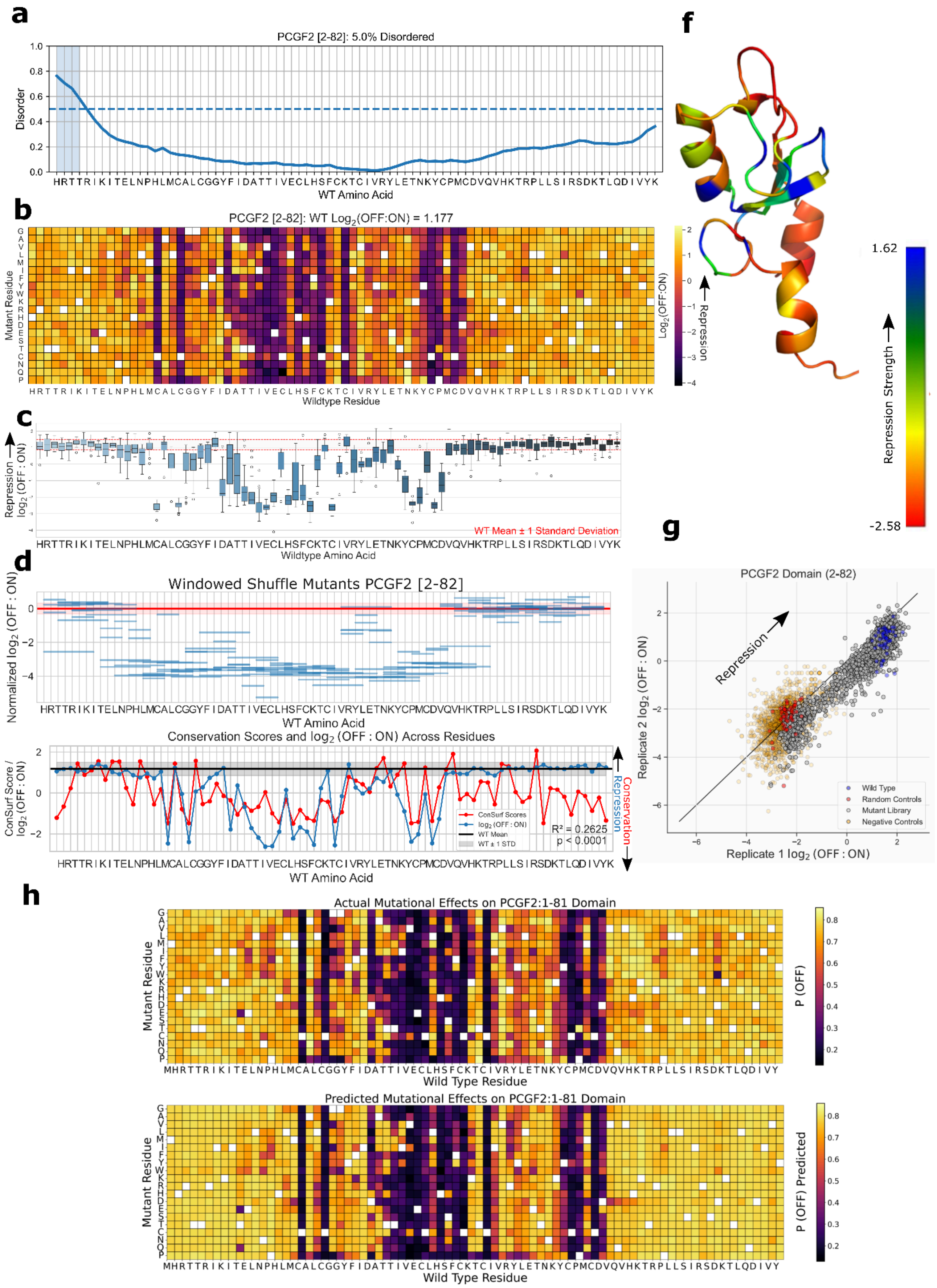

### PCGF2 [1-81] Zinc finger RING-type profile domain

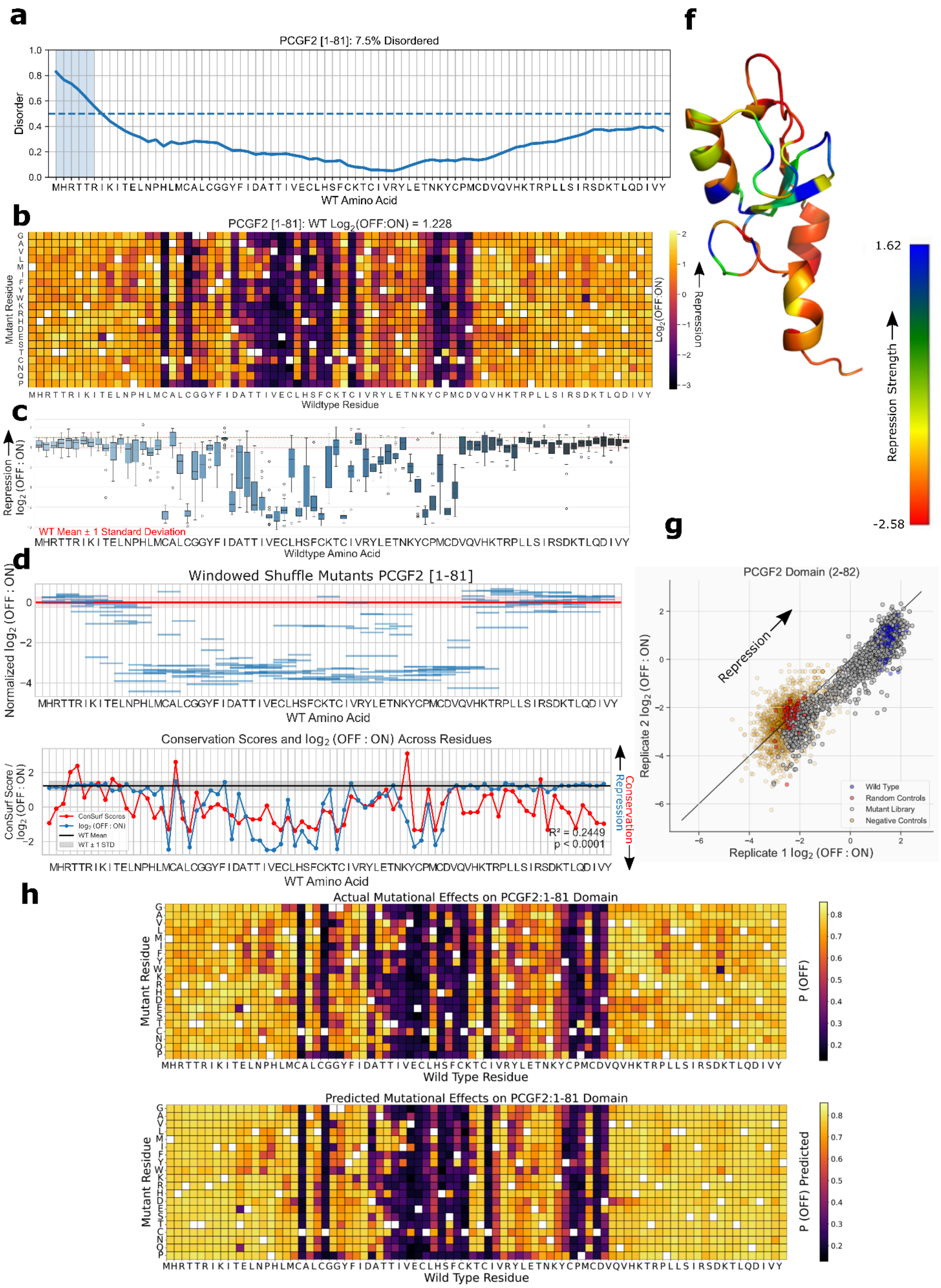

### REST [1011-1091] Zinc finger C2H2 type domain profile

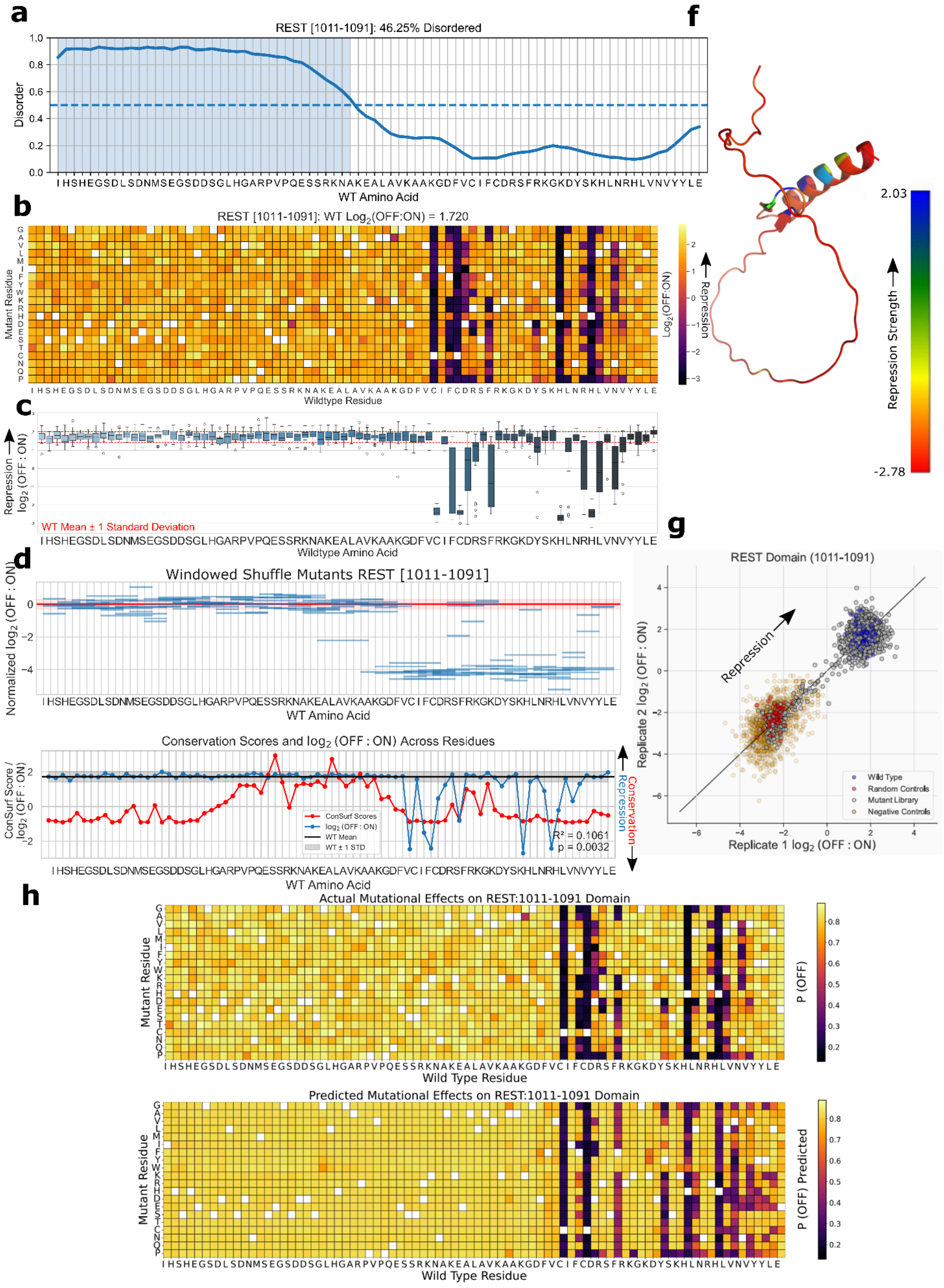

### RYBP [122-202] Yaf2/RYBP C-terminal binding motif domain

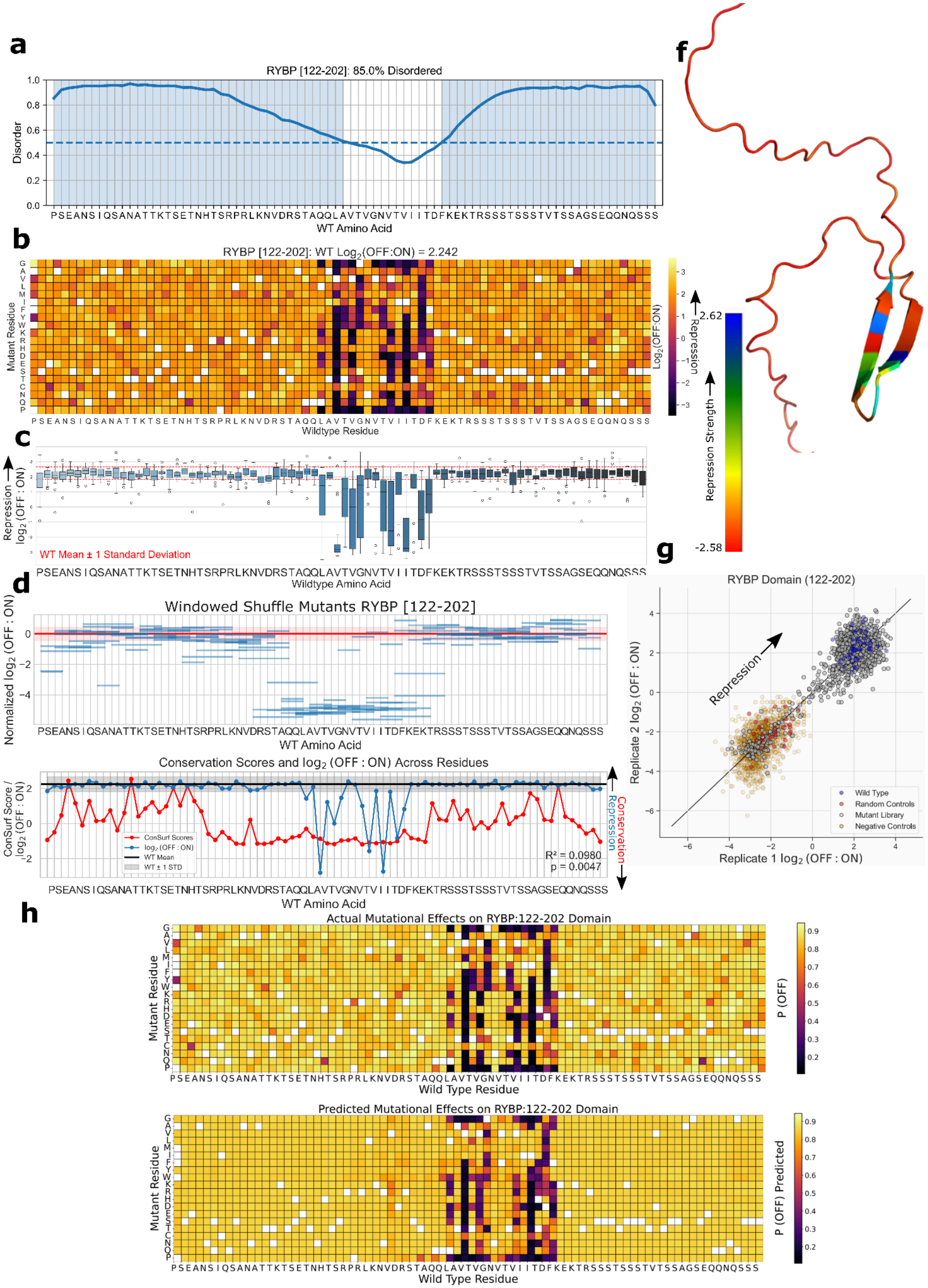

RYBP [121-201] Yaf2/RYBP C-terminal binding motif domain

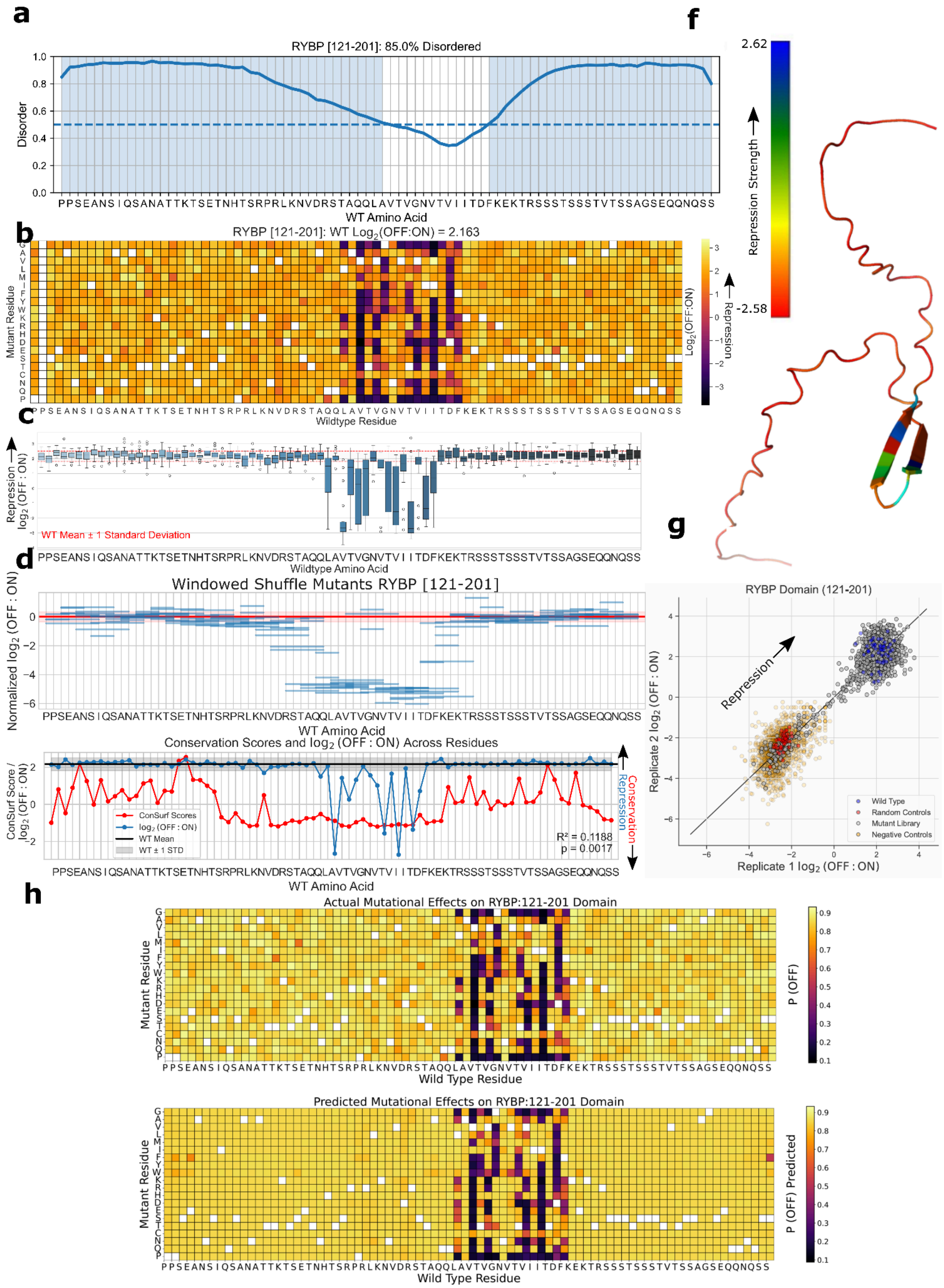

### SCML2 [621-701] SAM domain (Sterile alpha motif) domain

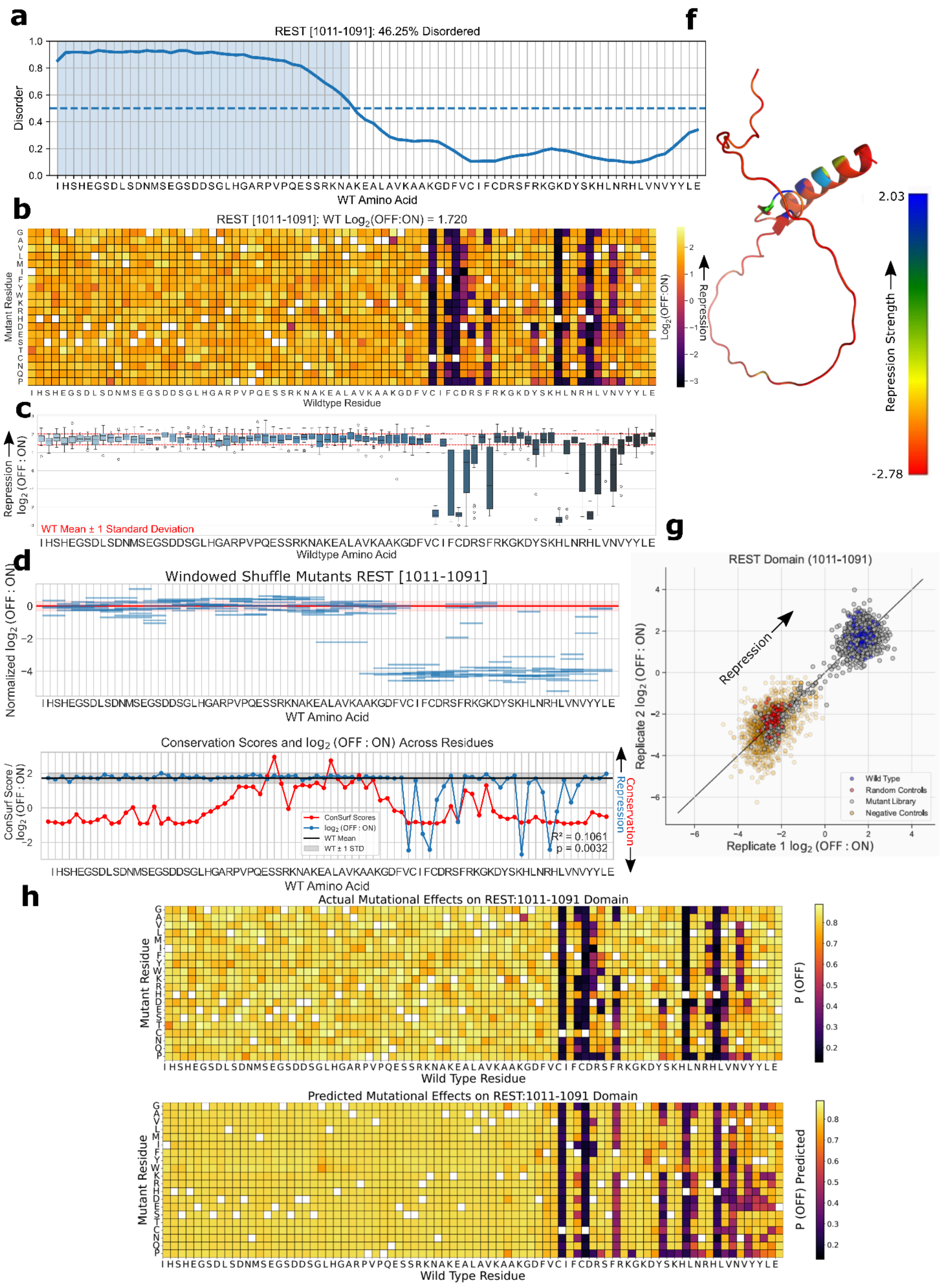

### SCX [63-143] Helix-loop-helix DNA binding domain

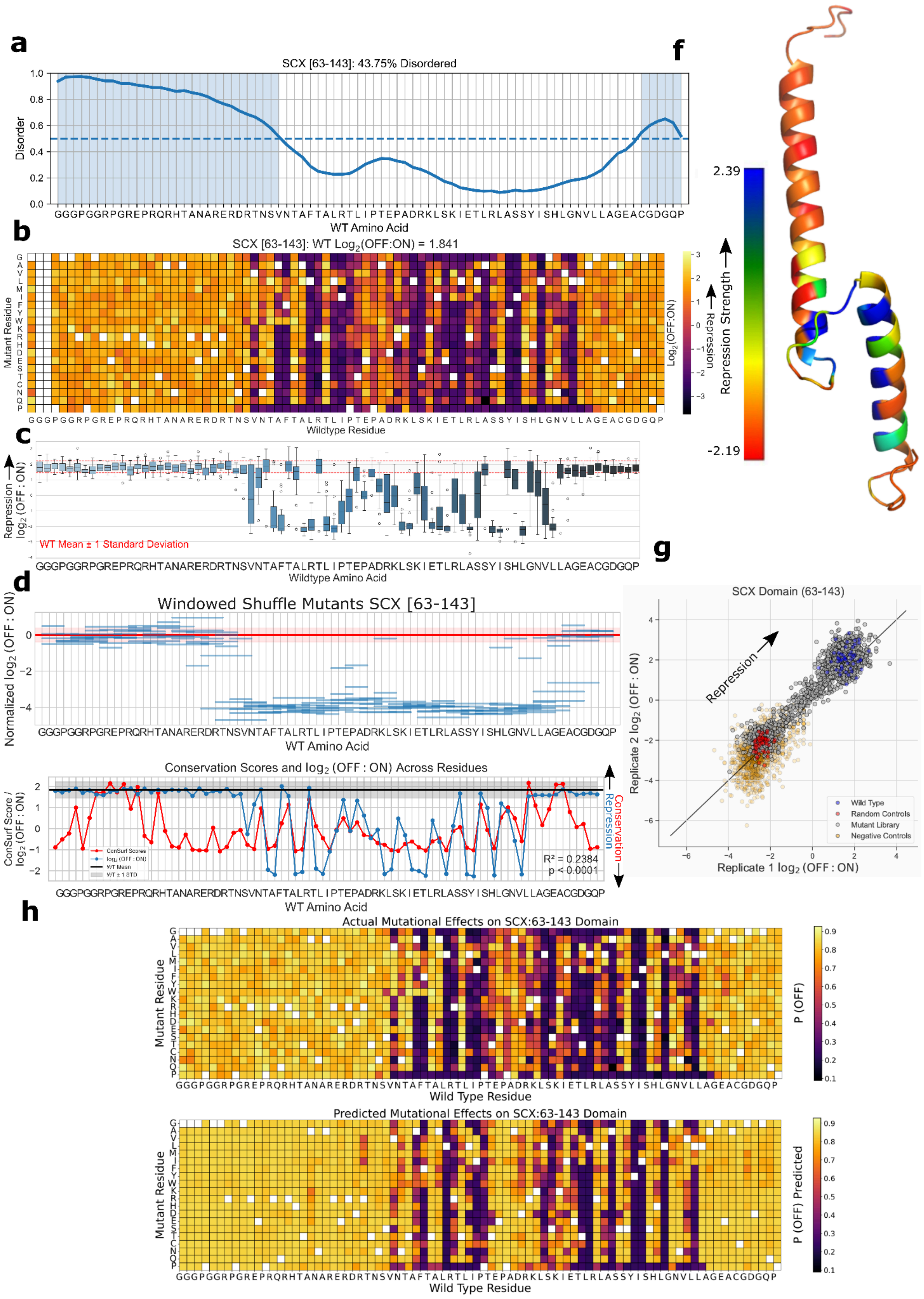

### SIN3B [31-111] Paired amphipathic helix repeat domain

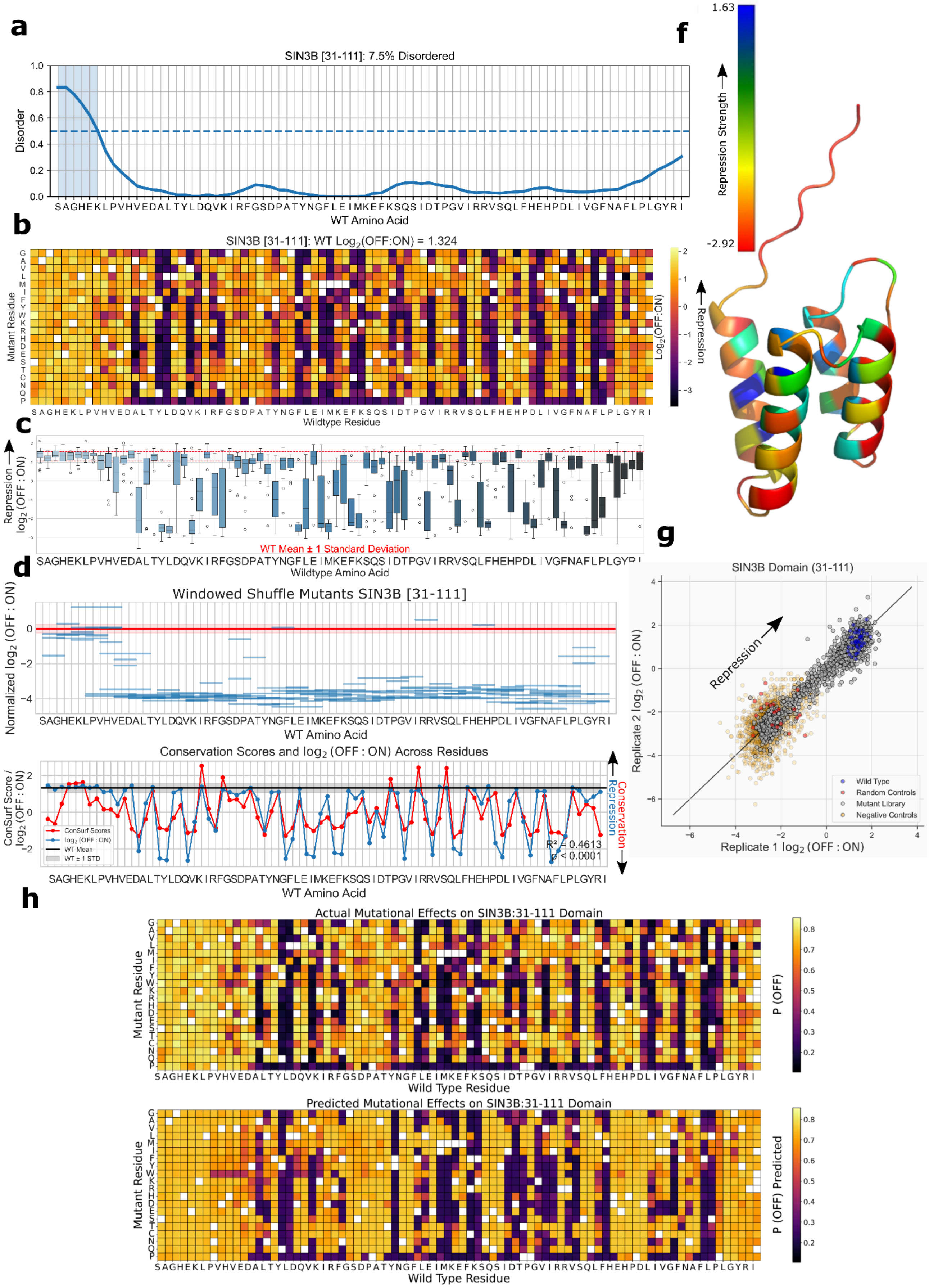

SUMO1 [18-98] Ubiquitin-2 like Rad60 SUMO-like domain

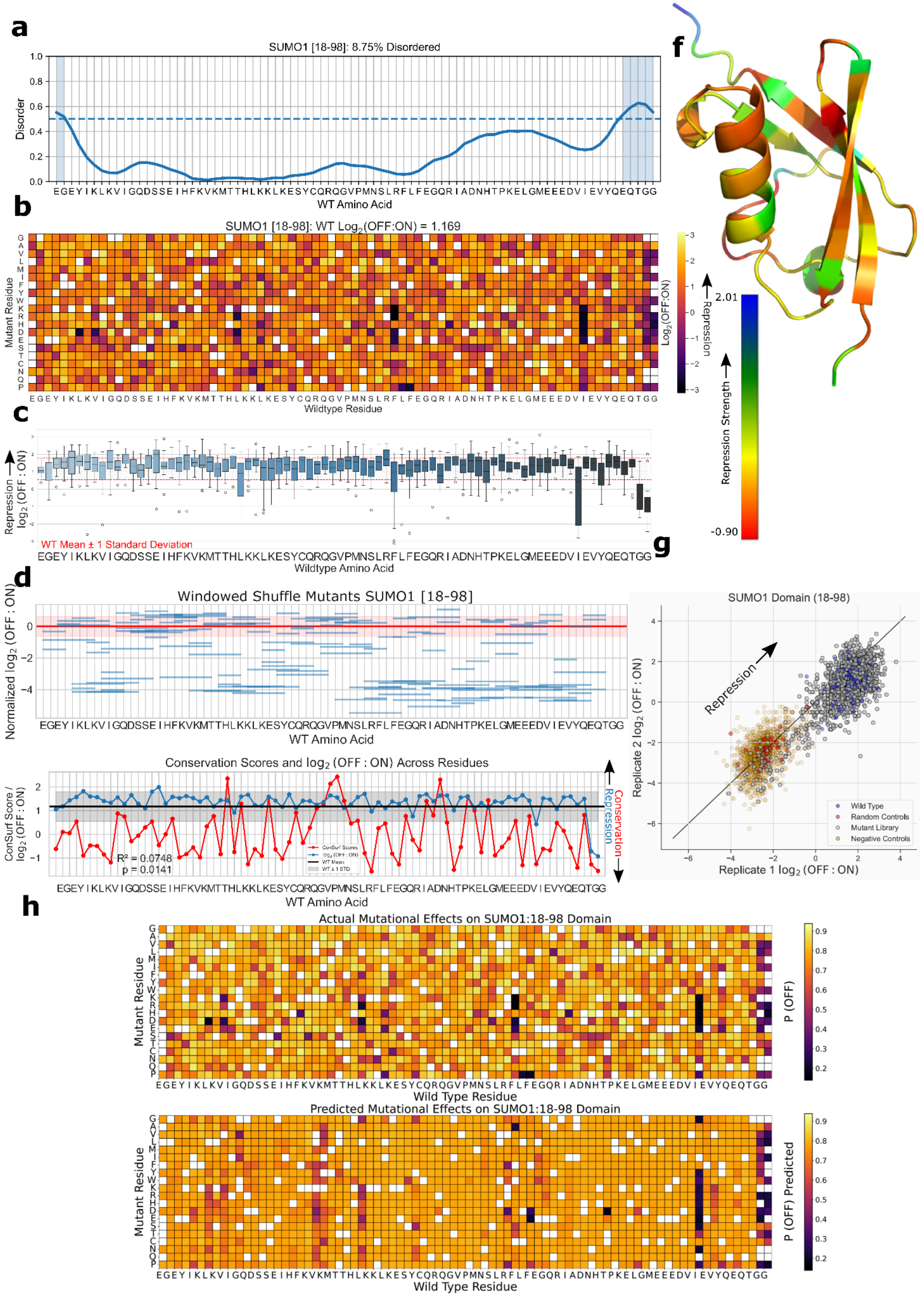

### SUMO3 [13-93] Ubiquitin-2 like Rad60 SUMO-like domain

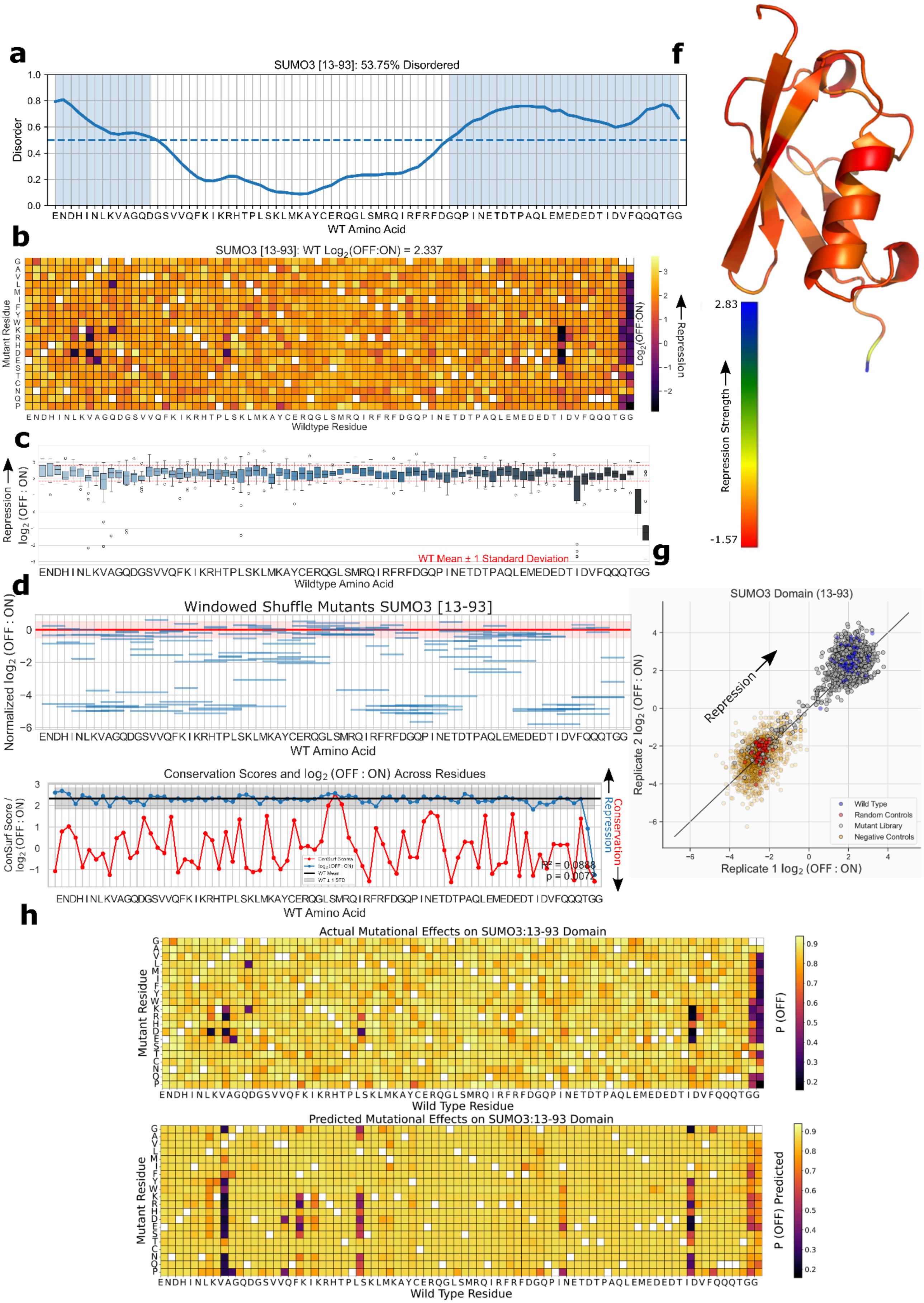

### TET2 [1920-2000] Unannotated domain

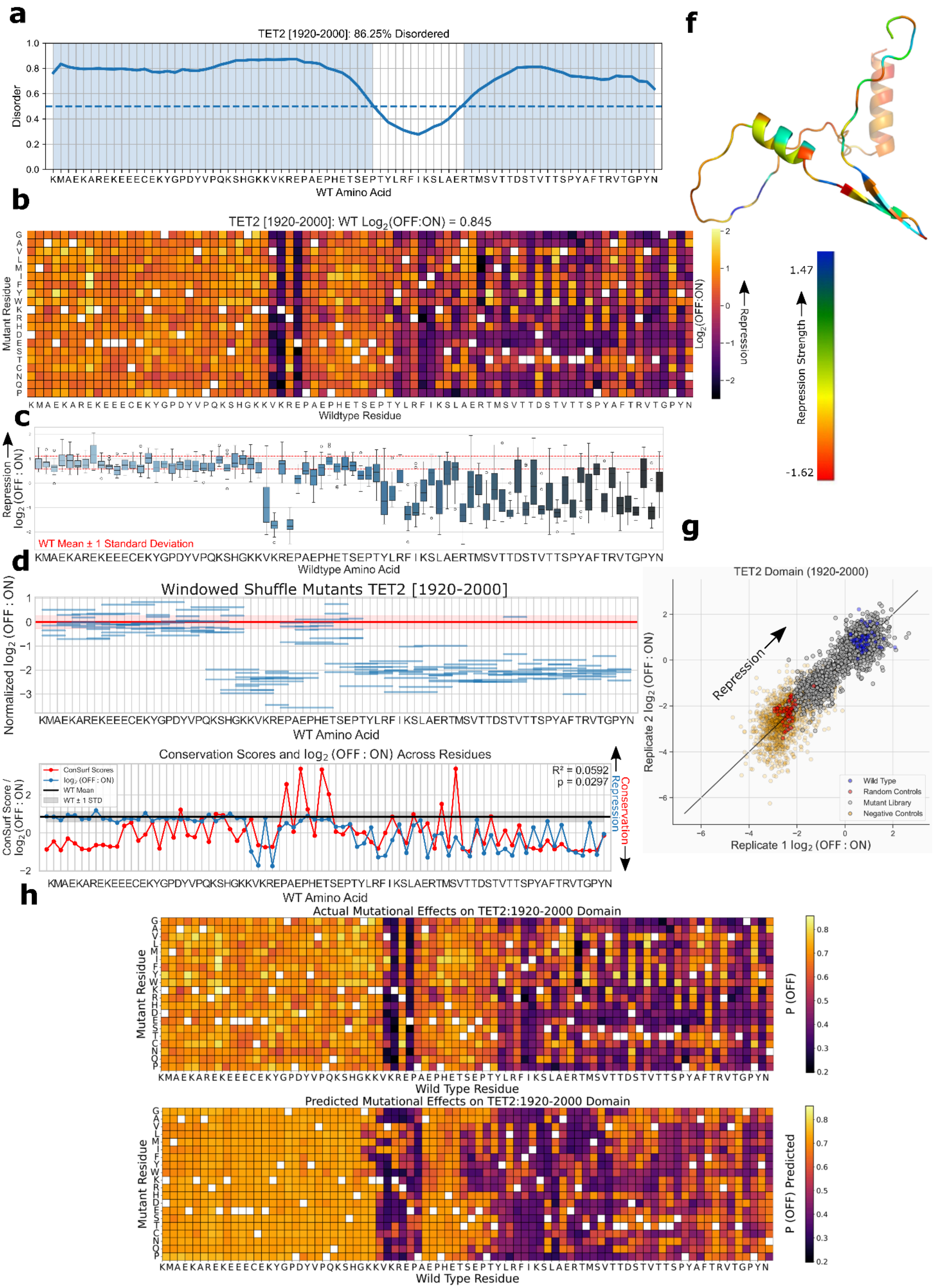

### TWST2 [53-133] Helix-loop-helix DNA binding domain

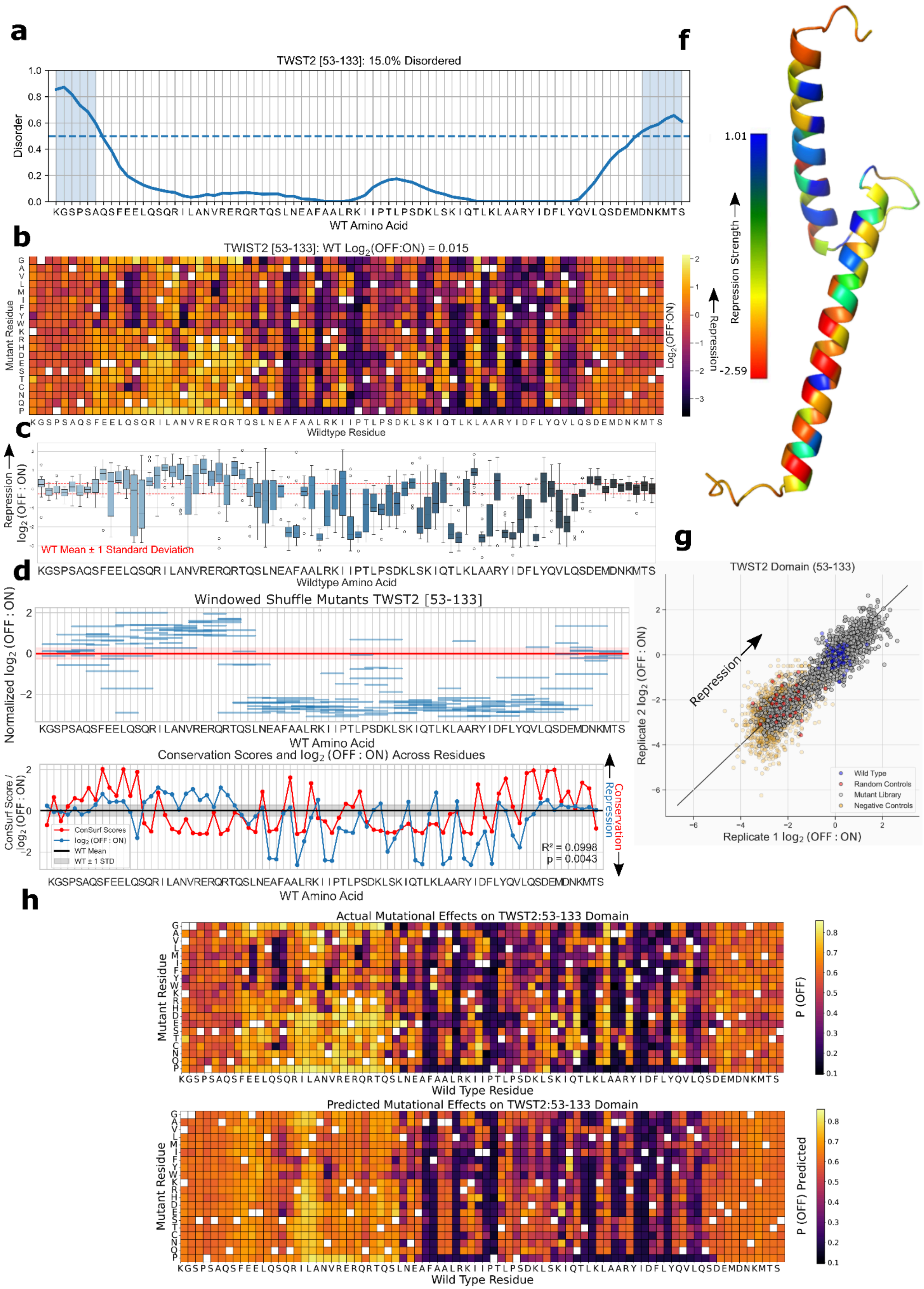

UHRF1 [321-401] PHD-finger domain

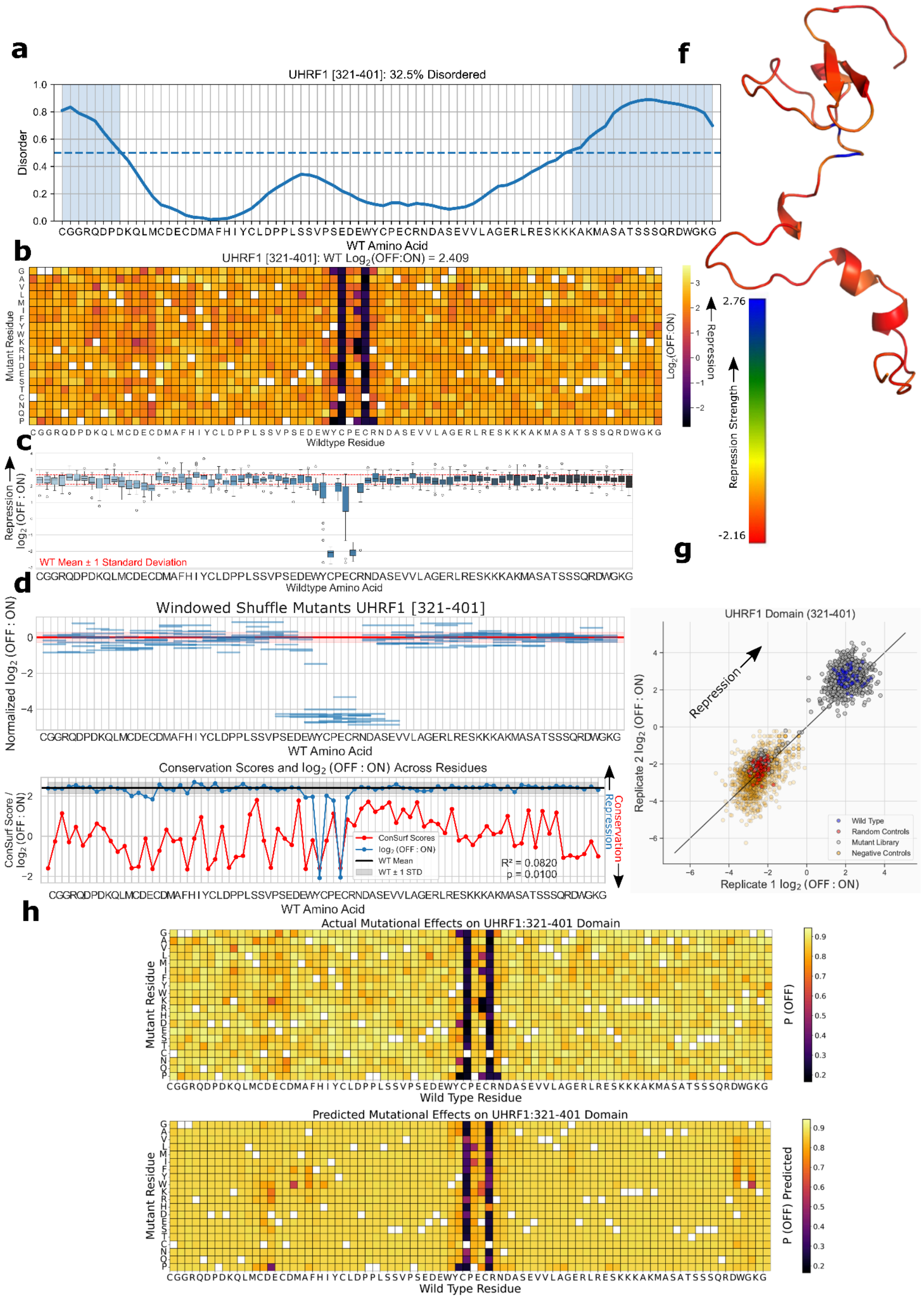

### YAF2 [79-159] Yaf2/RYBP C-terminal binding motif domain

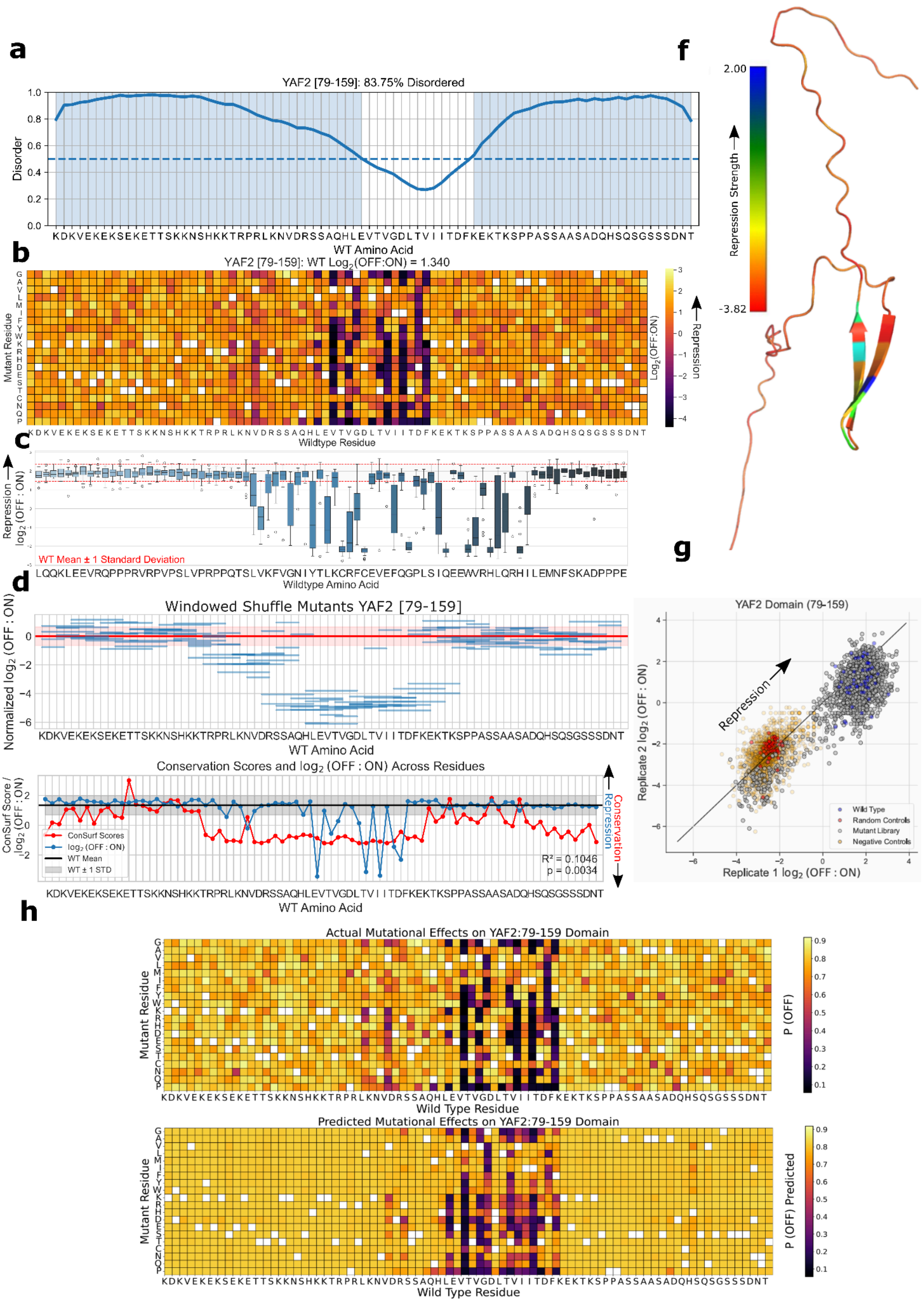

### ZNF10 [2-82] KRAB box domain

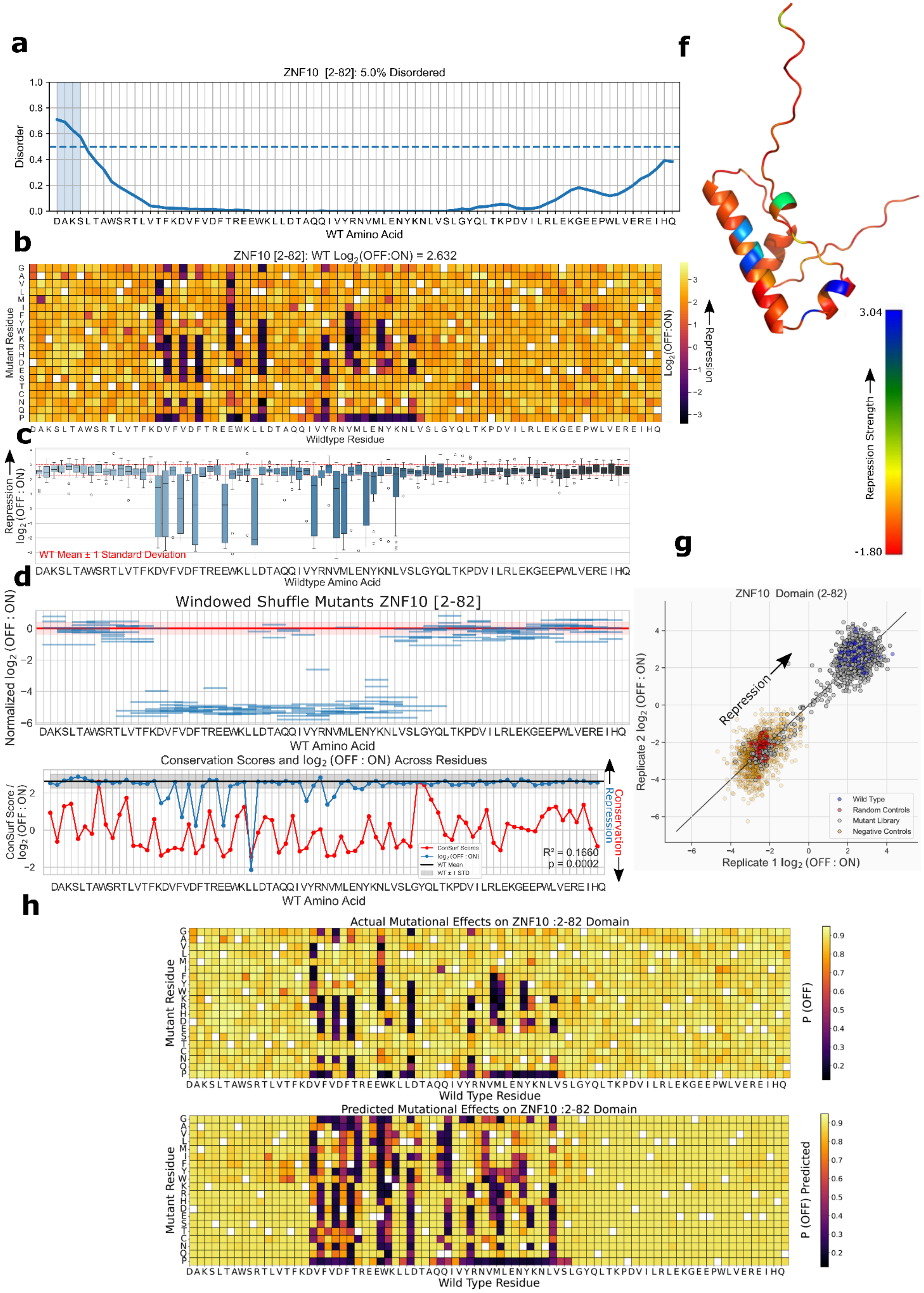
